# Histone deacetylation and cytosine methylation compartmentalize heterochromatic regions in the genome organization of *Neurospora crassa*

**DOI:** 10.1101/2023.07.03.547530

**Authors:** Ashley W. Scadden, Alayne S. Graybill, Clayton Hull-Crew, Tiffany J. Lundberg, Nickolas M. Lande, Andrew D. Klocko

**Author notes:** **Corresponding Author:** Andrew D. Klocko. **Author Contributions:** A.W.S and A.D.K. designed research; A.W.S., A.S.G., C.H-C., T.J.L., N.M.L., and A.D.K. performed research; A.W.S., A.S.G., C.H-C., T.J.L., N.M.L., and A.D.K. analyzed data; and A.W.S., A.S.G., C.H-C., T.J.L., N.M.L., and A.D.K. wrote the paper. **Competing Interest Statement:** The authors declare no conflict of interest. **Classification:** Biological Sciences, Genetics.

## Abstract

Chromosomes must correctly fold in eukaryotic nuclei for proper genome function. Eukaryotic organisms hierarchically organize their genomes, including in the fungus *Neurospora crassa*, where chromatin fiber loops compact into Topologically Associated Domain (TAD)-like structures formed by heterochromatic region aggregation. However, insufficient data exists on how histone post-translational modifications, including acetylation, affect genome organization. In Neurospora, the HCHC complex (comprised of the proteins HDA-1, CDP-2, HP1, and CHAP) deacetylates heterochromatic nucleosomes, as loss of individual HCHC members increases centromeric acetylation and alters the methylation of cytosines in DNA. Here, we assess if the HCHC complex affects genome organization by performing Hi-C in strains deleted of the *cdp-2* or *chap* genes. CDP-2 loss increases intra– and inter-chromosomal heterochromatic region interactions, while loss of CHAP decreases heterochromatic region compaction. Individual HCHC mutants exhibit different patterns of histone post-translational modifications genome-wide: without CDP-2, heterochromatic H4K16 acetylation is increased, yet smaller heterochromatic regions lose H3K9 trimethylation and gain inter-heterochromatic region interactions; CHAP loss produces minimal acetylation changes but increases heterochromatic H3K9me3 enrichment. Loss of both CDP-2 and the DIM-2 DNA methyltransferase causes extensive genome disorder, as heterochromatic-euchromatic contacts increase despite additional H3K9me3 enrichment. Our results highlight how the increased cytosine methylation in HCHC mutants ensures genome compartmentalization when heterochromatic regions become hyperacetylated without HDAC activity.

**Significance Statement:** The mechanisms driving chromosome organization in eukaryotic nuclei, including in the filamentous fungus *Neurospora crassa*, are currently unknown, but histone post-translational modifications may be involved. Histone proteins can be acetylated to form active euchromatin while histone deacetylases (HDACs) remove acetyl marks to form silent heterochromatin; these heterochromatic regions cluster, forming strong interactions, in Neurospora genome organization. Here, we show that mutants of a heterochromatin-specific HDAC, HCHC, increase heterochromatic histone acetylation genome-wide and contact probability between distant heterochromatic loci. HCHC loss also impacts cytosine methylation, and in strains lacking both the HCHC and cytosine methylation, heterochromatic regions interact more with euchromatin. Our results suggest cytosine methylation normally functions to segregate silent and active loci when heterochromatic acetylation increases.

## Introduction

Chromosomes must correctly fold in eukaryotic nuclei for proper genome function, including for transcription regulation (1–5). This genome organization includes the subnuclear compartmentalization of chromatin (2, 5) where the silent heterochromatin aggregates at the nuclear periphery to prevent RNA polymerase binding while active euchromatin is centrally localized and accessible to the transcription machinery (6–9). In euchromatin, long-range loops form between promoters and distant regulatory sequences, including enhancers or silencers, for transcriptional control (3, 10). Genome organization is frequently disrupted in human cancers, where promoters hijack regulatory sequences for improper gene expression (11–13). Given the direct connection between genome organization and function, there is a critical need to understand the factors involved in wild type (WT) chromosome folding.

Genome organization is assessed by chromosome conformation capture with high throughput sequencing (Hi-C), in which crosslinked chromatin is restriction enzyme digested and ligated to physically connect interacting DNA strands in a single DNA molecule; the updated *in situ* Hi-C protocol ligates interacting chromatin in the nucleus, thereby generating more accurate genome organization contact probability matrices (14–16). Hi-C has shown that eukaryotic genomes are hierarchically organized, where histone octamers wrap ∼146 basepairs (bp) of DNA to form the nucleosomes comprising chromatin fiber globules (“loops”) (17, 18). Globules compact into Topologically Associated Domains (TADs), where chromatin internal to the TAD is more apt to interact (15, 19–21). TADs of similar transcriptional states are compartmentalized, while individual metazoan chromosomes isolate into territories (6, 14, 15, 22–25).

Filamentous fungi, like *Neurospora crassa*, are excellent model organisms for understanding eukaryotic genome organization (16, 26–28). The predominantly haploid Neurospora genome, typical among fungi, is small (4.1 x 10^7^ bp) (29), which allows for cost-efficient genomic sequencing experiments. The Neurospora genome contains both euchromatic and heterochromatic loci, with the latter subdivided into permanently-silent constitutive heterochromatin comprised of gene-poor, repetitive, adenine-thymine (AT)-rich DNA, and temporarily-silent facultative heterochromatin covering gene-rich regions (29–32); each type is marked by specific histone post-translational modifications (PTMs). Neurospora constitutive heterochromatin is enriched with trimethylation of lysine 9 on histone H3 (H3K9me3, catalyzed by the **D**IM-5/-7/-9, **C**UL4, **D**DB1*^dim-8^* **C**omplex [DCDC]) and cytosine methylation (5^m^C), while the di-or trimethylation of lysine 27 on histone H3 (H3K27me2/3) or the ASH1-dependent di-methylation of lysine 36 on histone H3 (H3K36me2) demarcates facultative heterochromatin (30–38). Histone lysines in euchromatin are extensively acetylated by histone acetyltransferases (HATs), including on histone H3 lysine 9 (H3K9ac) or histone H4 lysine 16 (H4K16ac) (39–41). Unlike metazoans, the machinery catalyzing histone PTMs or 5^m^C in Neurospora is simplistic, non-redundant, and not required for viability, promoting the use of Neurospora for understanding chromatin function (32, 38, 42). Notably, the Neurospora genome is compacted and organized similarly to metazoan genomes, with a comparable genome size to nuclear volume ratio (16). Neurospora chromosomes (“Linkage Groups” [LG]) form TAD-like structures, also known as “Regional Globule Clusters”, in which the heterochromatic regions flanking euchromatin aggregate, forming loops that cluster several euchromatic globules together in an uninsulated manner (16, 26, 27). However, as opposed to the robust chromosome territories in metazoans (1, 22, 24, 25), fungal chromosomes form a Rabl conformation, in which centromeres bundle independently from telomere clusters at the nuclear periphery (16, 26–28, 43), as directly observed in Neurospora (26–28); this Rabl structure allows for weak chromosome territories with extensive inter-chromosomal contacts.

Chromatin compartmentalization may depend on individual histone PTMs. Since specific histone PTMs are known to define chromatin transcriptional status and accessibility (18, 44–46), histone PTMs may also be necessary for segregating euchromatin from heterochromatin in the nucleus. Recent reports detail how heterochromatic histone PTMs, or their cognate binding partners, organize eukaryotic genomes. This includes H3K9me3 and Heterochromatin Protein-1 (HP1), which compact heterochromatic regions during fungal asexual growth and facilitate pericentromere clustering during Drosophila zygote genome activation (27, 43, 47). In addition, sub-telomeric H3K27me2/3 helps telomere clusters associate with the nuclear periphery in Neurospora (26). However, little information exists about the roles of other histone PTMs, including histone acetylation.

Classically, acetylated histone tails denote open chromatin, as the euchromatic regions of fungi, fruit flies, mice, and humans are highly acetylated (33, 48–57). In contrast, heterochromatin is typically devoid of histone acetylation, as histone deacetylase (HDAC) complexes remove acetyl marks from silent loci (33, 39, 40, 58). Changes in acetylation enrichment impact genome organization, including the interactions between promoters and enhancers. Differential acetylation of lysine 27 on histone H3 (H3K27ac) across enhancers is observed during the differentiation of human epidermal cells (59) and mouse adipocyte cells (60). Histone acetylation changes also occur throughout the cell cycle, as mitotic chromosomes are globally hypoacetylated, which forms dense chromosomal structures allowing separation into daughter cells (61–65). H3K27 within promoters and enhancers is deacetylated during mitosis but rapidly re-acetylated in G1 for activating transcription and reforming TADs (66, 67). Further, the BRD4-NUT novel oncogenic protein recruits the HAT P300 to create hyperacetylated Mega-domains that aggregate into a transcriptionally activated sub-compartment (68); phase-separation might promote sub-compartment formation (69). However, it is unknown how HDAC activity affects fungal chromosome conformation.

*Neurospora crassa* encodes four class II HDACs (HDA-1 to HDA-4; class characterization based on primary structure conservation), with HDA-1 impacting heterochromatin (41, 58). HDA-1 is a component of the four-member HCHC complex, comprised of the proteins HDA-1, CDP-2 (***C***hromo***d***omain ***P***rotein-***2***), HP1, and CHAP (***C***DP-2 and ***H***DA-1 ***A***ssociated ***P***rotein) (39, 40). HDA-1 is the deacetylase catalytic subunit, HP1 binding to H3K9me3 recruits the complex to heterochromatin, CDP-2 mediates protein interactions between HP1 and HDA-1, and CHAP binds to AT-rich DNA in constitutive heterochromatin using two AT hook domains (39, 40). Both CDP-2 and CHAP recruit HDA-1 to heterochromatic genomic loci (39). Strains deleted of the genes *cdp-2*, *chap*, or *hda-1* have increased histone acetylation at select heterochromatic loci, as assessed by Chromatin Immunoprecipitation (ChIP) quantitative PCR, although genome-wide changes are unknown (40). Smaller heterochromatic regions in HCHC mutants lose 5^m^C but larger regions, including centromeres, gain 5^m^C by increasing recruitment of DIM-2, the only DNA methyltransferase encoded in Neurospora (39). Here, we show the HCHC complex impacts genome organization using Δ*cdp-2* and Δ*chap* deletion strains. CDP-2 loss increases intra– and inter-chromosomal heterochromatic region contacts, while deletion of CHAP reduces heterochromatic region compaction. Heterochromatic regions gain H4K16ac but not H3K9ac in a Δ*cdp-2* strain, while a Δ*chap* strain has little H4K16ac change. Smaller heterochromatic regions lose H3K9me3 in a Δ*cdp-2* strain, causing contact promiscuity between silent loci. A Δ*cdp-2*;Δ*dim-2* double mutant devoid of 5^m^C and HCHC activity gains H3K9me3 but loses heterochromatin – euchromatin compartmentalization, suggesting global genome disorder. We conclude that histone deacetylation is required for Neurospora genome organization, but if histones become hyperacetylated, 5^m^C helps maintain segregation of heterochromatin from euchromatin.

## Results

### Genome organization changes in strains deleted of either HCHC component CDP-2 or CHAP

The genome of the wild type (WT) *Neurospora crassa* strain N150 (74-OR23-IVA [Fungal Genetic Stock Center #2489]) is organized by heterochromatin aggregates, which form TAD-like structures (Figure 1A) (16, 26–28). Heterochromatic regions in Neurospora depend on the HDAC HCHC for deacetylation of histone tails, transcriptional silencing, and a normal 5^m^C distribution (40). To understand if the HCHC complex also organizes the Neurospora genome, we examined strains independently deleted of the genes encoding CDP-2 (Δ*cdp-2*) or CHAP (Δ*chap*) by *in situ* Hi-C. We chose CDP-2 and CHAP given the unique domains predicted by their primary protein structures and how their actions are epistatic to HDA-1 (39, 40), hypothesizing that we could observe more nuanced differences in genome organization upon loss of HCHC activity to clarify the individual roles of CDP-2 and CHAP, instead of only assessing the impact of HDA-1 catalytic activity loss. We independently performed chromatin-specific *in situ* Hi-C (16) with *Dpn*II (recognition sequence åGATC) to assess euchromatic contacts, or *Mse*I (TåTAA) to examine heterochromatic interactions, on Δ*cdp-2* and Δ*chap* strains. All Hi-C experiments reported in this manuscript are generated from populations of actively growing cells containing asynchronous nuclei, most of which are typically in interphase stages of the cell cycle with less than 5% mitotic nuclei (70, 71). Our replicate *in situ* Hi-C libraries are highly reproducible (Figures S1, S2), allowing similar restriction enzyme datasets to be merged. This produced high quality *Dpn*II Δ*cdp-2* (∼35.4 million [M] valid read pairs) and Δ*chap* (∼8.7M valid read pairs), as well as *Mse*I Δ*cdp-2* (∼13.9M valid read pairs) and Δ*chap* (∼4.8M valid read pairs) datasets; total and valid read counts are provided in Supplemental Table S1. The merged *Dpn*II datasets presented are Knight-Ruiz (KR) corrected to limit experimental bias (22, 72, 73), while *Mse*I datasets show raw read counts, since KR correction improperly removes heterochromatin signal upon bin averaging, possibly due to the discrepancy of *Mse*I sites between AT-rich heterochromatin and GC-rich euchromatin (16).

**Figure 1.**
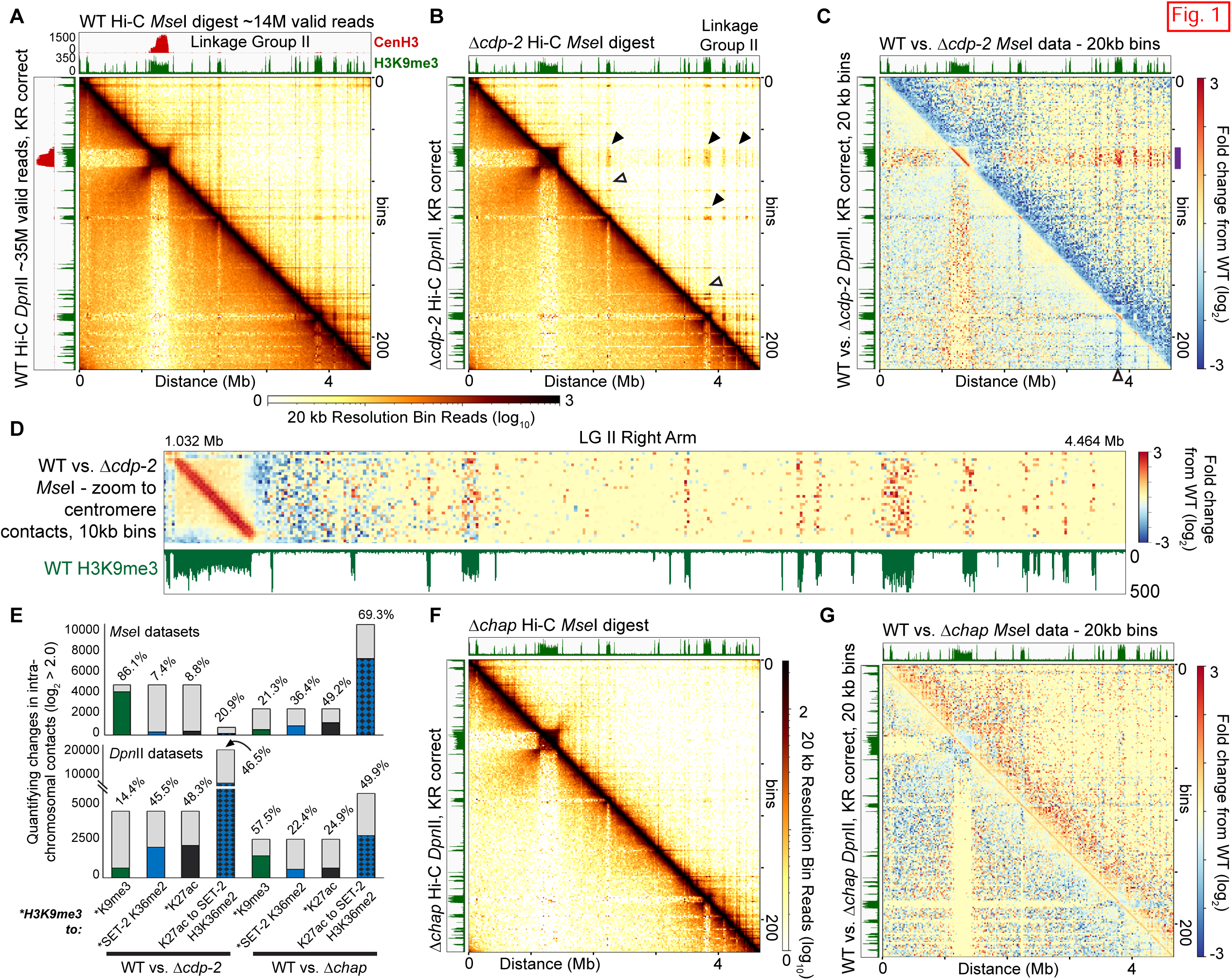
Genome organization changes across a single chromosome in Neurospora crassa Δcdp-2 and Δchap strains. (A-B) Heatmap of contact probabilities of 20 kb bins across LG II of (A) wild type (WT) *Neurospora crassa* (16) or (B) Δ*cdp-2* strains. Raw read counts of heterochromatin-specific *Mse*I Hi-C (above diagonal) and Knight Ruiz corrected (72, 73) read counts of euchromatin-specific *Dpn*II Hi-C (below diagonal) are presented in heatmaps here and throughout this work unless otherwise indicated. Bin numbers (vertical axis) and genomic distance, in Megabases (Mb), (horizontal axis) are indicated. The scale bar shows the log_10_ numbers of valid Hi-C contacts per bin. Integrative Genomics Viewer (IGV) (74) images of WT CenH3 (red; centromeres) and/or H3K9me3 (green; heterochromatic regions) ChIP-seq tracks are shown above and left. Solid arrowheads in B. show increased inter-heterochromatic contacts; open arrowheads show decreased heterochromatic-euchromatic contacts. (C) The fold change in contact strength (log_2_ scale) in a Δ*cdp-2* strain relative to a normalized WT strain of *Mse*I *in situ* Hi-C (above diagonal) or *Dpn*II *in situ* Hi-C (below diagonal) datasets; purple line indicates the region in D. The open arrowhead shows a heterochromatic region gaining internal contacts at the expense of euchromatic contacts. (D) Heatmap of contact probability changes between heterochromatic regions (green) and the LG II centromere in Δ*cdp-2 vs.* WT comparison heatmaps at 10 kb resolution. (E) Quantification of strongly increased (log2 ≥ 2.0), intra-chromosomal contacts in Δ*cdp-2* or Δ*chap* datasets, relative to normalized WT datasets, between bins enriched for select histone PTMs (see Supplemental Materials and Methods). For these graphs, each bar shows the total number of contacts that are strongly (log2 ≥ 2.0) increased. Numbers of contacts originating from bins enriched with the first histone PTM and end in a bin enriched for the second histone PTM are indicated by non-gray colors, while the gray portion of the bar shows contacts originating from the first enriched histone PTM bin but terminating in a non-enriched bin. The percentage of total contacts enriched for both histone PTMs is indicated above. (F) Contact probability heatmap of a Δ*chap* dataset, as in A. (G) Heatmap of contact probability changes between Δ*chap* and WT datasets, as in C.

The *Dpn*II and *Mse*I Δ*cdp-2* contact probability heatmaps of a single chromosome (Figures 1B) look comparable to WT *Dpn*II or *Mse*I Hi-C datasets with nearly identical valid read numbers (Figure 1A), but upon closer examination, the Δ*cdp-2* dataset has increased interactions between distant heterochromatic regions. Specifically, the LG II centromere forms prominent interactions with heterochromatic regions up to three Megabases (Mb) distant in the heterochromatin specific *Mse*I heatmap (Figure 1B, black arrowheads), while heterochromatic regions form fewer contacts with nearby euchromatin (Figure 1B, open arrowheads). These effects are seen across all seven Neurospora chromosomes (Figures S3). To highlight changes in a Δ*cdp-2* strain, we directly compared the WT and Δ*cdp-2 Dpn*II or *Mse*I Hi-C contact probability matrices. In comparison heatmaps, the strong increases between centromeres and distant heterochromatic regions in a Δ*cdp-2* strain are among the most prominent across LG II (Figure 1C) and all seven Neurospora LGs (Figure S4). At a high resolution, the Δ*cdp-2* centromeric chromatin shows increased contacts, as evident by the LG II centromere having a strong red diagonal indicative of increased contacts between centromeric nucleosomes, in addition to having more intense interactions with distant H3K9me3-marked constitutive heterochromatic regions (Figure 1D). In fact, each heterochromatic region in Δ*cdp-2 Mse*I datasets has increased local interactions on the diagonal that depletes heterochromatic-euchromatic contacts (Figures 1B-C, S4). Quantification of strongly changed contacts between WT and Δ*cdp-2 Mse*I datasets showed that most H3K9me3 enriched bins had stronger (log_2_ ≥ 2.0) interactions with another H3K9me3 enriched bin; few H3K9me3 enriched bins had strong interaction changes with bins enriched for euchromatic marks in *Mse*I datasets (Figure 1E). The contact paucity in off diagonal euchromatic interactions in a Δ*cdp-2 Mse*I dataset, relative to WT (Figure 1C, above diagonal), could indicate chromatin in TAD-like structures interact less or is due to fewer euchromatic *Mse*I sites for Hi-C ligation (16) coupled with increased heterochromatic contacts (Figure 1C). As evident in *Dpn*II datasets, interactions of distant euchromatic regions are globally decreased, including between the left and right chromosome arms and between distant TAD-like structures (Figure 1C). However, individual bins quantitatively have strongly changed interactions when comparing WT and Δ*cdp-2 Dpn*II datasets, as more bins enriched for two euchromatic histone PTMs, H3K27ac and SET-2-specific H3K36me2 (33), were increased, although some H3K9me3 marked bins interacted more with euchromatin (Figure 1E). All told, CDP-2 loss increases constitutive heterochromatin interactions which alters the normal conformation of Neurospora chromosomes.

Next, we examined the genome organization of a Δ*chap* strain by *Dpn*II and *Mse*I Hi-C. The Δ*chap* data has fewer valid reads, with ∼four-fold less valid reads in the Δ*chap Dpn*II dataset and ∼three-fold less valid reads in the Δ*chap Mse*I dataset, relative to the Δ*cdp-2* datasets (Supplemental Table S1). However, sufficient data exists in Δ*chap* datasets to allow comparison with normalized WT datasets containing similar valid read numbers, so the impact of CHAP loss can be assessed. Examination of single chromosomes show how the Δ*chap Dpn*II and *Mse*I Hi-C datasets appear devoid of most strong long-range contacts (Figures 1F, S5). In each chromosome, more distant contacts are rarer upon CHAP loss; an enhanced image of the right arm of LG II highlights that regional interactions still form in both *Dpn*II and *Mse*I datasets (Figure S6). When comparing WT and Δ*chap* datasets, heterochromatic regions on all LGs exhibit reduced internal compaction, and fewer contacts between heterochromatic regions are observed (Figures 1G; S6B, arrowheads; S7). Also apparent are the decreased interactions between the TAD-like Regional Globule Clusters (16) in the Δ*chap Dpn*II data, which may stem from the slight increase in on-diagonal, local euchromatic interactions (Figures 1G, S6-S7). Quantification of strongly increased contacts in Δ*chap Dpn*II or *Mse*I datasets relative to WT shows that interactions between bins enriched for the euchromatic marks H3K27ac and SET-2-specific H3K36me2 (33) are the most frequently increased when CHAP is lost (Figure 1E).

Similarly, loss of CDP-2 and CHAP impact the inter-chromosomal clustering of heterochromatic regions across the entire Neurospora genome. As shown in raw *Mse*I and KR-corrected *Dpn*II contact probability heatmaps of a Δ*cdp-2* strain, or when these Δ*cdp-2* data are compared to normalized WT contact matrices, Δ*cdp-2* centromeres gain interactions with the interspersed heterochromatic regions across all seven LGs, concomitantly weakening the inter-chromosomal centromere interactions and the contacts between centromere-proximal euchromatin (Figures S8-S9), although these reductions may result if one dataset (Δ*cdp-2*) has more heterochromatin contacts when comparing datasets with equal valid read pairs. The Δ*cdp-2 Dpn*II Hi-C data have stronger inter-chromosomal arm contacts (Figure S9); similar changes are observed in KR-corrected *Mse*I data and raw *Dpn*II data when comparing WT and Δ*cdp-2* contact matrices (Figure S10A). Examination of the interactions between two chromosomes, LG II and LG III, in Δ*cdp-2 Mse*I Hi-C contact heatmaps show the gain of centromeric and interspersed heterochromatic region contacts and the concomitant reduction of contacts between silent and active chromatin (Figure S8A). Plotting the inter-chromosomal interaction changes of WT and Δ*cdp-2 Mse*I datasets between LG II and LG III highlight how distant heterochromatic regions between chromosomes are more apt to interact with the centromere bundle (Figure 2A). Decreased contacts include the inter-centromeric and pericentromeric euchromatin (Figure 2A). CHAP loss reduces heterochromatic interactions across the Neurospora genome, with few inter-chromosomal heterochromatic contacts observed in Δ*chap Mse*I contact matrices (Figure S11), and prominent reductions in centromeric contacts that are independent of matrix correction when WT and Δ*chap Mse*I data is compared (Figures S9B, S10B). A plot of the interactions between LG II and LG III in Δ*chap* shows the paucity of inter-chromosomal constitutive heterochromatic contacts (Figure S11A), which is highlighted when comparing WT and Δ*chap Mse*I Hi-C data (Figure 2B). The decrease in heterochromatin bundling compromises the inter-chromosomal arm interactions characteristic of the Rabl conformation (28), as reduced euchromatin interactions across the left and right chromosomal arms are evident in a WT and Δ*chap Dpn*II Hi-C comparison (Figure S10B). Quantification of the strongest inter-chromosomal interaction gains between histone PTM-enriched bins in a Δ*cdp-2 Mse*I dataset, relative to WT, shows 81.5% of H3K9me3 enriched bins gain interactions, but few H3K9me3-enriched bins gain strong contacts with bins enriched with euchromatic marks, such as H3K27ac (Figure 2C, left). The Δ*cdp-2 Dpn*II Hi-C data show changes in strong interactions between bins enriched with H3K9me3 and euchromatic histone PTMs, such as H3K27ac and SET-2 specific H3K36me2, or between two euchromatic enriched bins (Figure 2C, right). In contrast, Δ*chap Mse*I and *Dpn*II Hi-C datasets have minimal gains in interactions relative to WT, independent of histone PTMs, consistent with the loss of CHAP reducing heterochromatic interactions (Figure 2C). To highlight the differential effects of CDP-2 and CHAP loss on genome organization, we compared equal numbers of valid reads to highlight interaction changes in a Δ*chap* strain relative to a Δ*cdp-2* strain. Here, the decreases in heterochromatin compaction, inter-heterochromatic region contacts, and contacts between chromosomal arms, with the concomitant increase in local euchromatic interactions that occur upon loss of CHAP, relative to the CDP-2 changes, is evident (Figure S12). All told, our Δ*cdp-2* and Δ*chap* Hi-C data reveal the differential effects on genome organization by individual members of a histone deacetylase complex.

**Figure 2.**
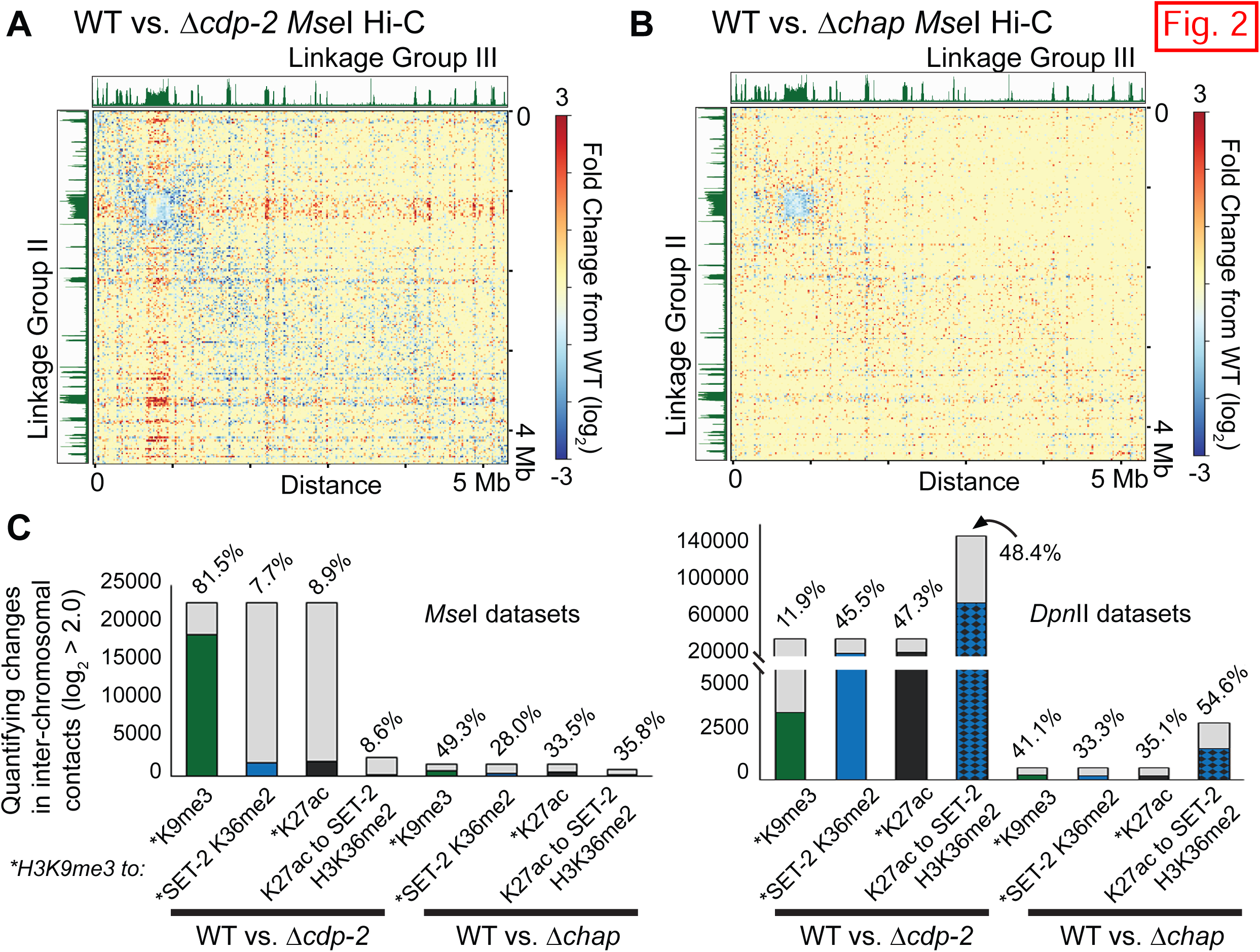
Δcdp-2 and Δchap strains alter the inter-chromosomal genome organization. (A-B) The log_2_ contact strength change of (A) Δ*cdp-2* or (B) Δ*chap* datasets, relative to normalized WT dataset between LG II and LG III within *Mse*I *in situ* Hi-C datasets, displayed similarly as Figure 1C. (C) Quantification of strongly increased, inter-chromosomal contacts in Δ*cdp-2* or Δ*chap* datasets relative to a WT dataset, as in Figure 1E.

### Histone acetylation changes in single deletion strains of the HCHC components CDP-2 and CHAP

To understand how the genome organization changes caused by deficient HCHC function affects genome function, we wished to correlate changes in Hi-C contact probability to any genome-wide changes in histone acetylation in Δ*cdp-2* and Δ*chap* strains. However, no Chromatin Immunoprecipitation-sequencing (ChIP-seq) datasets of HCHC mutants were publicly available, as recent publications only used ChIP qPCR to conclude that heterochromatic loci in HCHC deficient strains are hyperacetylated (40). Therefore, we performed two ChIP-seq replicates of histone acetylation on two lysine residues, K9 on histone H3 (H3K9ac) and K16 on histone H4 (H4K16ac), in Δ*cdp-2* and WT strains; we also performed H4K16ac ChIP-seq on a Δ*chap* strain. Both H3K9ac and H4K16ac were proposed as targets for deacetylation by the HCHC complex (40). Our H3K9ac and H4K16ac ChIP-seq replicates are qualitatively reproducible in each strain (Figure S13, S14), so we merged H3K9ac or H4K16ac replicates for Integrative Genome Viewer (IGV) display (74); all ChIP-seq data reported in this manuscript are normalized by Reads Per Kilobase per Million reads (RPKM) to ensure proper comparison between experiments, although the use of spike-in DNA (75–77) might have detected additional enrichment changes. Consistent with published ChIP qPCR, CDP-2 loss caused an increase in H4K16ac enrichment only in AT-rich heterochromatic regions but minimally effected H3K9ac levels across Neurospora chromosomes (Figure 3A, top). Closer examination of the LG II right arm highlights how each H3K9me3-marked heterochromatic region gains H4K16ac, but not H3K9ac, in a Δ*cdp-2* strain (Figure 3A, bottom, arrowheads), while euchromatic acetylation of either histone lysine is relatively unchanged (Figure 3A). In contrast, CHAP loss had little effect on H4K16ac enrichment within constitutive heterochromatic or genic regions (Figure S15A).

**Figure 3.**
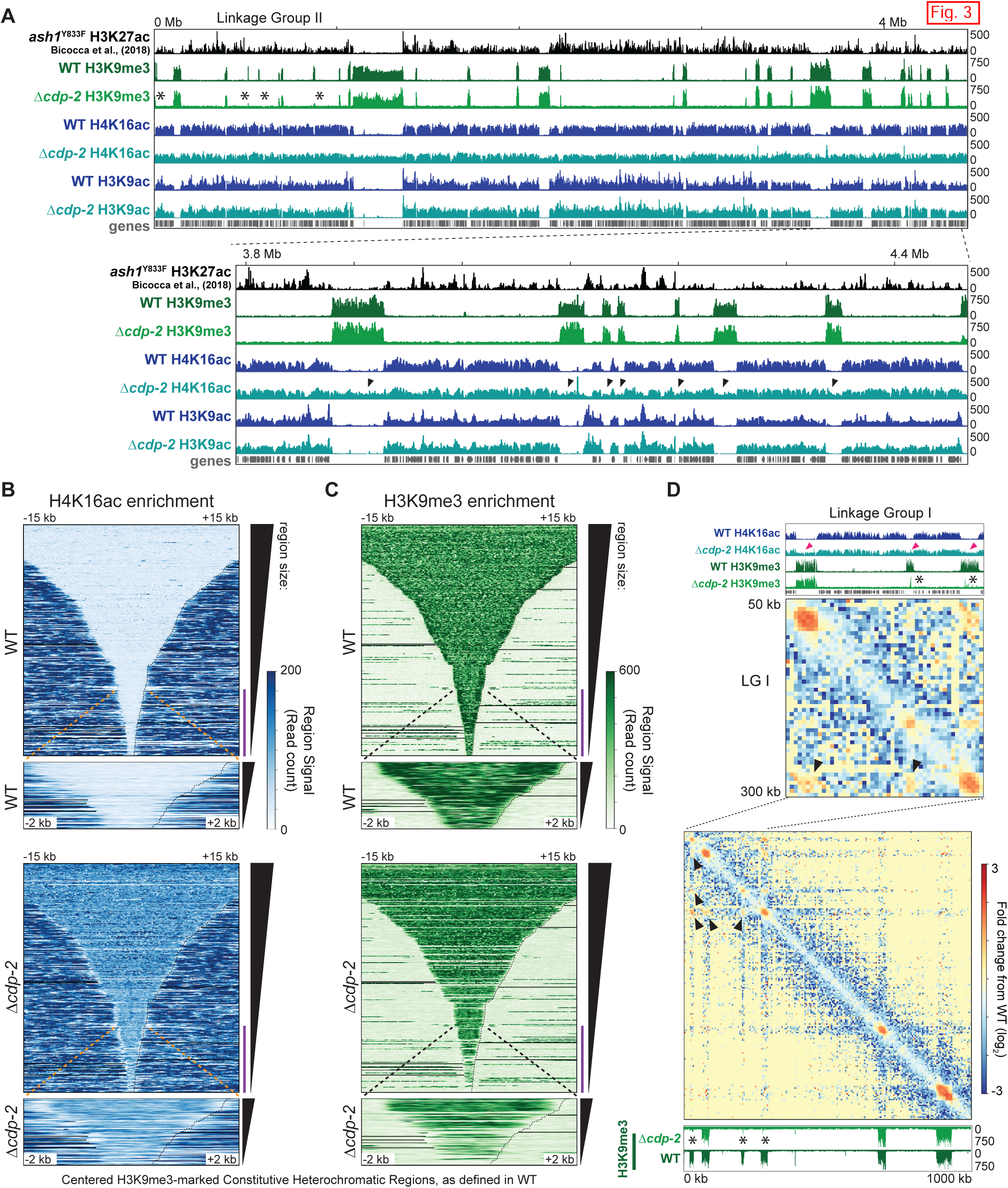
CDP-2 deletion increases heterochromatic H4K16ac while smaller heterochromatic regions lose H3K9me3. (A) IGV images of H3K9me3, H4K16ac, and H3K9ac ChIP-seq enrichment tracks from WT and Δ*cdp-2* strains across LG II (top) and the LG II right arm (bottom). SET-2-specific H3K27ac (from a Δ*ash1*^Y833F^ strain (33)) shows euchromatin. Arrowheads highlight heterochromatic regions gaining H4K16ac enrichment, while asterisks show AT-rich regions losing H3K9me3 in Δ*cdp-2*. (B-C) Average enrichment heatmaps of the (B) H4K16ac or (C) H3K9me3 enrichment in WT and Δ*cdp-2* strains centered over WT-defined constitutive heterochromatic regions (16), extending +/− 15 kb from heterochromatic region centers (top) or +/− 2 kb of the smallest regions (bottom; indicated by the purple line in +/-15 kb plots). (D) Heatmaps of contact probability change between Δ*cdp-2* and WT *Mse*I datasets of a small region of LG I. H3K9me3 and/or H4K16ac ChIP-seq track images shown with each heatmap. Asterisks indicate regions losing H3K9me3, pink arrowheads show regions gaining H4K16ac in a Δ*cdp-2* strain, and black arrowheads highlight gains in inter-heterochromatic region contacts in a Δ*cdp-2* strain.

These acetylation changes are highlighted by average enrichment plots and heatmaps of normalized H4K16ac and H3K9ac datasets across H3K9me3-enriched heterochromatic regions and genes. By aligning heterochromatic region centers and ordering these silent regions by length, we observe that essentially all AT-rich heterochromatic regions, independent of size, have substantial increases in H4K16ac enrichment in a Δ*cdp-2* strain relative to a WT strain (Figure 3B). Similar gains in H4K16ac in a Δ*cdp-2* strain are seen when each heterochromatic region is scaled to 10 kb and ordered by region length (Figure S16A), while the H3K9ac enrichment in Δ*cdp-2* heterochromatin may be slightly lower than that of WT (Figure S17A). In contrast, this approach shows minimal gains of heterochromatic H4K16ac in a Δ*chap* strain (Figure S15B). When assessing H4K16ac and H3K9ac over genes (scaled to 2.5 kb), a Δ*cdp-2* strain has minimal H4K16ac reduction, while H3K9ac slightly shifts towards the 3’ end of genes, relative to WT (Figures S16B and S17B); a Δ*chap* strain has a nominal decrease in genic H4K16ac (Figure S15C). To visualize the H4K16ac and H3K9ac deposition changes, we plotted the normalized, averaged signal of each region with boxplots. The heterochromatic regions in Δ*cdp-2* have robust increases in H4K16ac signal relative to WT (Figure S16C), while H3K9ac enrichment is generally decreased (Figure S17C), although some regions in either dataset go against these trends. Considering genic acetylation, a Δ*cdp-2* strain has a small but statistically significant H4K16ac decrease (Figure S16D) and H3K9ac increase (Figure S17D). Together, HCHC loss differentially effects enrichment of histone acetylation across the Neurospora genome.

### Changes in H3K9me3-deposition and heterochromatin bundling in Δcdp-2 and Δchap strains

Given the gain of constitutive heterochromatic acetylation in HCHC mutants, we examined H3K9me3 enrichment by ChIP-seq to assess other possible histone PTM differences in permanently silent loci in Δ*cdp-2* and Δ*chap* strains. The H3K9me3 ChIP-seq replicates of each strain are reproducible (Figure S18), allowing us to merge the replicates from each strain. Most H3K9me3 enrichment in WT and Δ*cdp-2* strains is similar (Figure 3A). However, several moderate to small-sized heterochromatic regions normally enriched with H3K9me3 in WT are nearly devoid of H3K9me3 in a Δ*cdp-2* strain (Figures 3A, S19A asterisks), while nearby heterochromatic regions still retain H3K9me3; regions that lose H3K9me3 do not gain H3K9ac (Figure S19A). Zooming into two affected regions, we observe background levels of H3K9me3 across the entire locus, indicating read mapping is unaffected and arguing against underlying DNA changes causing H3K9me3 loss (Figure S19A). In contrast, CHAP removal causes subtle H3K9me3 gains in most heterochromatic regions, relative to WT, even with RPKM normalization, (Figure S15A). Average enrichment plots and heatmaps, the latter ordered by region length, show most heterochromatic regions, but not genes, gain H3K9me3 enrichment in a Δ*chap* strain (Figure S15D-E). Conversely, while many larger heterochromatic regions have unchanged H3K9me3 with CDP-2 loss, there is a tendency of smaller AT-rich loci to have a near-complete loss of H3K9me3 (Figures 3C, S19B). Boxplots of average signal across scaled, WT-defined heterochromatic regions highlight the lower H3K9me3 signal across all AT-rich regions in a Δ*cdp-2* strain (Figure S19C). Interestingly, borders of heterochromatic regions often have the most pronounced H4K16ac increase and H3K9me3 depletion upon CDP-2 loss (Figure S20A-B). To highlight if the same regions that gain H4K16ac also lose H3K9me3, we k-means clustered the signal of these two marks across heterochromatic regions. K-means cluster heatmaps extending +/-15 kb or +/-2 kb show how individual AT-rich regions gain H4K16ac and lose H3K9me3 in a Δ*cdp-2* strain (Figure S20C-D).

We hypothesized that the H3K9me3 loss and H4K16ac gain might affect the genome organization of these formerly heterochromatic regions. To this end, we plotted the Hi-C contact probability changes between WT and Δ*cdp-2* datasets across two regions on LG I and LG III, each of which encompass AT-rich regions losing H3K9me3 in a Δ*cdp-2* strain (Figures 3D, S21). In both instances, H3K9me3-deficient regions strongly gain contacts with other AT rich regions retaining this silencing mark. For example, on LG I, the loss of H3K9me3 from (and concomitant gain of H4K16ac in) multiple AT-rich loci caused gains in inter-region contacts, while several nearby regions retaining H3K9me3 are unaffected (Figure 3D, closed arrowheads). On LG III, the contacts between two nearby AT-rich genomic regions are increased when one region loses H3K9me3, while two similarly sized heterochromatic regions have few long-range interaction changes (Figure S21, compare closed and open arrowheads). All told, the loss of CDP-2 impacts histone PTM enrichment, which causes increases in contact probability between AT-rich loci in heterochromatin bundles.

### Changes in histone PTMs and genome organization in a Δcdp-2;Δdim-2 double mutant

To this point, we have shown that the heterochromatic regions of strains individually deleted of HCHC complex members gain acetylation, which alters H3K9me3 deposition and heterochromatin bundling. However, CDP-2 or CHAP removal also alters cytosine methylation levels: centromeres and other large interspersed heterochromatic regions gain 5^m^C, while smaller AT-rich loci lose 5^m^C (39, 40). Hypothetically, the extra heterochromatic 5^m^C in an HCHC mutant may confer additional properties to AT-rich loci, meaning 5^m^C gains should be considered when assessing genome organization. In support, a Δ*cdp-2*;Δ*dim-2* strain devoid of the only DNA methyltransferase in Neurospora, has a compromised growth rate relative to single HCHC mutants (40).

To understand if an increase in 5^m^C alters histone PTM deposition following HCHC complex loss, we examined levels of H4K16ac and H3K9me3 in a Δ*cdp-2*;Δ*dim-2* double mutant strain. Our ChIP-seq replicate experiments are qualitatively comparable (Figures S14, S18), allowing us to merge identical, RPKM normalized, ChIP-seq histone PTM replicates. The distribution of H4K16ac in a Δ*cdp-2*;Δ*dim-2* strain mirrors that of Δ*cdp-2*, as both mutant strains gain H4K16ac across all heterochromatic regions, regardless of size, while genic acetylation is minimally altered (Figure 4A-B, S22A). The average heterochromatic H4K16ac signal in a Δ*cdp-2*;Δ*dim-2* strain is slightly reduced compared to Δ*cdp-2* but strongly increased relative to WT H4K16ac levels (Figure S22B). Interestingly, most AT-rich regions in a Δ*cdp-2*;Δ*dim-2* strain strongly gain H3K9me3 when compared to WT and Δ*cdp-2* strains (Figures 4A), as evident in heatmaps of the H3K9me3 enrichment across sorted regions (Figures 4C, S22C), or boxplots of the average H3K9me3 deposition across scaled AT-rich regions (Figure S22D). However, the same, typically smaller, AT-rich regions completely lack H3K9me3 in both the Δ*cdp-2* and double mutant strains (Figures 4A, asterisks; 4C). Thus, histone acetylation gains coupled with 5^m^C loss at AT-rich loci signals for increased H3K9me3 by the DCDC (38).

**Figure 4.**
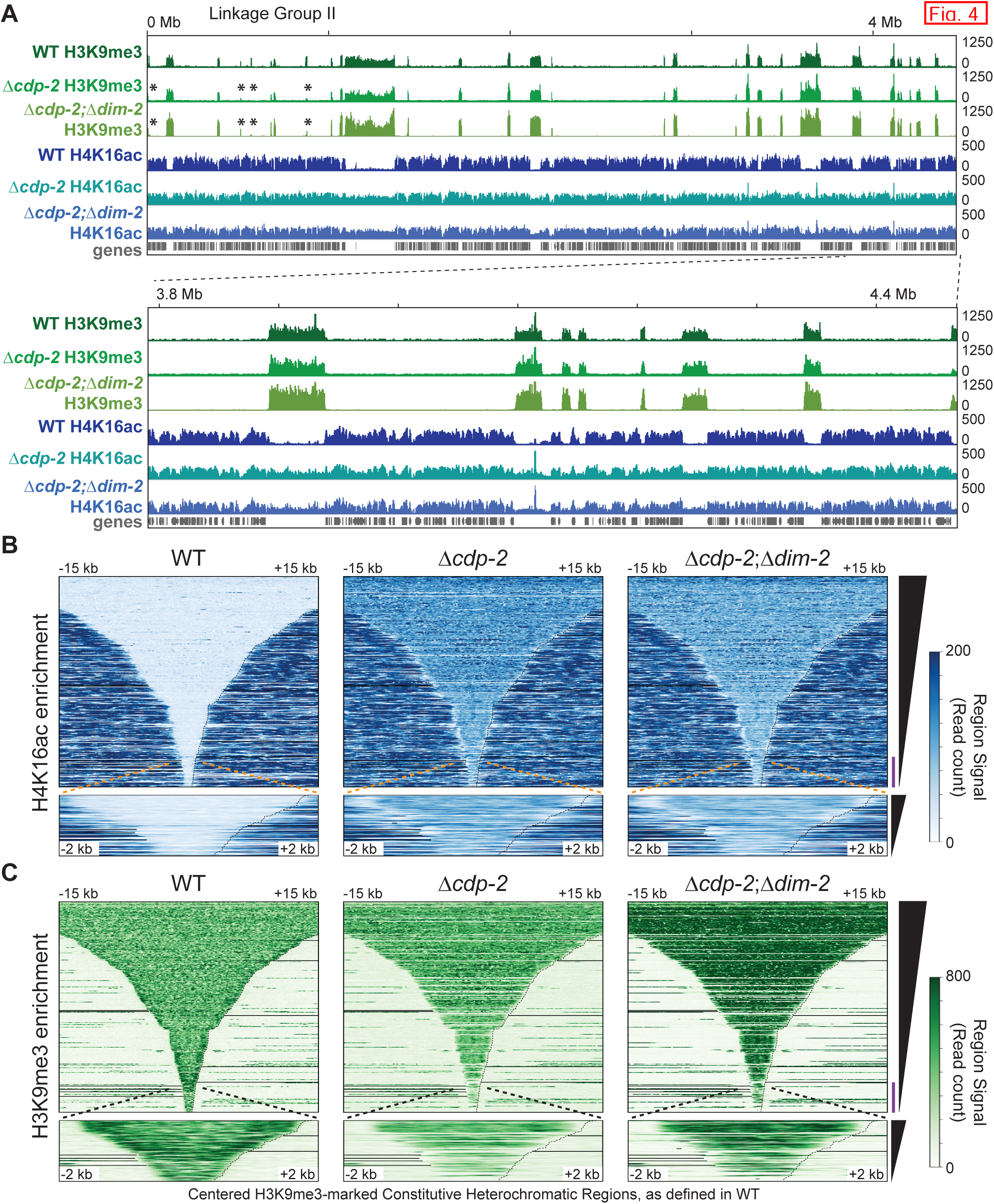
Constitutive heterochromatic regions gain H4K16ac and H3K9me3 enrichment upon CDP-2 and DIM-2 loss. (A) IGV images of H3K9me3 and H4K16ac ChIP-seq enrichment tracks of WT, Δ*cdp-2*, and Δ*cdp-2;*Δ*dim-2* strains, as in Figure 3A. Asterisks indicate AT-rich regions losing H3K9me3 enrichment in both Δ*cdp-2* and Δ*cdp-2;*Δ*dim-2* strains. (B-C) Average enrichment heatmaps of (B) H4K16ac or (C) H3K9me3 levels in WT, Δ*cdp-2*, and Δ*cdp-2;*Δ*dim-2* strains centered over WT-defined constitutive heterochromatic regions, as in Figures 3B-C.

To assess if enhanced heterochromatic acetylation coupled with 5^m^C loss impacts genome organization, we performed euchromatin-specific (*Dpn*II) *in situ* Hi-C to generally assess genome organization of the Δ*cdp-2*;Δ*dim-2* strain. We generated two highly correlated *in situ Dpn*II Hi-C replicates of a Δ*cdp-2*;Δ*dim-2* strain (Figure S23), which we merged into a single *Dpn*II Δ*cdp-2*;Δ*dim-2* Hi-C dataset containing 23.1M valid read pairs (Table S1). The contact frequency heatmap for the Δ*cdp-2*;Δ*dim-2* strain still shows some segregation of heterochromatin from euchromatin, as observed across each individual chromosome (Figure S24) and the entire genome (Figure S25). The KR-corrected whole genome contact probability heatmap also shows centromere bundles independent of telomere clusters typical of the Rabl conformation (Figure S25B).

However, comparison of equal numbers of valid read pairs between the Δ*cdp-2*;Δ*dim-2* and WT *Dpn*II Hi-C datasets showed evidence of genome disorder in the double mutant, with heterochromatic regions in Δ*cdp-2*;Δ*dim-2* interacting more with surrounding euchromatin. Both raw and KR corrected heatmaps comparing WT and Δ*cdp-2*;Δ*dim-2* contact probabilities show that AT-rich, heterochromatic loci, including the centromere, gain interactions with flanking euchromatic DNA across all LGs (Figure S26), including LG II (Figure 5A). Notably, the increased interactions between heterochromatic regions evident upon CDP-2 loss (e.g., Figures 1B-D) are not apparent in a Δ*cdp-2*;Δ*dim-2* strain. Further, obvious reductions in local and regional euchromatic interactions are observed (Figure 5A). We quantified the intra-chromosomal contact probability changes between the Δ*cdp-2*;Δ*dim-2* and WT datasets among bins enriched for heterochromatic or euchromatic histone PTMs. Few H3K9me3-enriched bins had strongly increased contacts with other heterochromatic bins, but more H3K9me3-marked bins gained contacts with euchromatic bins enriched for SET-2 H3K36me3 or H3K27ac; most contact probability gains occur between euchromatic bins (Figure 5B). Comparison of WT and Δ*cdp-2*;Δ*dim-2* datasets across the whole genome, independent of KR correction, shows increases in inter-chromosomal contact probability between heterochromatin and euchromatin, which depletes local intra-chromosomal contacts (Figure S27). Quantifying the strongly increased inter-chromosomal contacts shows H3K9me3-marked bins gaining interactions with bins enriched with euchromatic SET-2-dependent H3K36me2 and H3K27ac; the number of H3K27ac-enriched bins gaining inter-chromosomal contacts to SET-2-dependent H3K36me2-enriched bins is also increased (Figure 5B). We conclude that the Δ*cdp-2*;Δ*dim-2* heterochromatin bundle is disordered with histone hyperacetylation and 5^m^C depletion.

**Figure 5.**
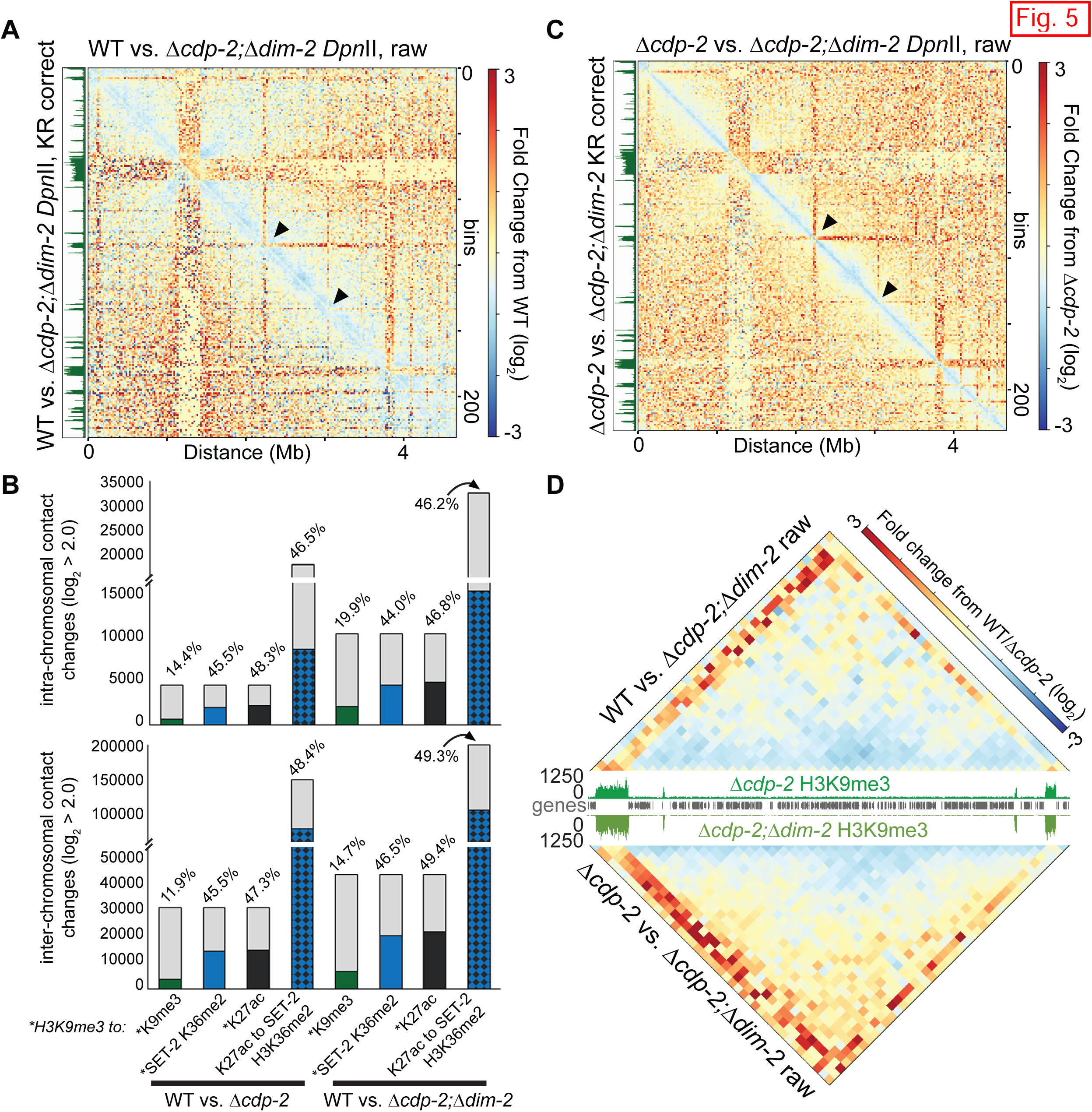
A Δcdp-2;Δdim-2 strain gains cross-compartment contacts. (A,C) Heatmaps of raw (above diagonal) or KR-corrected (below diagonal) contact probability changes across LG II in a Δ*cdp-2*;Δ*dim-2 Dpn*II *in situ* Hi-C dataset relative to (A) WT or (C) Δ*cdp-2* datasets. Arrowheads indicate the heterochromatic regions in panel D. (B) Quantification of strongly changed intra-chromosomal (top) or inter-chromosomal (bottom) contacts in Δ*cdp-2* or Δ*cdp-2*;Δ*dim-2* datasets relative to a WT dataset, as in Figure 1E. (D) Heatmap of the raw contact probability changes of two heterochromatic regions in a Δ*cdp-2*;Δ*dim-2* strain relative to (top) WT or (bottom) Δ*cdp-2* strains. IGV image of Δ*cdp-2* or Δ*cdp-2*;Δ*dim-2* H3K9me3 enrichment is in the middle.

We compared the *Dpn*II Hi-C data of the Δ*cdp-2* single mutant and the Δ*cdp-2*;Δ*dim-2* double mutant to highlight the genome organization changes caused by both hyperacetylation and loss of 5^m^C. Across LG II, the Δ*cdp-2*;Δ*dim-2* strain presents with stronger gains in contact frequency between heterochromatin and euchromatin, increases in regional euchromatic contacts across TAD-like structures, and decreased local (on-diagonal) contacts relative to a Δ*cdp-2* strain (Figure 5C); similar changes occur over the entire Neurospora genome and across the other LGs (Figures S28, S29). Closer examination of the regional contact changes between AT-rich loci and euchromatin highlight how loss of both CDP-2 and DIM-2 impacts chromosome conformation. Relative to WT, individual H3K9me3-enriched loci in a Δ*cdp-2*;Δ*dim-2* strain more strongly contact euchromatin, thereby depleting euchromatic clustering; this phenotype is exacerbated when comparing Δ*cdp-2*;Δ*dim-2* and Δ*cdp-2* contact probabilities (Figure 5D). We conclude that heterochromatic hyperacetylation and 5^m^C loss compromises the normal chromosome conformation in Neurospora nuclei.

## Discussion

In this work, we examined the role of CDP-2 and CHAP, as members of the constitutive heterochromatin-specific histone deacetylase complex HCHC, in modulating histone PTM enrichment and the genome organization of *Neurospora crassa*. Here, heterochromatin hyperacetylation in a Δ*cdp-2* strain causes increased contact probability within heterochromatic loci, as well as between distant intra– and inter-chromosomal heterochromatic regions, while CHAP loss reduces heterochromatin aggregation without strong histone acetylation changes. Neither gene deletion abolished the Rabl chromosome conformation typical of fungal genomes (28), suggesting the overall genome organization can still form despite HCHC loss. Together, our data suggest that CDP-2 and CHAP have distinct functions in the HCHC complex that differentially impact genome organization. Our model (Figure 6) is that CDP-2 recruits the HDA-1 deacetylase to properly compact individual heterochromatic regions while CHAP mediates heterochromatic region aggregation at the nuclear periphery after deacetylation. Specifically, the hyperacetylation seen with CDP-2 loss could open chromatin fibers and/or loosen the DNA wrapped about histones in heterochromatic nucleosomes, which would increase heterochromatin accessibility to allow greater contact probability with distant heterochromatic nucleosomes in the silent “B” compartment at the nuclear periphery (14) and indirectly increase the association of euchromatic regions across chromosome arms. One intriguing hypothesis is that the increased chromatin accessibility that occurs from enhanced heterochromatic nucleosome turnover, as observed in HDAC mutants in *S. pombe* and *N. crassa* (78, 79), could amplify the contact probability within and between heterochromatic regions upon CDP-2 loss. Regarding CHAP, this HCHC complex member compacts heterochromatic regions into dense structures and mediates inter-heterochromatic region contacts, possibly through direct AT-rich DNA binding by its AT-hook motifs (39). Since CDP-2 and CHAP recruit HDA-1 to AT-rich loci (39), these proteins most likely act prior to HDA-1, meaning the genome organizational changes we observe reflect inadequate HDA-1 targeting or deficient HCHC complex formation or function. Notably, the loss of the fourth HCHC complex member, HP1, has a genome organization distinct from that of Δ*cdp-2* or Δ*chap* strains, as flanking euchromatin emanating from heterochromatic regions has reduced contact probabilities in WT vs. Δ*hpo* datasets, similar to that of a Δ*dim-5* strain lacking H3K9me3 (27), however *in situ* Hi-C could reveal other Δ*hpo*-specific genome organization changes. Thus, HP1 binding to H3K9me3 presumably occurs prior to, or in parallel with, CHAP condensing AT-rich regions; CHAP and HP1 could both act prior to CDP-2 binding HP1 for HDA-1 recruitment (39). This proposed mechanism for Neurospora HCHC activity may be homologous to yeast HDAC complexes. In *Saccharomyces cerevisiae*, Sirtuin-2 (Sir2; a Class III HDAC)-specific deacetylation is required for the binding of the Sir3/Sir4 dimer onto deacetylated histone tails to form silenced superstructures at heterochromatic loci in the budding yeast genome (50, 58, 80, 81). *S. cerevisiae* also employs the HDACs Rpd3 (Class I) and HDA-1 (Class II) for euchromatic deacetylation (58, 82–84); Rpd3 forms two distinct HDAC complexes, the Rpd3L (large) complex targeting promoters and the Rpd3S (small) complex deacetylating gene bodies (84–86). In *Schizosaccharomyces pombe*, the multisubunit HDAC complex SHREC (***S***nf2/***H***dac-containing ***Re***pressor ***C***omplex) is targeted to heterochromatic loci by the HP1-homolog Swi6 for transcriptional gene silencing, although SHREC can localize to euchromatin independent of Swi6 (87–89). The SHREC catalytic subunit Clr3 is homologous to Neurospora HDA-1, and at least three other HDA-1 paralogs and a Rpd3 homolog exist in Neurospora (41). Notably, we used the general HDAC inhibitor Trichostatin A (TSA) to examine if all HDACs have roles in genome organization, yet TSA severely compromised WT growth, suggesting HDAC paralogs also influence fungal genome function. Examination of other HDACs in Neurospora by Hi-C or ChIP-seq could elucidate how HDAC paralogs impact chromosome conformation or histone PTM deposition.

**Figure 6.**
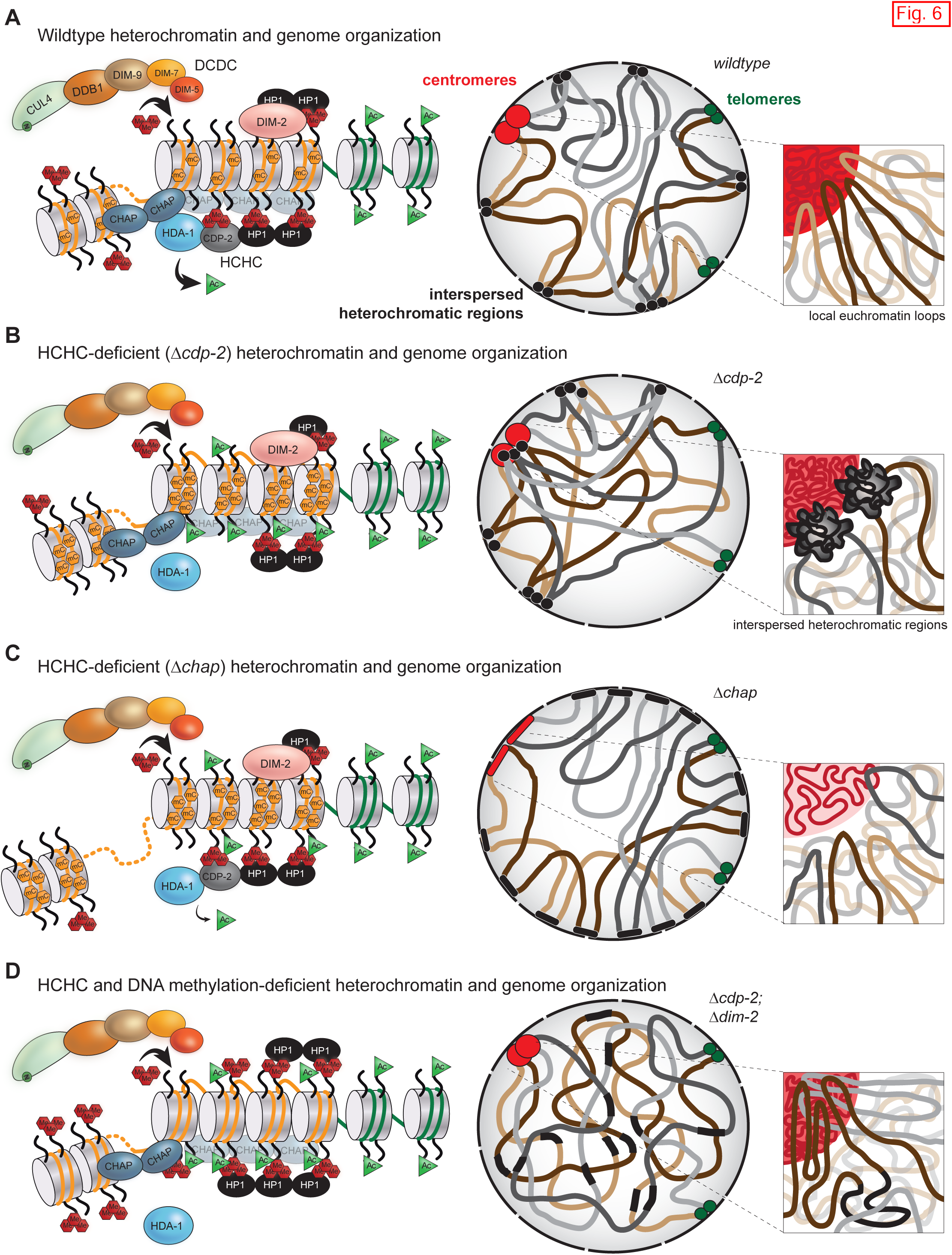
Models for how HCHC loss impacts heterochromatin formation and genome organization in Neurospora crassa. (A-D) For WT or HCHC mutant strains, heterochromatin formation at nucleosomes (left) and the organization of two example chromosomes (right) are presented. (A) In a WT strain, the HCHC (HDA-1, CDP-2, HP1, and CHAP) complex removes acetyl marks (green triangles) from histone tails. CDP-2 and HP1, which binds H3K9me3 (red hexagons) while CHAP binds AT-rich DNA (orange lines) between distant nucleosomes to recruit HDA-1. The DCDC (DIM-5/7/9, CUL4, DDB1*^dim-8^*, Complex) tri-methylates H3K9 on deacetylated histone tails, and HP1 binds H3K9me3 to directly recruit the DNA methyltransferase DIM-2 for cytosine methylation (orange hexagons). The activity of both HCHC and DCDC establishes the Rabl chromosome conformation characterized by a centromere (red circles) cluster distinct from (sub)telomeres bundles (green circles) and the clustering of interspersed heterochromatic regions (black circles), all of which occur at the nuclear periphery. (B) CDP-2 loss increases heterochromatic histone acetylation and alters (here, gains) 5^m^C while maintaining H3K9me3 at most AT-rich loci, which together increases intra-heterochromatic region contacts and decreases inter-chromosomal centromeric contacts. (C) CHAP loss relaxes heterochromatic region compaction and reduces inter-heterochromatic region interactions, despite increases in 5^m^C and H3K9me3. (D) Loss of both CDP-2 and DIM-2 increases histone acetylation, H3K9me3 enrichment, and genome disorder as heterochromatic regions amalgamate with euchromatin.

As previously shown by ChIP qPCR, multiple histone lysines become hyperacetylated upon loss of HDA-1 and CDP-2; the extent of hyperacetylation depends on both the heterochromatic region assessed and which HCHC member was deleted (40). This published ChIP qPCR data, as well as our genome-wide ChIP-seq acetylation data, hints at the HCHC complex having preferential deacetylation activity for certain lysine residues in histones. Specifically, our ChIP-seq data shows heterochromatin in a Δ*cdp-2* strain becoming hyperacetylated at H4K16 but minimally changed at H3K9, while heterochromatic H4K16ac enrichment is minimally altered in a Δ*chap* strain; perhaps HP1 or CDP-2, both of which can bind to H3K9me3 (40, 90, 91), can compensate for CHAP loss and still allow for HDA-1 deacetylation. The unchanged H3K9ac enrichment in a Δ*cdp-2* strain may reflect DCDC-dependent (38) tri-methylation of H3K9, given the mutually exclusive deposition of either acetylation or trimethylation on H3K9 (92, 93). Currently, it is unknown if H4K16ac is globally enriched in Δ*hpo* heterochromatin, or whether other histone acetyl marks have altered enrichment in HCHC mutants. Future ChIP-seq experiments on other acetylated histone lysine residues in Δ*cdp-2,* Δ*chap*, Δ*hpo*, or Δ*hda-1* strains should uncover genome-wide changes to acetylation patterns for understanding the lysine specificity of the HCHC complex.

Interestingly, while Δ*cdp-2* or Δ*chap* strains gain heterochromatin-specific H4K16ac relative to WT, these mutant strains display no change in euchromatic acetylation, suggesting the HCHC complex specifically deacetylates heterochromatic histones. Since both the class II HDAC HDA-1 in Neurospora and the class III HDACs Hst2p and Sir2 in *S. cerevisiae* are both known to deacetylate H4K16 (63, 81), yet they share little primary structure similarity, it is possible that different HDACs can deacetylate identical histone lysines but the targeting mechanism of each HDAC complex determines which genomic loci are deacetylated. Given that HDA-1 is targeted to heterochromatic nucleosomes by CHAP binding AT-rich DNA and HP1/CDP-2 binding H3K9me3 (39, 40), these factors would render the HCHC complex as a heterochromatin-specific HDAC. This explains the copious HCHC mutants recovered in a selection for strains with compromised heterochromatin silencing (94); it is possible the HCHC complex is the only heterochromatin specific HDAC in Neurospora. Our data also suggest that unknown HAT(s) specifically acetylate heterochromatic H4K16 – which is only seen with compromised HCHC HDAC activity – given that euchromatic acetylation is minimally altered in HCHC mutants. Presumably, Neurospora could repress HCHC expression or activity to hyperacetylate heterochromatin for altering genome organization at certain developmental or cell cycle stages, including S phase in which a more open chromosome conformation exists in fission yeast (95). Conversely, Neurospora may increase expression of the HCHC complex and/or other HDACs to hypoacetylate mitotic chromatin for segregating condensed chromosomes to daughter cells; rapid HDAC repression would allow chromatin to be rapidly acetylated following cytokinesis, as previously observed (63, 64). It is noteworthy that the increased contact probability occurring between hyperacetylated, yet distant, heterochromatic regions in Δ*cdp-2* nuclei is analogous to the clustering of hyperacetylated chromatin in BRD4-NUT expressing oncogenic cells (68). Together, changes in acetylation patterns that alter chromatin accessibility could represent a simple mechanism that eukaryotes employ to differentially organize their genomes.

Curiously, smaller AT-rich regions in a Δ*cdp-2* strain lose H3K9me3 yet this mark remains unchanged at larger heterochromatic loci, while a Δ*chap* strain has modest H3K9me3 gains at most AT-rich regions, highlighting the differential effects of HCHC mutants on heterochromatic histone PTMs. Apart from size, other possible causes of locus-specific H3K9me3 loss in CDP-2 deficient strains include compromised subnuclear positioning, reduced pairing with other AT-rich regions (96), or unrecognized transposon relict motifs in the underlying DNA (97). In contrast, the H3K9me3 gain in a Δ*chap* strain may result from increased DCDC levels or activity, which would promote further AT-rich region compaction through additional HP1 binding sites. Similar increases in the DCDC subunits DIM-7 and DIM-9 occur in a *dim-3* strain to possibly rescue reduced H3K9me3 levels (98). Our data suggest that WT heterochromatic nucleosomes are not saturated with H3K9me3, given that greater levels of H3K9me3 are possible in some mutant HCHC strains. Notably, changes in H3K9me3 deposition can impact the enrichment of the H3K27me2/3 histone PTM that marks facultative heterochromatin, for in DCDC deficient strains, H3K27me2/3 relocates to AT-rich genomic loci (99, 100). While ChIP qPCR showed no H3K27me2/3 change at a few loci in HCHC deletion strains (99), examination of the global enrichment of H3K27me2/3 in HCHC deficient backgrounds would be prudent, given the changes in H3K9me3 deposition reported here.

Changes in H3K9me3 are most apparent in a Δ*cdp-2*;Δ*dim-2* double mutant, where most heterochromatic regions gain considerable H3K9me3 enrichment, although smaller AT-rich regions similarly lose H3K9me3 as a single Δ*cdp-2* mutant; all AT-rich loci gain H4K16ac in both strains. These results suggest the loss of 5^m^C signals for the H3K9me3 increase but does not affect histone acetylation. Importantly, the genome organization of the double mutant is disordered, as AT-rich heterochromatic regions amalgamate with euchromatin despite greater H3K9me3 deposition (and presumably HP1 binding). These results may explain the severe growth defect of Δ*cdp-2*;Δ*dim-2* strains (40), as improper positioning of AT-rich regions into the nucleus center may cause aberrant transcription of repetitive DNA or limit normal genic transcription, the latter being particularly deleterious given how heterochromatic-euchromatic contacts mediate proper gene expression in filamentous fungi (16, 101). Perhaps the increased H3K9me3 enrichment in the Δ*cdp-2*;Δ*dim-2* strain is a “last resort” by the fungus to repress AT-rich region transcription in the nucleus center, where increased H3K9me3 and HP1 compaction could minimize RNA Pol II recruitment to this mislocalized chromatin. A crucial, unanswered question is: “how is H3K9me3 enrichment increased?” We speculate that reduced AT-rich region compaction, observed in both Δ*chap* and Δ*cdp-2*;Δ*dim-2* strains, signals for increased H3K9me3 deposition. Consistent with this hypothesis, intergenic region nucleosomal disorder causes subtle increases in H3K9me3 in Neurospora *dim-1* strains (102), but this remains to be seen in HCHC mutants.

Importantly, our double mutant Hi-C results suggest that cytosine methylation maintains chromatin compartmentalization by segregating hyperacetylated heterochromatin from euchromatin. Specifically, only in strains devoid of both CDP-2 and DIM-2 are AT-rich regions more apt to contact euchromatin. However, this change only moderately affects the global genome topology, since the Rabl chromosome conformation is still maintained, although the weak chromosome territories typically seen in Neurospora are diminished in Δ*cdp-2*;Δ*dim-2* nuclei. These results suggest heterochromatin can cluster, as in WT, but unlike the situation in a Δ*cdp-2* strain, the inter-heterochromatic contact gains are abrogated by 5^m^C loss. We speculate the additional methyl groups on AT-rich DNA in a Δ*cdp-2* strain allow heterochromatic regions to remain associated with the nuclear periphery, either by active recruitment by a methyl-DNA binding protein (MRBP-1) (103, 104) or passively by Liquid-Liquid Phase Separation (105). The identity of MRBP-1 in Neurospora is unknown, as Neurospora does not encode a protein with a strong MBD domain homologous to human MeCP-2, but perhaps DIM-10, which compromises heterochromatin silencing but not 5^m^C deposition or H3K9me3 enrichment (94), could bind methylated DNA in Neurospora. Regardless, our results suggest a new role for 5^m^C in maintaining chromatin compartmentalization when heterochromatin is compromised, thereby safeguarding genome organization and function. Future experiments should discern if 5^m^C has a similar role in maintaining genome topology in fungal pathogens or higher eukaryotes.

## Materials and Methods

The strains used for all experiments, the abbreviated Hi-C and ChIP-seq protocols, and the bioinformatic methods are found in the online Materials and Methods, provided in Supplementary File 1. *In situ* Hi-C was performed as in (16), using the program hicExplorer (73) to process mapped reads for building the contact matrices, which were used for all downstream analyses and image generation. ChIP-seq performed as in (32, 102), with minor modifications. The complete *in situ* Hi-C protocol is provided in Supplementary File 2, while the ChIP-seq protocol is found in Supplementary File 3.

## Data availability

All *Neurospora crassa* strains are available upon request. The *in situ* Hi-C and ChIP-seq high-throughput sequencing data are available in the NCBI GEO superseries accession number GSE232935, which includes the ChIP-seq data accession number GSE232933 and the Hi-C data accession number GSE232934.

## Supporting information

Scadden_Supplementary-Fig_S1

Scadden_Supplementary-Fig_S2

Scadden_Supplementary-Fig_S3

Scadden_Supplementary-Fig_S4

Scadden_Supplementary-Fig_S5

Scadden_Supplementary-Fig_S6

Scadden_Supplementary-Fig_S7

Scadden_Supplementary-Fig_S8

Scadden_Supplementary-Fig_S9

Scadden_Supplementary-Fig_S10

Scadden_Supplementary-Fig_S11

Scadden_Supplementary-Fig_S12

Scadden_Supplementary-Fig_S13

Scadden_Supplementary-Fig_S14

Scadden_Supplementary-Fig_S15

Scadden_Supplementary-Fig_S16

Scadden_Supplementary-Fig_S17

Scadden_Supplementary-Fig_S18

Scadden_Supplementary-Fig_S19

Scadden_Supplementary-Fig_S20

Scadden_Supplementary-Fig_S21

Scadden_Supplementary-Fig_S22

Scadden_Supplementary-Fig_S23

Scadden_Supplementary-Fig_S24

Scadden_Supplementary-Fig_S25

Scadden_Supplementary-Fig_S26

Scadden_Supplementary-Fig_S27

Scadden_Supplementary-Fig_S28

Scadden_Supplementary-Fig_S29

Scadden_Supplementary-File_S1

Scadden_Supplementary-File_S2

Scadden_Supplementary-File_S3

Scadden_Supplementary-Table_S1

## Acknowledgements

The authors wish to thank UCCS Andrew Reckard, Sara Rodriguez, Yulia Shtanko, Victoria Toscano, and Farh Kaddar (UCCS) for assistance with library generation, preliminary data analysis, or critical reading of the manuscript, Doug Turnbull and Jeff Bishop (University of Oregon Genomics and Cell Characterization Core Facility) for Illumina library sequencing service, and all Klocko lab members and colleagues of the UCCS Department of Chemistry & Biochemistry for helpful comments and discussions. Funding was provided by start-up funds from the UCCS College of Letters, Arts, and Sciences and an Academic Research Enhancement Award (AREA) grant from the National Institutes of Health (1R15GM140396-01) to A.D.K.

