## Supplementary material for "Histone deacetylation and cytosine methylation compartmentalize heterochromatic regions in the genome organization of *Neurospora crassa*": Scadden_Supplementary-Fig_S2

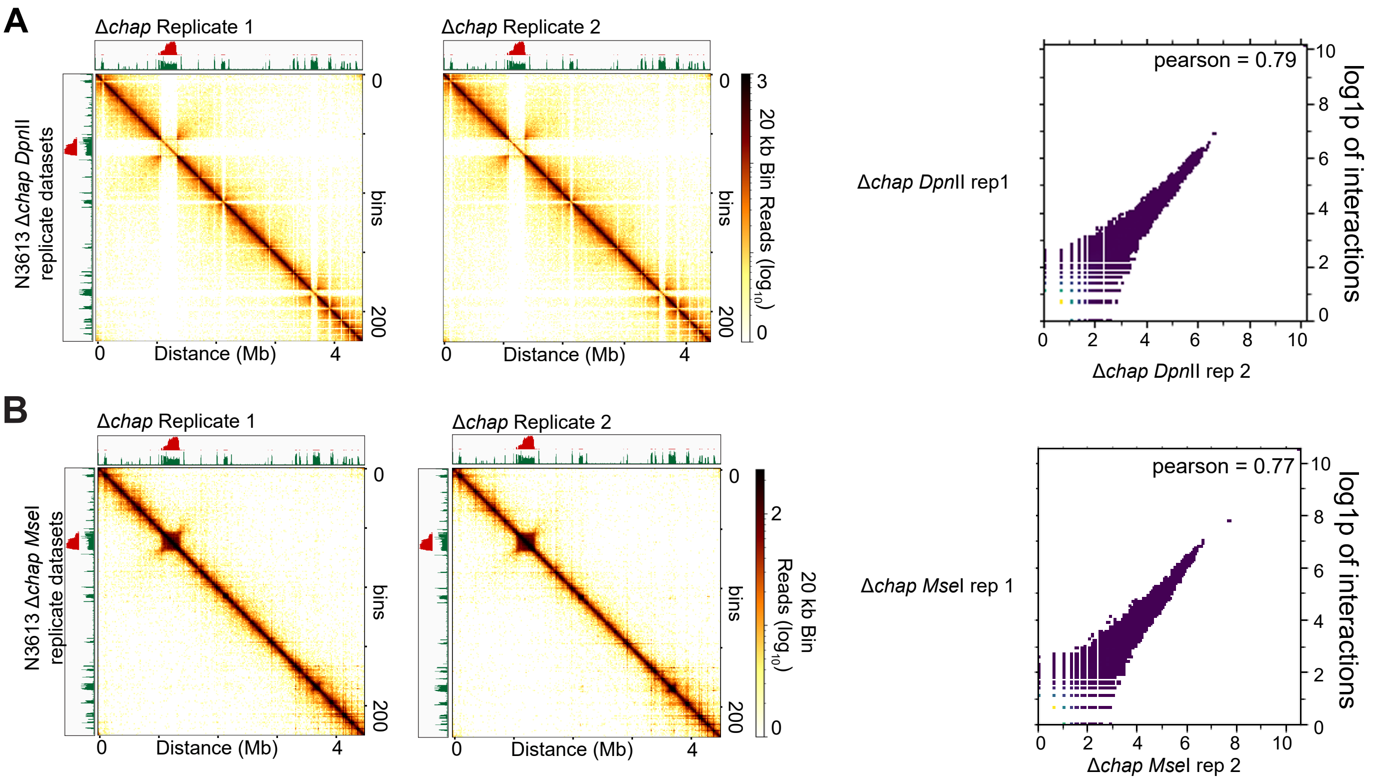


**Figure S2. Replicate analysis of *in situ* Hi-C genome organization datasets, using the *Dpn*II or *Mse*I restriction enzymes, of the Δ*chap* strain of *Neurospora crassa*.** (A-B) Heatmaps and scatterplots of Δ*chap* replicates digested with (A) *Dpn*II or (B) *Mse*I. In each panel, the left images show the raw count Hi-C heatmaps of genomic interactions across LG II of *in situ* Hi-C replicate datasets at 20 kb bin resolution. The right images show scatter plots comparing interactions between each replicate matrix at 20 kb bin resolution; log1p values of interactions presented. Pearson correlation values of each binary comparison shown at the upper right. Image produced by the hicCorrelate program in hicExplorer.
