## Supplementary material for "Histone deacetylation and cytosine methylation compartmentalize heterochromatic regions in the genome organization of *Neurospora crassa*": Scadden_Supplementary-Fig_S4

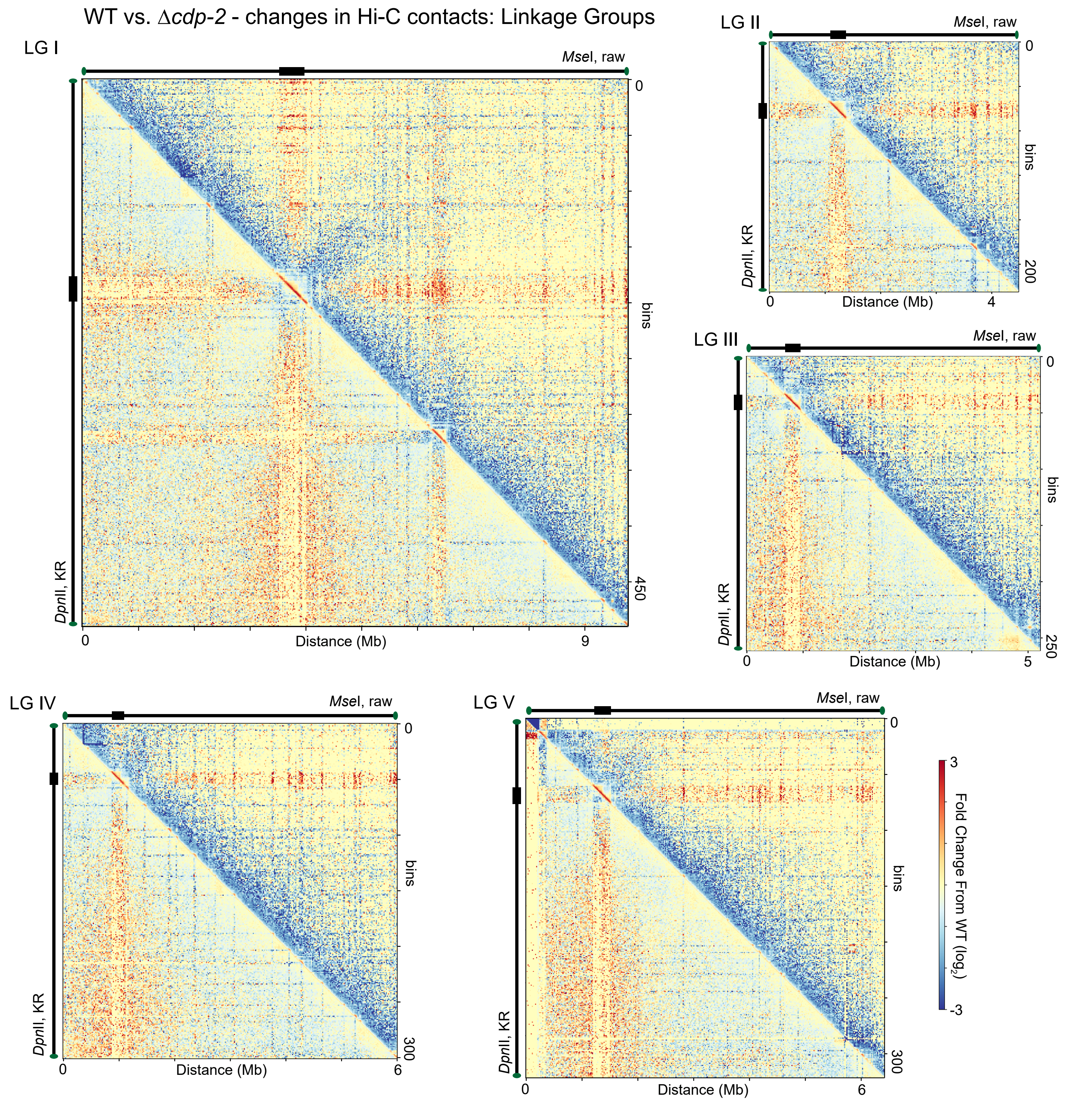


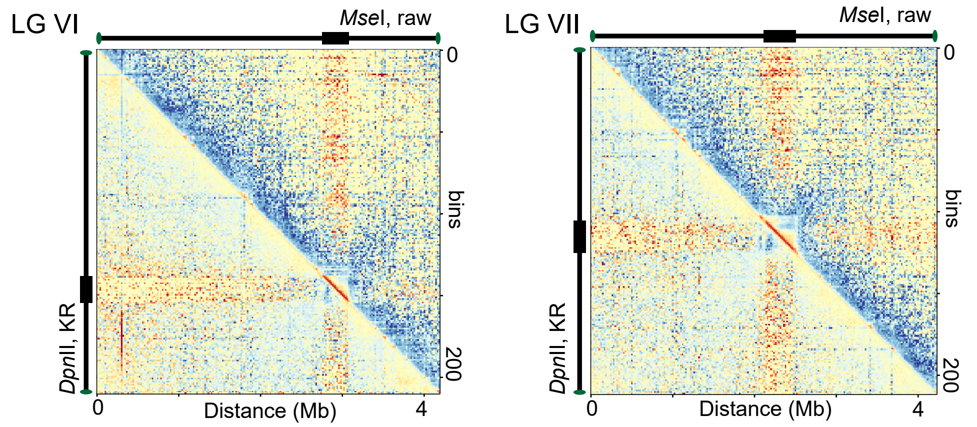


**Figure S4. Changes in chromosome conformation of the seven individual chromosomes between the WT and Δ*cdp-2* strains of Hi-C datasets generated using the restriction enzymes *Mse*I and *Dpn*II.** Heatmaps displaying $\log_2$ difference in *in situ* Hi-C contacts, either in *Mse*I raw read count datasets (above diagonal) or the datasets of *Dpn*II KR-corrected counts (below diagonal) between the WT strain and the Δ*cdp-2* strain across the seven Linkage Groups (LG) at 20 kb bins. Chromosomes are sized according to their length.
