## Supplementary material for "Histone deacetylation and cytosine methylation compartmentalize heterochromatic regions in the genome organization of *Neurospora crassa*": Scadden_Supplementary-Fig_S5

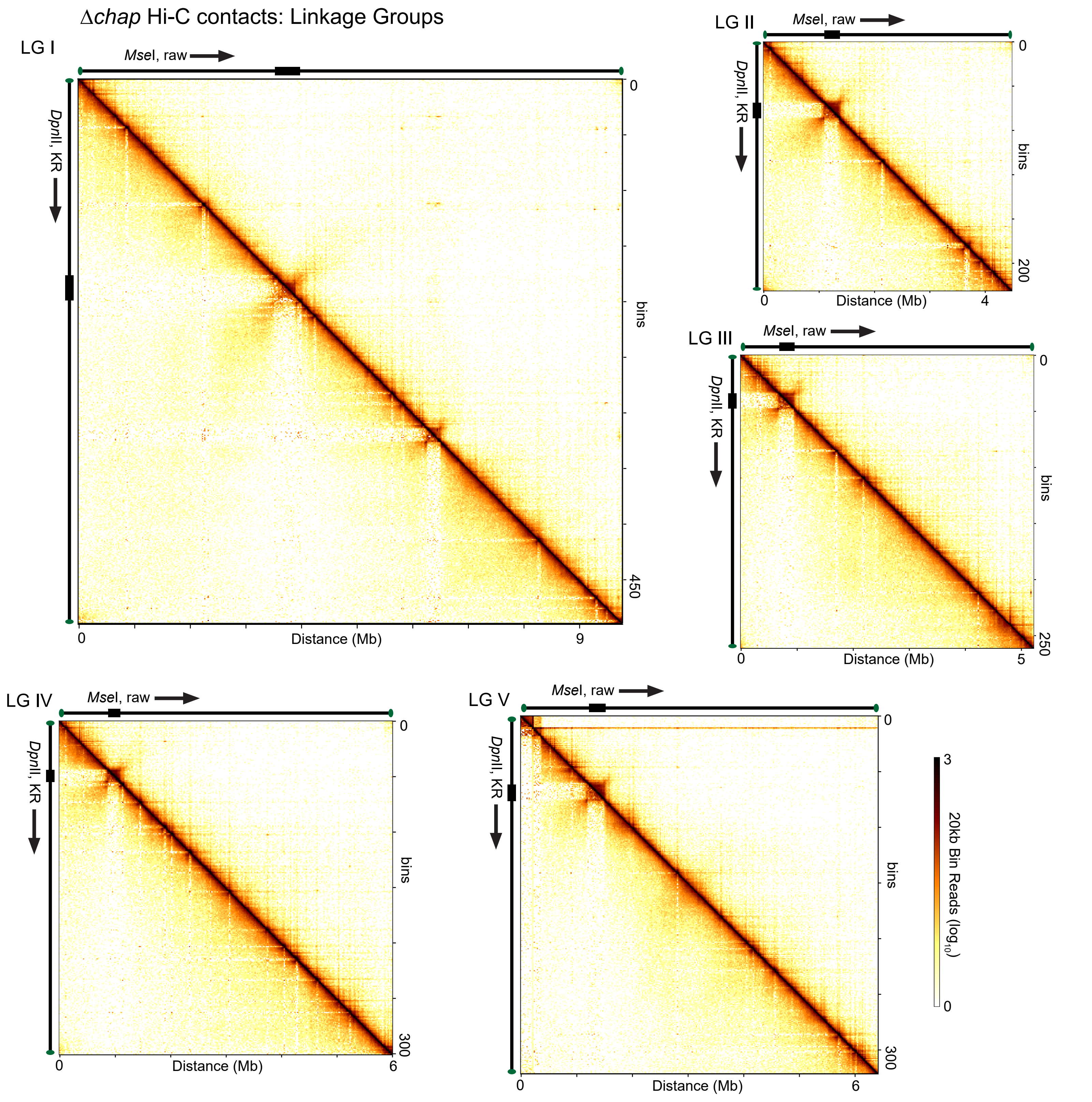


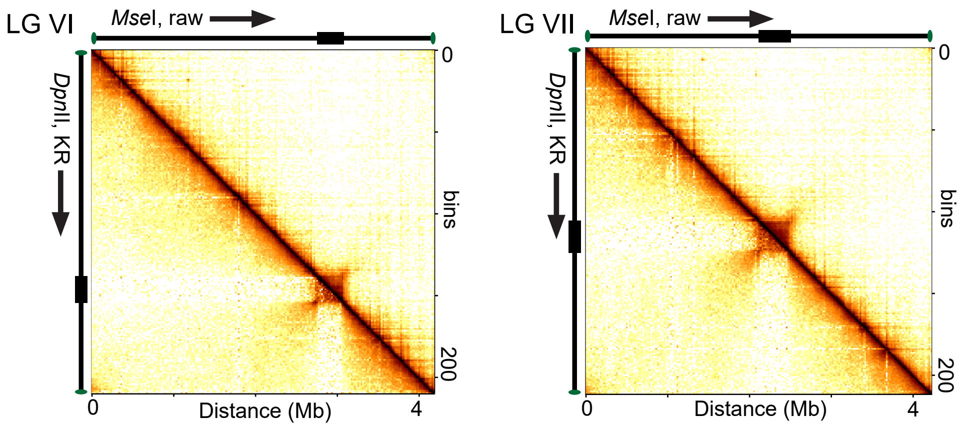


**Figure S5. Chromosome conformation of the seven individual chromosomes of the Δ*chap* strain using the restriction enzymes *Mse*I and *Dpn*II.** Heatmaps displaying *in situ* Hi-C contacts, either as *Mse*I raw read counts (above diagonal) or *Dpn*II KR-corrected counts (below diagonal) for the Δ*chap* strain across the seven Linkage Groups (LG) at 20 kb bins. Chromosomes are sized according to their length.
