## Supplementary material for "Histone deacetylation and cytosine methylation compartmentalize heterochromatic regions in the genome organization of *Neurospora crassa*": Scadden_Supplementary-Fig_S6

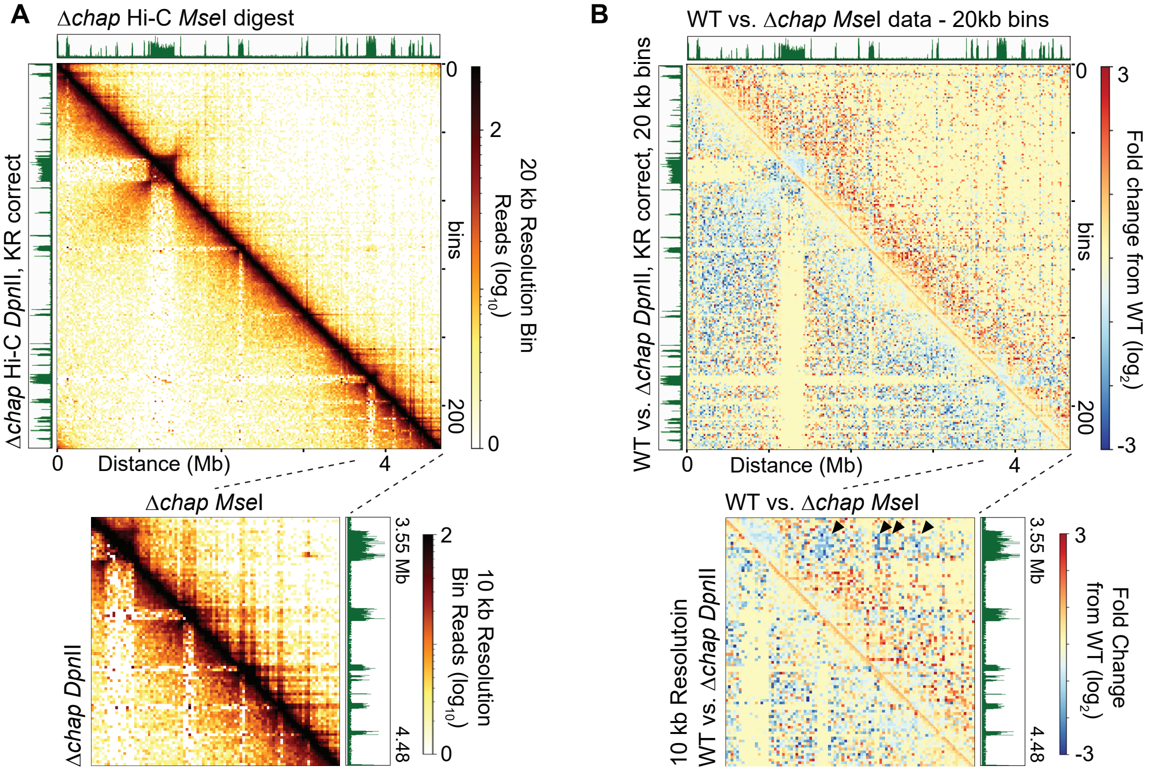


**Figure S6. Regional examination of the Δ*chap* dataset shows dense coverage of Hi-C data that is different from WT.** (A) Heatmap of contact probabilities of 20 kb bins across LG II of a Δ*chap* strain, displayed, as in Figure 1A. Heatmap below shows the contact probabilities of the right arm of LG II of the Δchap dataset at 10 kb resolution; H3K9me3 enrichment shown to the right. (B) The fold change in contact strength ($log_2$ scale) in a Δ*chap* strain relative to a normalized WT strain, displayed as in Figure 1C the entire LG II or the right arm of LG II (zoomed image below).
