## Supplementary material for "Histone deacetylation and cytosine methylation compartmentalize heterochromatic regions in the genome organization of *Neurospora crassa*": Scadden_Supplementary-Fig_S8

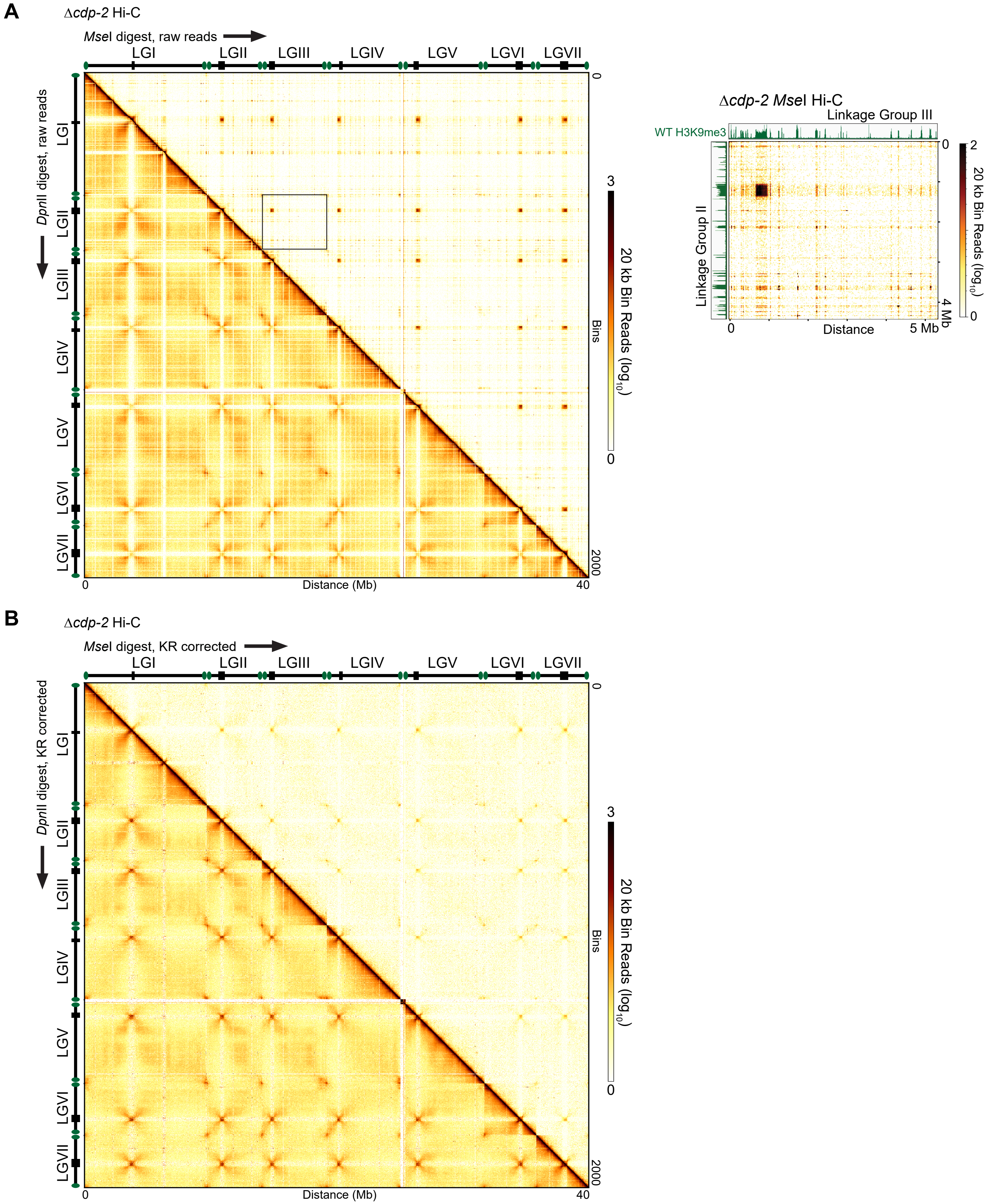


**Figure S8. Chromosome conformation of the whole genome of the Δ*cdp-2* strain using the restriction enzymes *Mse*I and *Dpn*II, either as raw read counts or KR corrected counts.** (A-B) Heatmaps displaying *in situ* Hi-C contacts, either as (A) raw read counts or (B) KR-corrected counts for the Δ*cdp-2* strain across the entire *Neurospora crassa* genome at 20 kb bins. *Mse*I derived Hi-C datasets are presented above the diagonal while *Dpn*II derived Hi-C datasets are shown below the diagonal. Enhanced region (box in A) shows the specific interactions between LG II and LG III of *Mse*I data from a Δ*cdp-2* strain.
