## Supplementary material for "Histone deacetylation and cytosine methylation compartmentalize heterochromatic regions in the genome organization of *Neurospora crassa*": Scadden_Supplementary-Fig_S9

**A**

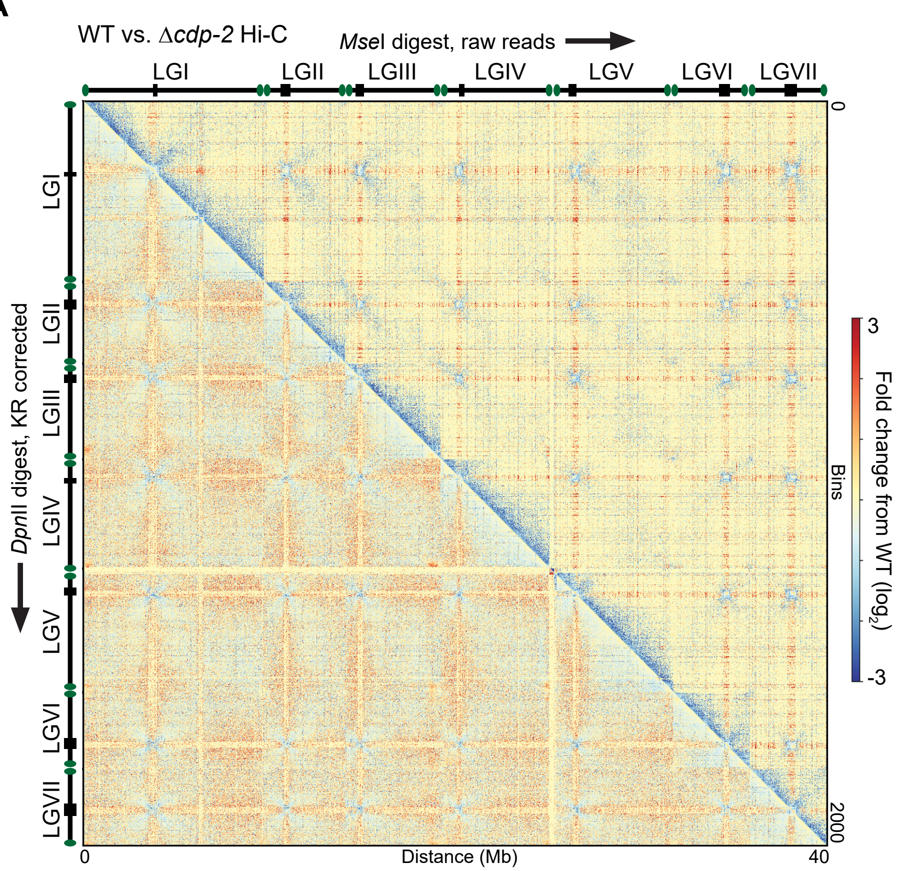


**B**

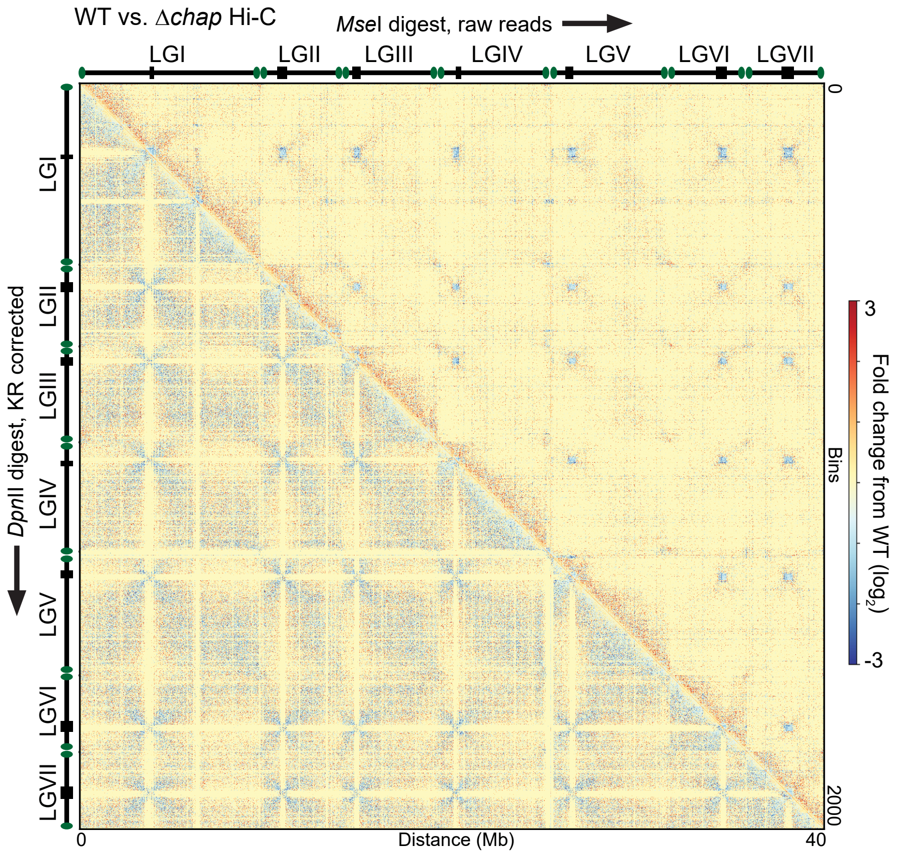


**Figure S9. Changes in the chromosome conformation across the whole genome of HCHC mutant strains, relative to WT strains, using the restriction enzymes *Mse*I and *Dpn*II.** (A-B) Heatmaps displaying the $\log_2$ difference in *in situ* Hi-C contacts, relative to WT, from either *Mse*I Hi-C datasets of raw read counts (above diagonal) or *Dpn*II Hi-C datasets of KR-corrected counts (below diagonal) for the (A) Δ*cdp-2* strain or (B) Δ*chap* strain across the entire *Neurospora crassa* genome at 20 kb bins.
