## Supplementary material for "Histone deacetylation and cytosine methylation compartmentalize heterochromatic regions in the genome organization of *Neurospora crassa*": Scadden_Supplementary-Fig_S10

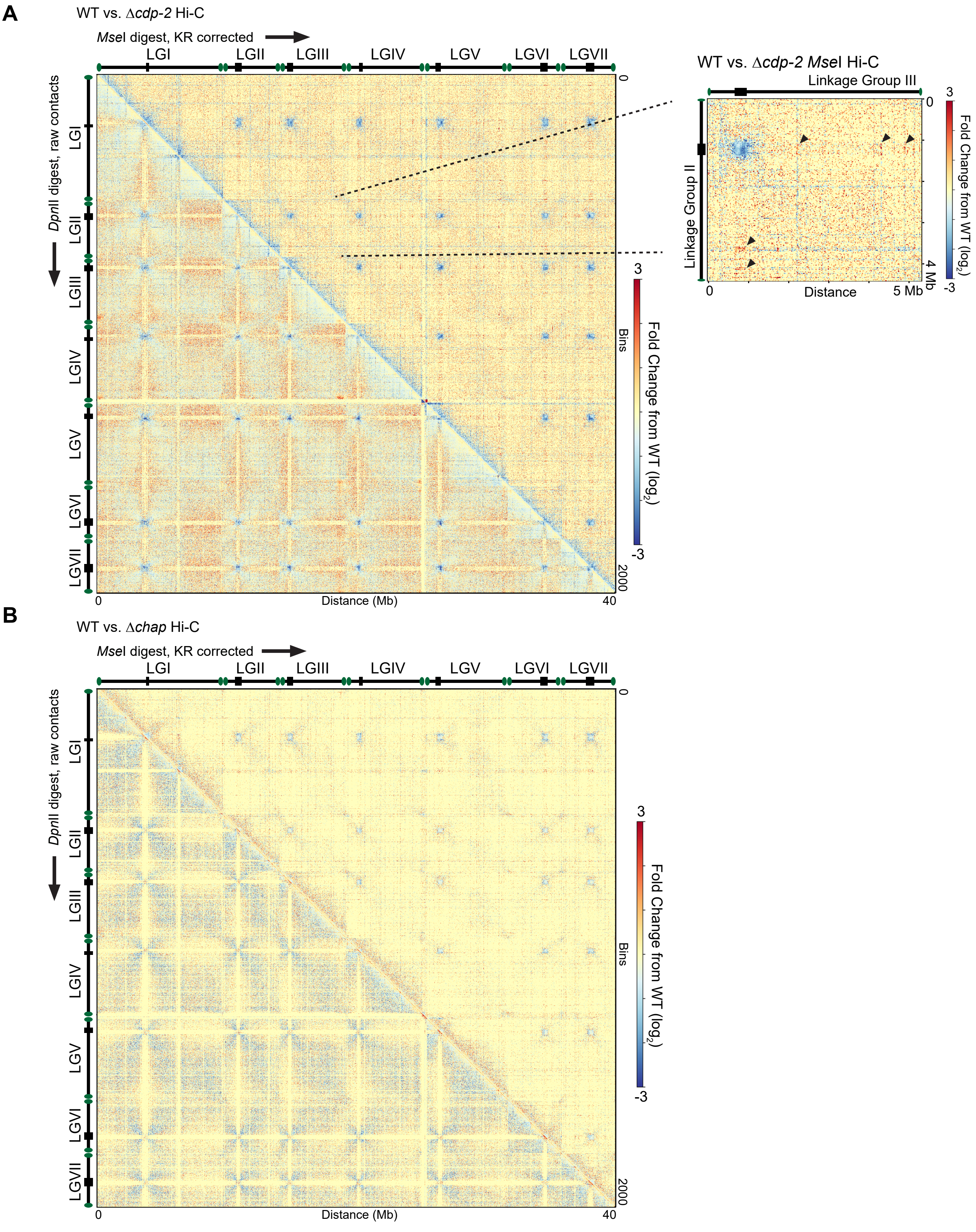


**Figure S10. Changes in the chromosome conformation across the whole genome of HCHC mutant strains, relative to WT strains, of KR-corrected *Mse*I and raw *Dpn*II data.** (A-B) Heatmaps displaying the $\log_2$ difference in *in situ* Hi-C contacts, relative to WT, from either *Mse*I Hi-C datasets of KR-corrected read counts (above diagonal) or *Dpn*II Hi-C datasets of raw counts (below diagonal) for the (A) Δ*cdp-2* strain or (B) Δ*chap* strain across the entire *Neurospora crassa* genome at 20 kb bins. Enhanced region in (A) shows the changes between two chromosomes, Linkage Groups II and III, for the Δ*cdp-2* strain; black arrowheads indicate increases in centromeric-heterochromatic interactions that are still observed in the KR corrected *Mse*I-data.
