## Supplementary material for "Histone deacetylation and cytosine methylation compartmentalize heterochromatic regions in the genome organization of *Neurospora crassa*": Scadden_Supplementary-Fig_S12

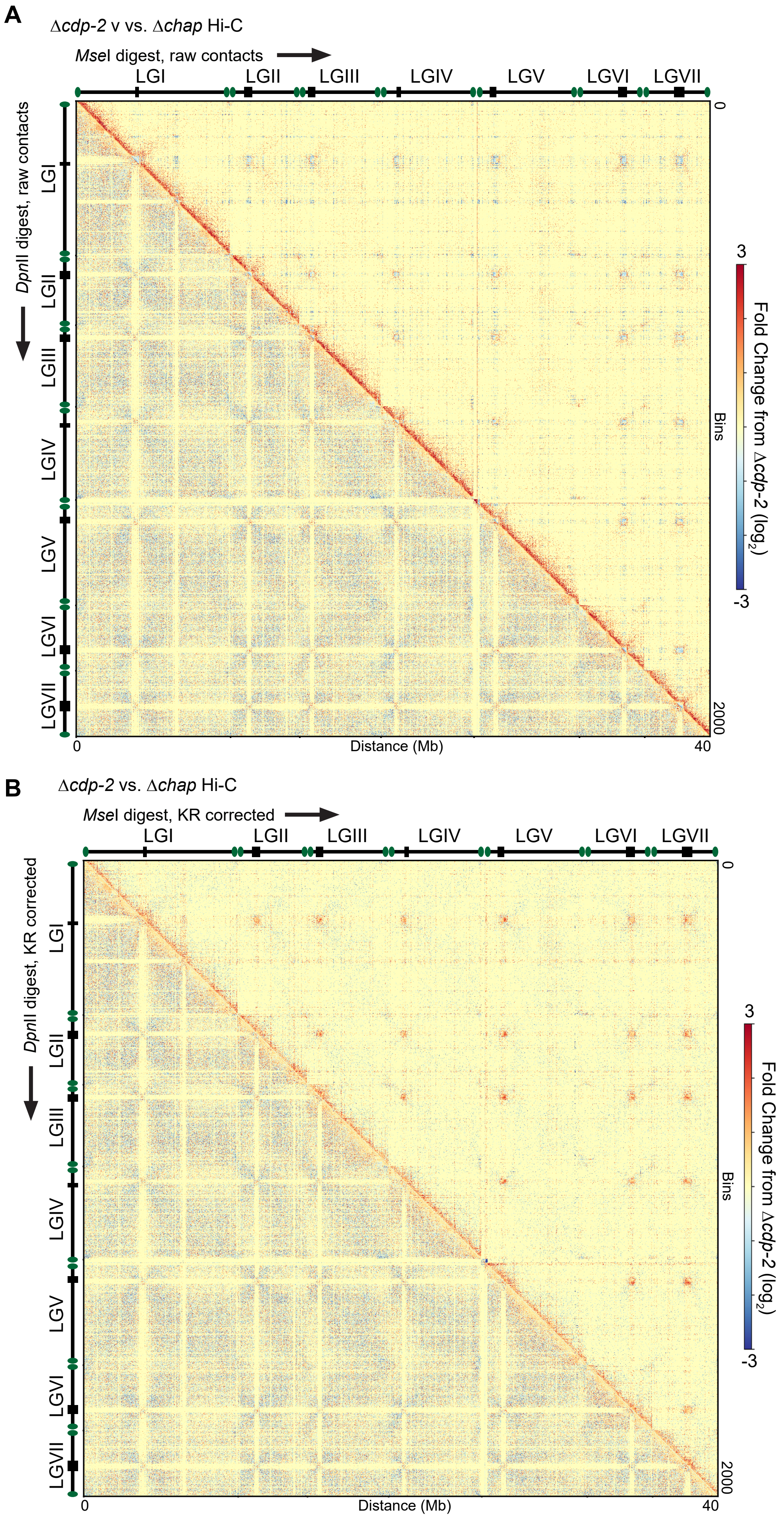


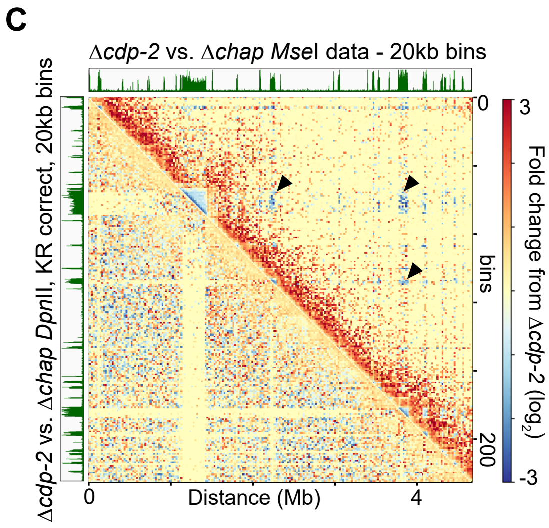


**Figure S12. Chromosome conformation changes across the whole genome, or a single LG, between the two HCHC mutant strains, using the restriction enzymes *Mse*I and *Dpn*II.** (A-B) Heatmaps displaying *in situ* Hi-C contacts, either as (A) raw read counts or (B) KR-corrected counts for the difference in Hi-C contacts of a Δ*chap* strain, relative to a Δ*cdp-2* strain, across the entire *Neurospora crassa* genome at 20 kb bins. *Mse*I derived Hi-C datasets are presented above the diagonal while *Dpn*II derived Hi-C datasets are shown below the diagonal. (C) The fold change in contact strength in a Δ*chap* strain relative to a Δ*cdp-2* strain across LG II, as displayed in Figure 1C. Arrowheads show the decrease in heterochromatic contacts in a Δ*chap* strain relative to a Δ*cdp-2* strain.
