## Supplementary material for "Histone deacetylation and cytosine methylation compartmentalize heterochromatic regions in the genome organization of *Neurospora crassa*": Scadden_Supplementary-Fig_S13

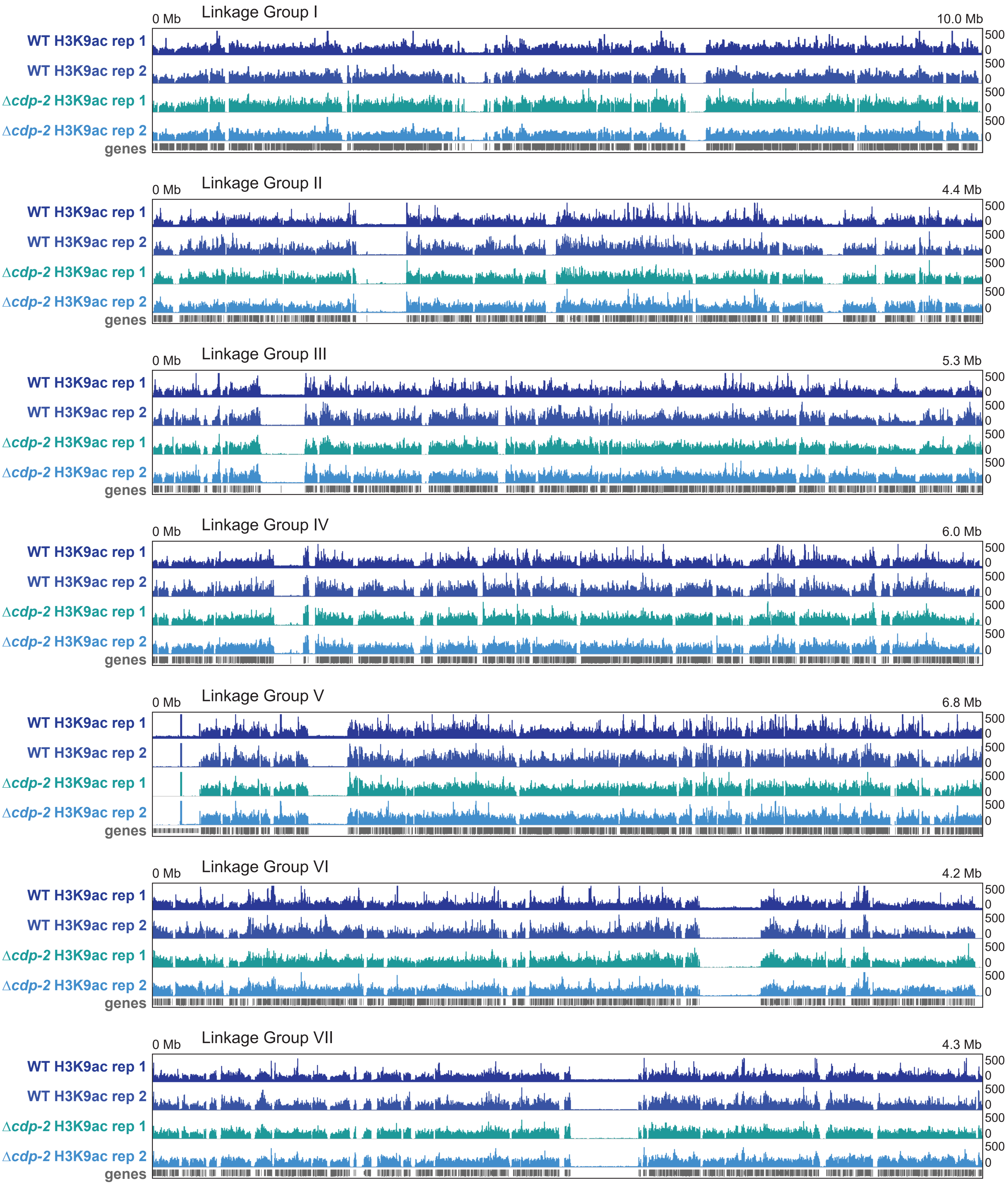


**Figure S13. Replicates of the H3K9ac Chromatin Immunoprecipitation-sequencing (ChIP-seq) of WT and Δ*cdp-2* strains are reproducible.** Integrative Genomics Viewer (IGV) images of the H3K9ac enrichment, assessed with bigwig files at 25 basepair (bp) resolution, across the seven Neurospora chromosomes (Linkage Groups). Wild type (WT) datasets shown in darker blue hues and Δ*cdp-2* datasets shown in lighter blue hues. Genes shown in gray. Enrichment values noted at the right, while chromosome distances presented at the top.
