## Supplementary material for "Histone deacetylation and cytosine methylation compartmentalize heterochromatic regions in the genome organization of *Neurospora crassa*": Scadden_Supplementary-Fig_S14

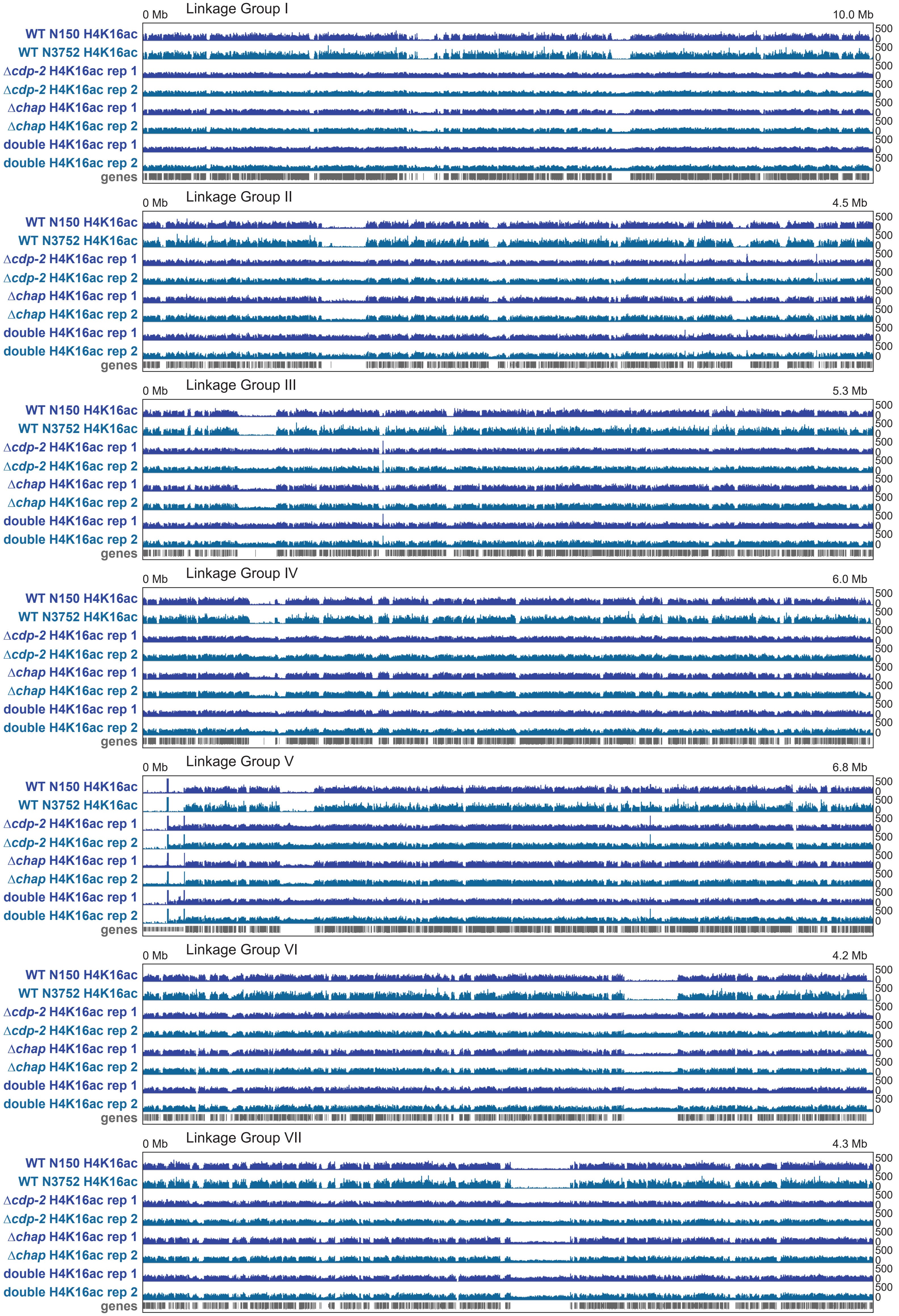


**Figure S14. Replicates of the H4K16ac Chromatin Immunoprecipitation-sequencing (ChIP-seq) of WT, Δ*cdp-2*, Δ*chap*, and Δ*cdp-2*;Δ*dim-2* strains are reproducible.** Integrative Genomics Viewer (IGV) images of the H4K16ac enrichment, assessed with bigwig files at 25 basepair (bp) resolution, across the seven Neurospora chromosomes (Linkage Groups). The first replicate of each dataset pair is shown in a dark blue hue and while the second replicate dataset shown in a light blue hue. Genes shown in gray. Enrichment values noted at the right, while chromosome distances presented at the top.
