## Supplementary material for "Histone deacetylation and cytosine methylation compartmentalize heterochromatic regions in the genome organization of *Neurospora crassa*": Scadden_Supplementary-Fig_S15

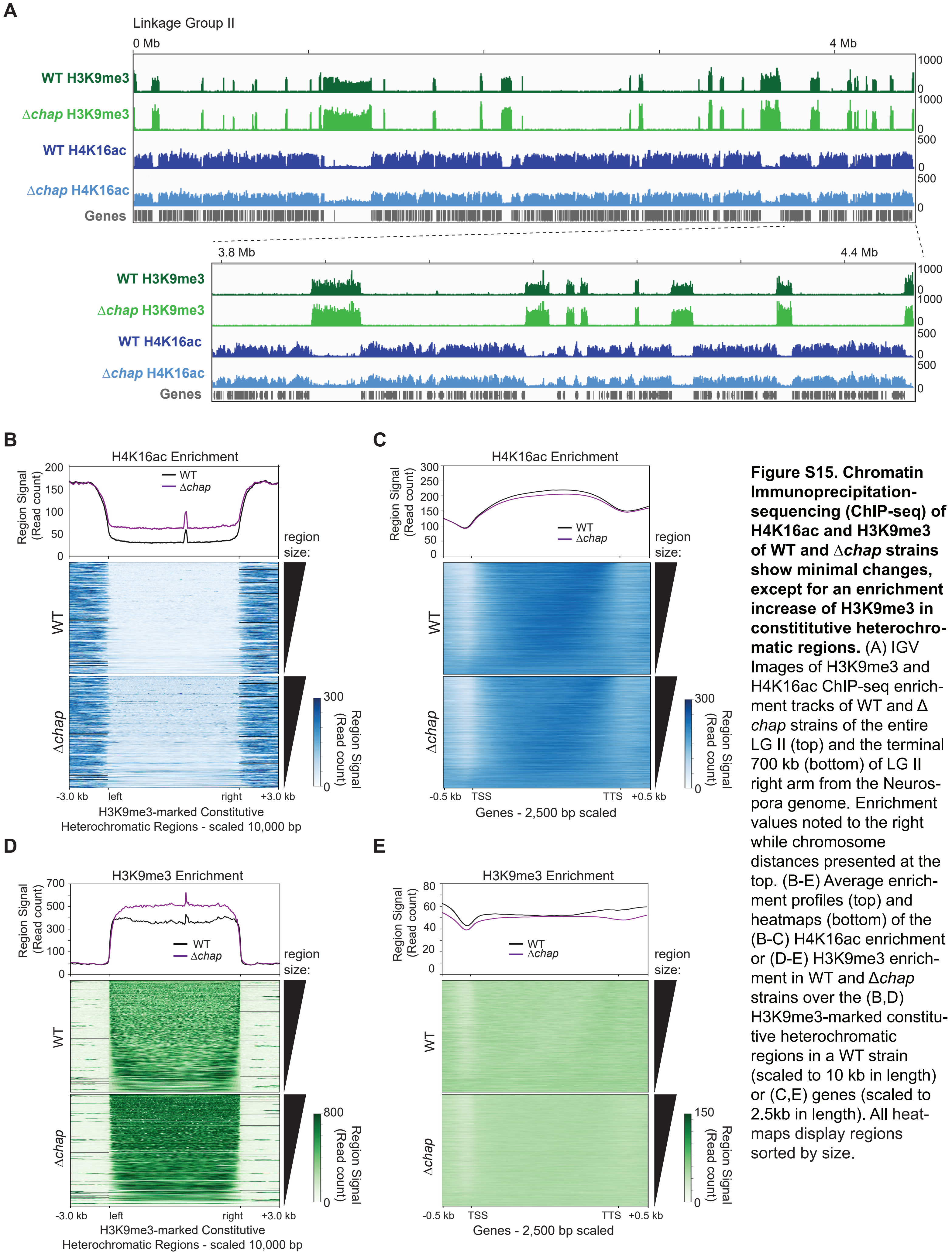


**Figure S15. Chromatin Immunoprecipitation-sequencing (ChIP-seq) of H4K16ac and H3K9me3 of WT and Δ*chap* strains show minimal changes, except for an enrichment increase of H3K9me3 in constititutive heterochromatic regions.** (A) IGV Images of H3K9me3 and H4K16ac ChIP-seq enrichment tracks of WT and Δ *chap* strains of the entire LG II (top) and the terminal 700 kb (bottom) of LG II right arm from the Neurospora genome. Enrichment values noted to the right while chromosome distances presented at the top. (B-E) Average enrichment profiles (top) and heatmaps (bottom) of the (B-C) H4K16ac enrichment or (D-E) H3K9me3 enrichment in WT and Δ*chap* strains over the (B,D) H3K9me3-marked constitutive heterochromatic regions in a WT strain (scaled to 10 kb in length) or (C,E) genes (scaled to 2.5kb in length). All heatmaps display regions sorted by size.
