## Supplementary material for "Histone deacetylation and cytosine methylation compartmentalize heterochromatic regions in the genome organization of *Neurospora crassa*": Scadden_Supplementary-Fig_S16

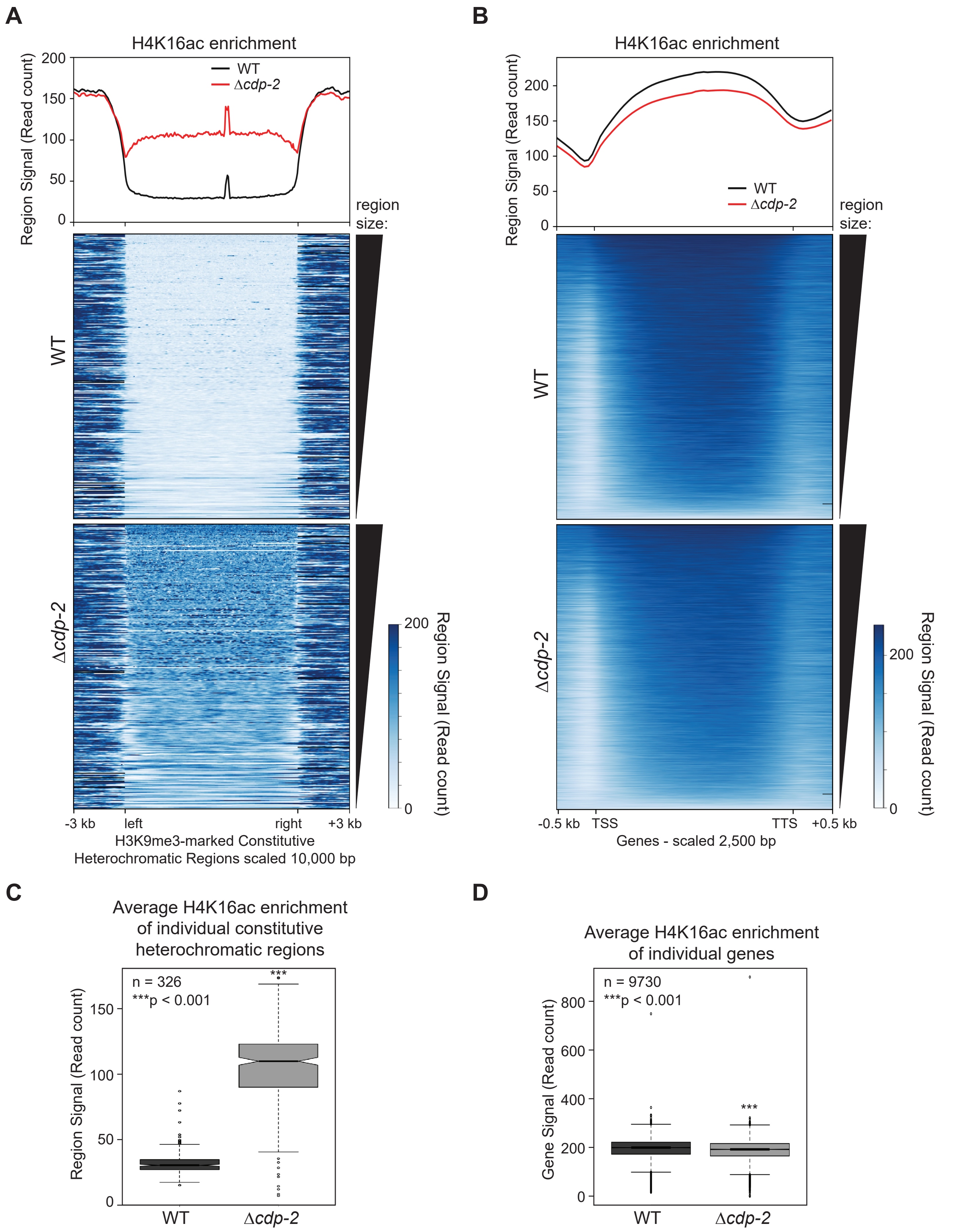


**Figure S16. Enrichment of H4K16ac over constitutive heterochromatic regions is significantly increased but essentially unchanged over genes in a Δ*cdp-2* strain.** (A-B) Average enrichment profiles (top) and heatmaps (bottom) of the H4K16ac enrichment in WT and Δ*cdp-2* strains over (A) heterochromatic regions (scaled to 10 kb in length) or (B) genes (scaled to 2.5 kb in length). Both heatmaps display regions sorted by size. (C-D) Boxplots of the normalized H4K16ac signal (see Materials and Methods) of each (C) heterochromatic region or (D) genes in WT and Δ*cdp-2* stains. Asterisks show significant ($p < 0.001$) differences in signal.
