## Supplementary material for "Histone deacetylation and cytosine methylation compartmentalize heterochromatic regions in the genome organization of *Neurospora crassa*": Scadden_Supplementary-Fig_S17

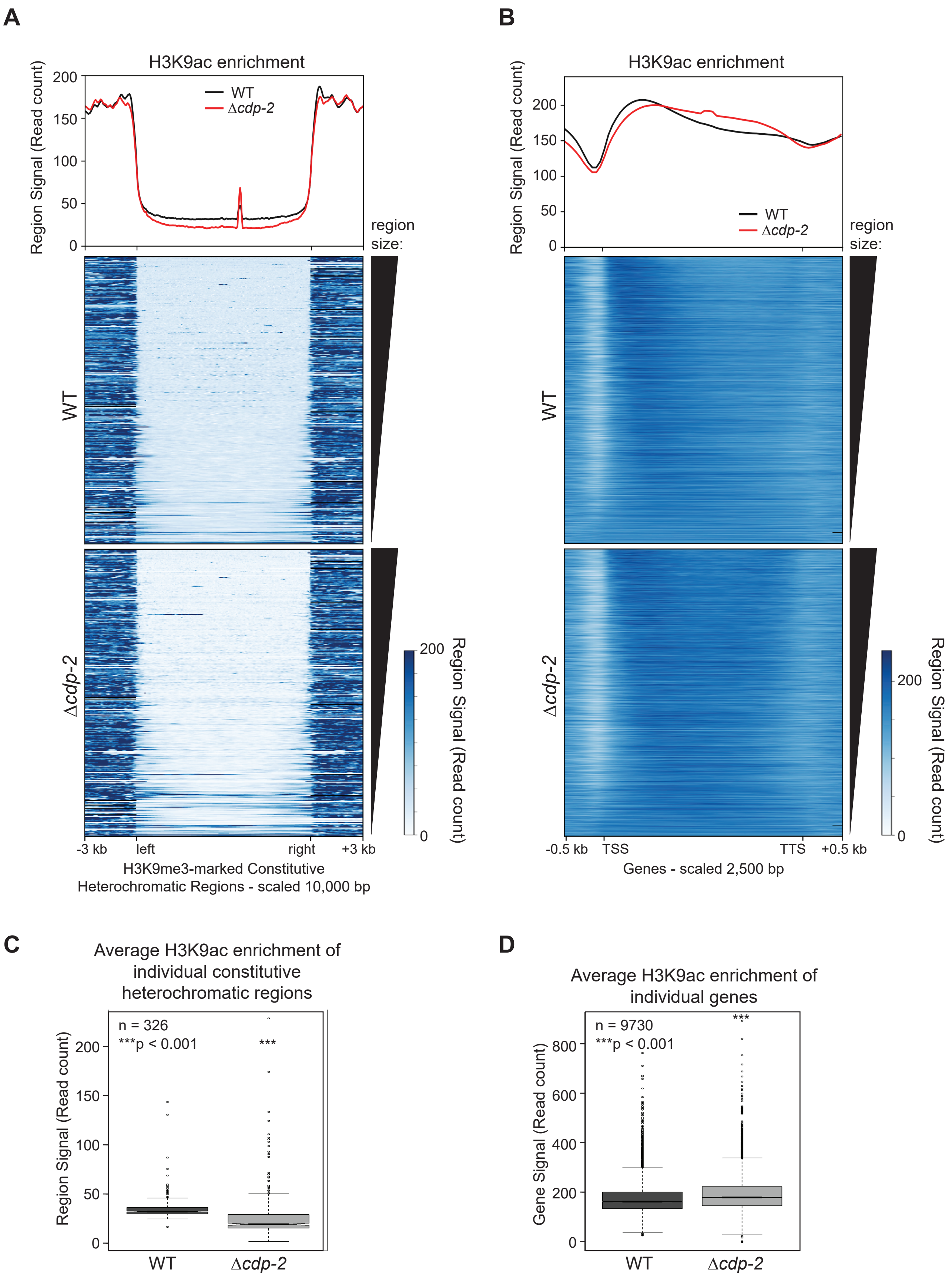


**Figure S17. Chromatin Immunoprecipitation-sequencing (ChIP-seq) average enrichment profiles of H3K9ac of WT and Δ*cdp-2* strains show minimal changes across constitutive heterochromatin and genes.** (A-B) Average enrichment profiles (top) and heatmaps (bottom) of the H3K9ac enrichment in WT and Δ*cdp-2* strains over the (A) H3K9me3-marked constitutive heterochromatic regions in a WT strain (scaled to 10 kb in length) or (B) genes (scaled to 2.5kb in length). Heatmaps are ordered by region length. (C-D) Boxplots of the normalized H3K9ac signal (see Materials and Methods) of each heterochromatic region in WT and Δ*cdp-2* stains. Asterisks show significant ($p < 0.001$) differences in signal.
