## Supplementary material for "Histone deacetylation and cytosine methylation compartmentalize heterochromatic regions in the genome organization of *Neurospora crassa*": Scadden_Supplementary-Fig_S18

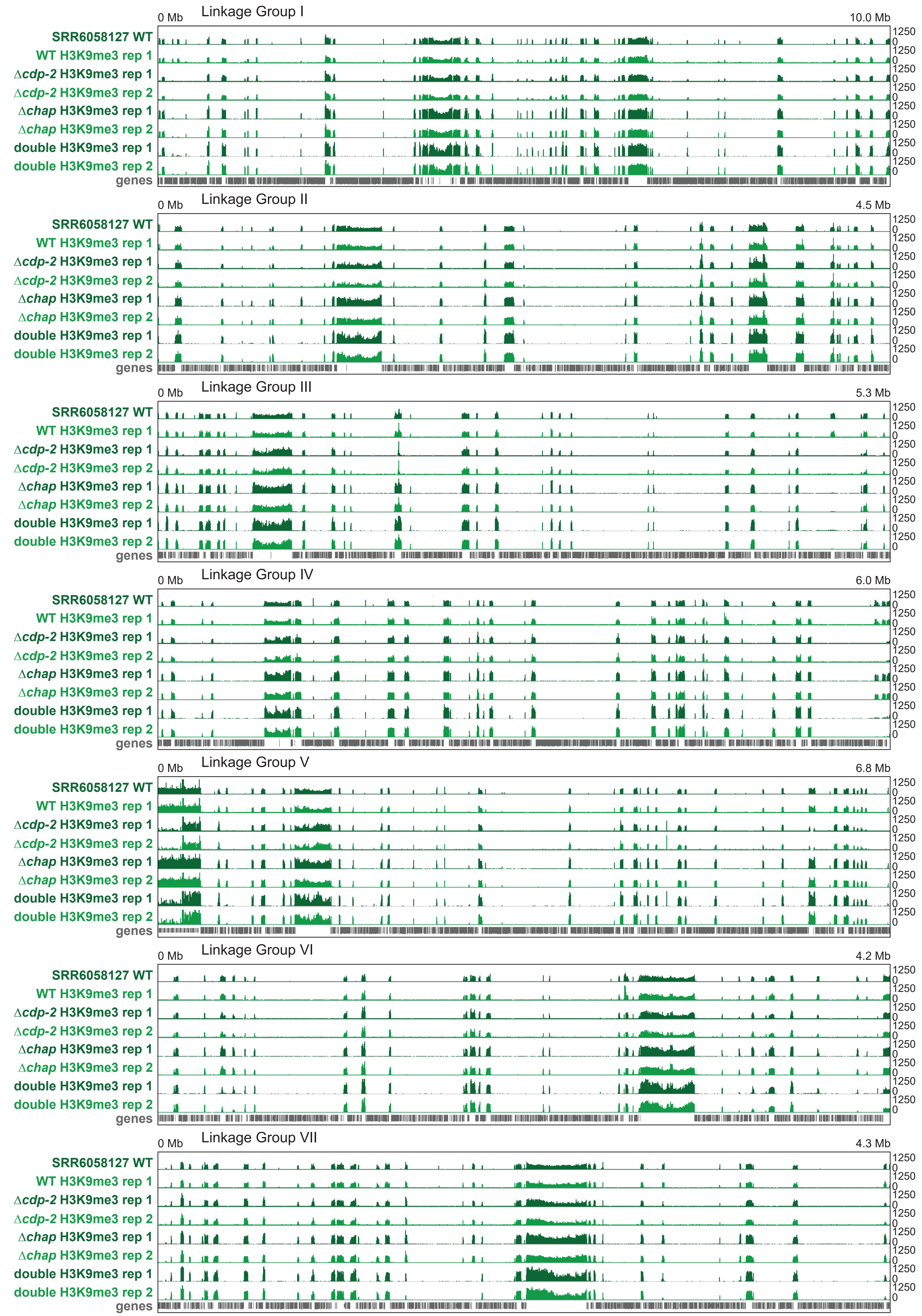


**Figure S18. Replicates of the H3K9me3 Chromatin Immunoprecipitation-sequencing (ChIP-seq) of WT, Δ*cdp-2,* Δ*chap*, and Δ*cdp-2;*Δ*dim-2* strains are reproducible.** Integrative Genomics Viewer (IGV) images of the H3K9me3 enrichment, assessed with bigwig files at 25 basepair (bp) resolution, across the seven Neurospora chromosomes (Linkage Groups). The first replicate of each dataset pair is shown in a dark green hue and while the second replicate dataset shown in a light green hue. Genes shown in gray. Enrichment values noted at the right, while chromosome distances presented at the top.
