## Supplementary material for "Histone deacetylation and cytosine methylation compartmentalize heterochromatic regions in the genome organization of *Neurospora crassa*": Scadden_Supplementary-Fig_S19

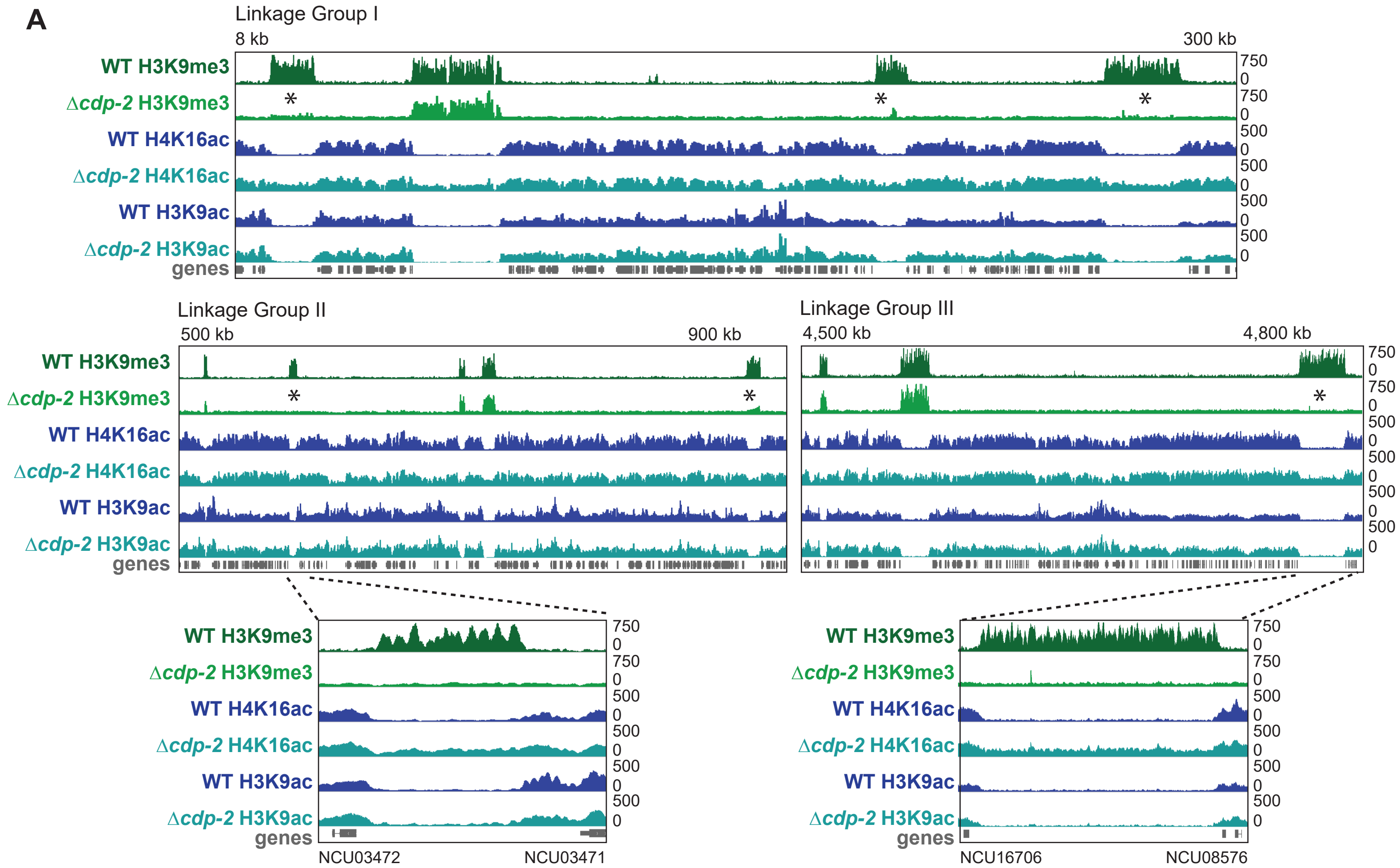


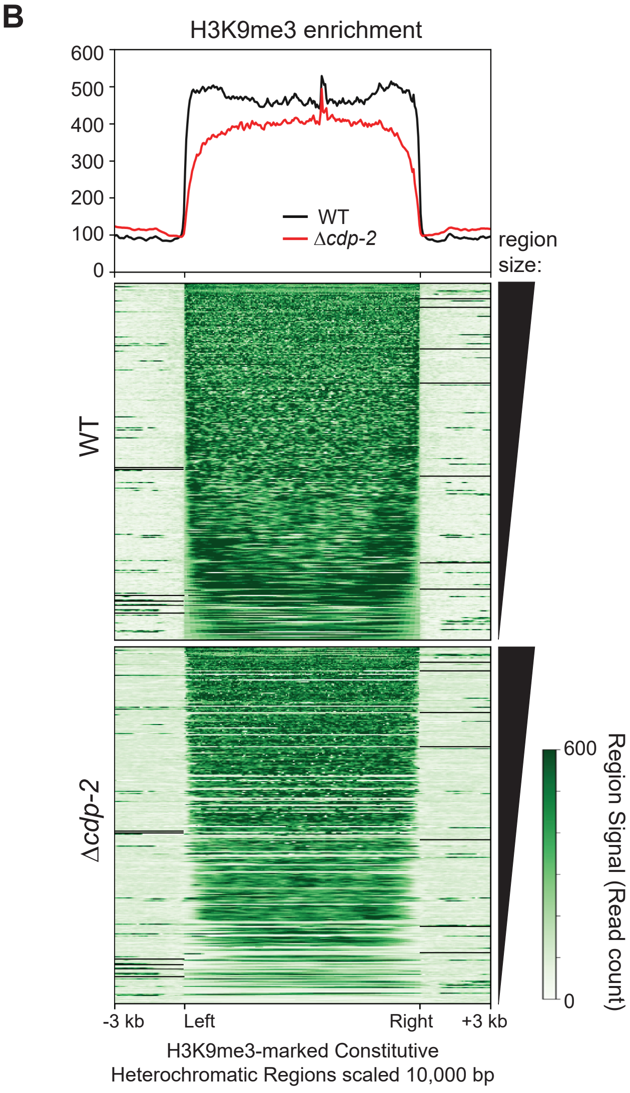


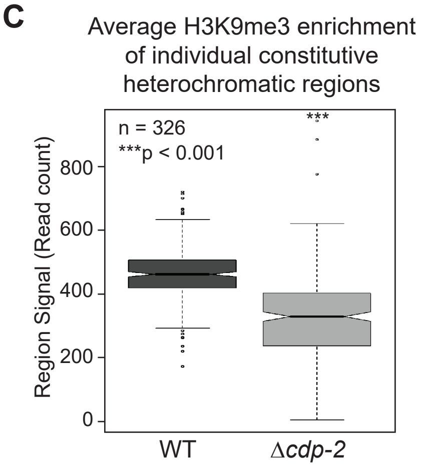


**Figure S19. Changes in H3K9me3 deposition and genome organization of constitutive heterochromatic regions upon CDP-2 loss across the Neurospora crassa genome.** (A) Images of H3K9me3, H4K16ac, and H3K9ac ChIP-seq enrichment tracks of WT and Δ*cdp-2* strains, displayed on IGV, of three regions on LG I, LG II, and LG III, and two further zoomed in images of heterochromatic regions on LG II and LG III. The enhanced images show the enrichment of H3K9me3, H4K16ac, and H3K9ac for individual heterochromatic regions. Asterisks show regions that lose H3K9me3 in a Δ*cdp-2* strain (B) Average enrichment profile (top) and heatmap (bottom) of the H3K9me3 enrichment in WT and Δ*cdp-2* strains over heterochromatic regions (scaled to 10 kb in length). The heatmap displays regions sorted by size.(C) Boxplot of the normalized H3K9me3 signal of each heterochromatic region in WT or Δ*cdp-2* stains.
