## Supplementary material for "Histone deacetylation and cytosine methylation compartmentalize heterochromatic regions in the genome organization of *Neurospora crassa*": Scadden_Supplementary-Fig_S20

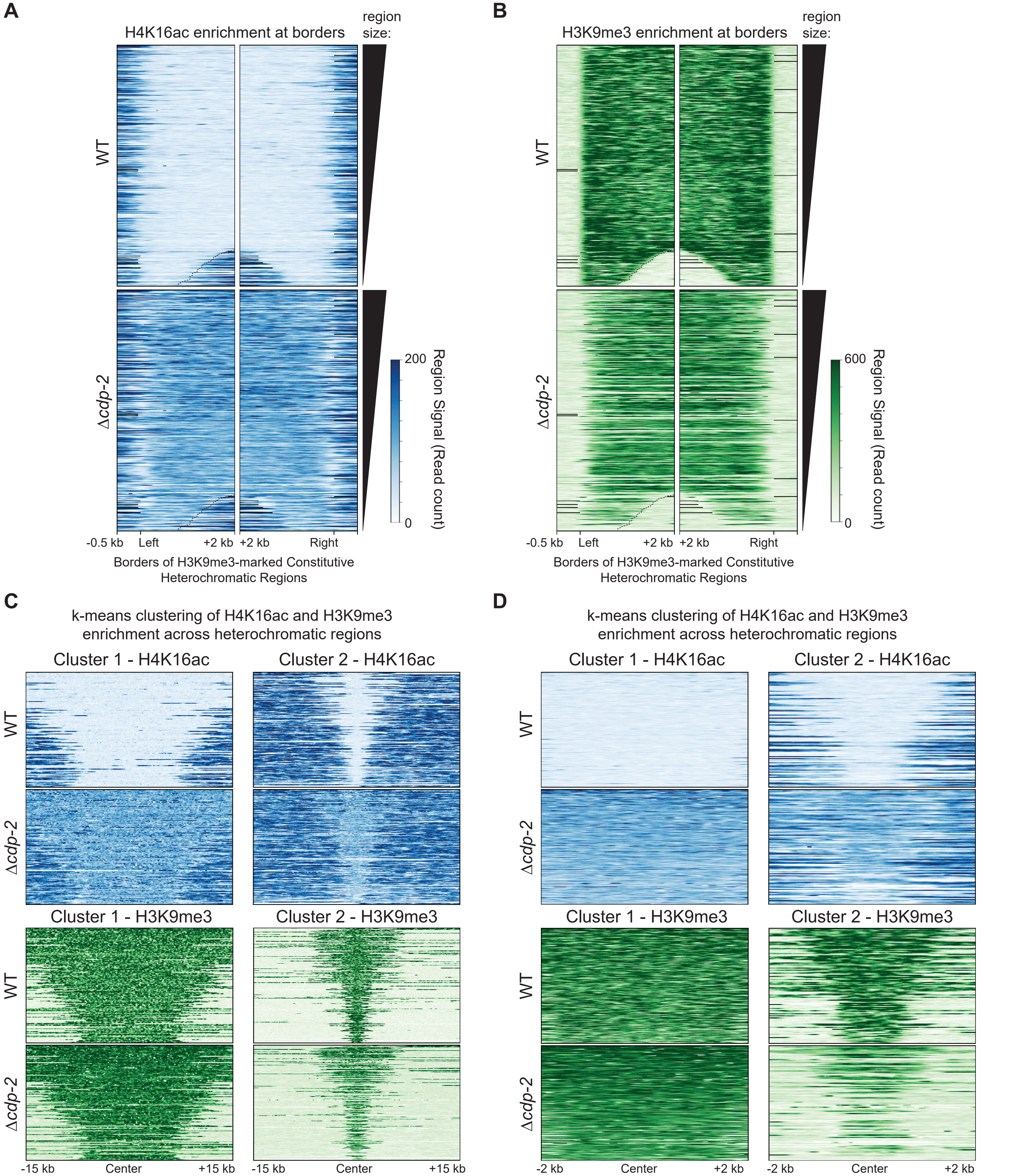


**Figure S20. Borders of heterochromatic regions have unchanged H4K16ac but lose H3K9me3 and k-means clustering of H4K16ac and H3K9me3 enrichment over constitutive heterochromatic regions shows regions losing H3K9me3 still are acetylated at H4K16 in a Δ*cdp-2* strain.** (A-B) Average enrichment heatmaps of the (A) H4K16ac or (B) H3K9me3 enrichment in WT and Δ*cdp-2* strains over the borders of heterochromatic regions. Heatmaps show 500 bp before and 2000 bp after borders, and display regions sorted by size. (C-D) K-means=2 clustering of the center of heterochromatic regions and extending (C) +/- 15 kb or (D) +/- 2 kb for H4K16ac (top) or H3K9me3 (bottom) signal in WT and Δ*cdp-2* stains.
