## Supplementary material for "Histone deacetylation and cytosine methylation compartmentalize heterochromatic regions in the genome organization of *Neurospora crassa*": Scadden_Supplementary-Fig_S21

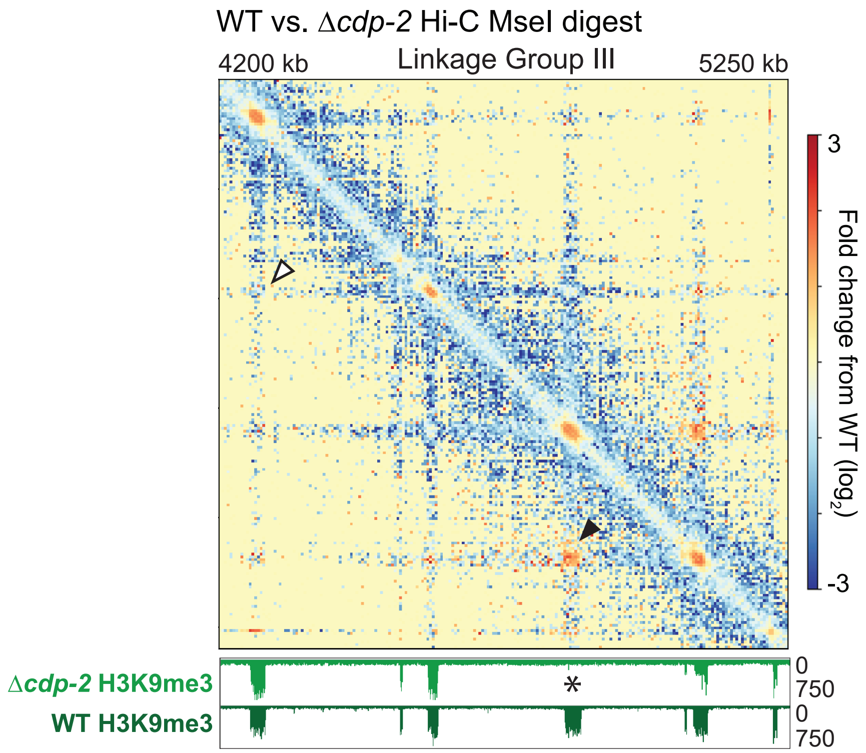


**Figure S21. Constitutive heterochromatic regions that lose H3K9me3 upon CDP-2 loss gain inter-heterochromatin contacts.** Heatmap of the change in contact probabilities between a Δ*cdp-2* strain and a WT strain of a genomic region on LG III containing at least one AT-rich locus that loses H3K9me3 in a Δ*cdp-2* strain. WT and Δ*cdp-2* H3K9me3 ChIP-seq track images below the Hi-C heatmap. Asterisk indicate the region that loses H3K9me3, while black arrowheads highlight gains in inter-heterochromatic region contacts in a Δ*cdp-2* strain; white arrowhead shows an inter-heterochromatic interaction that does not gain interactions despite these silent regions being similarly spaced.
