## Supplementary material for "Histone deacetylation and cytosine methylation compartmentalize heterochromatic regions in the genome organization of *Neurospora crassa*": Scadden_Supplementary-Fig_S22

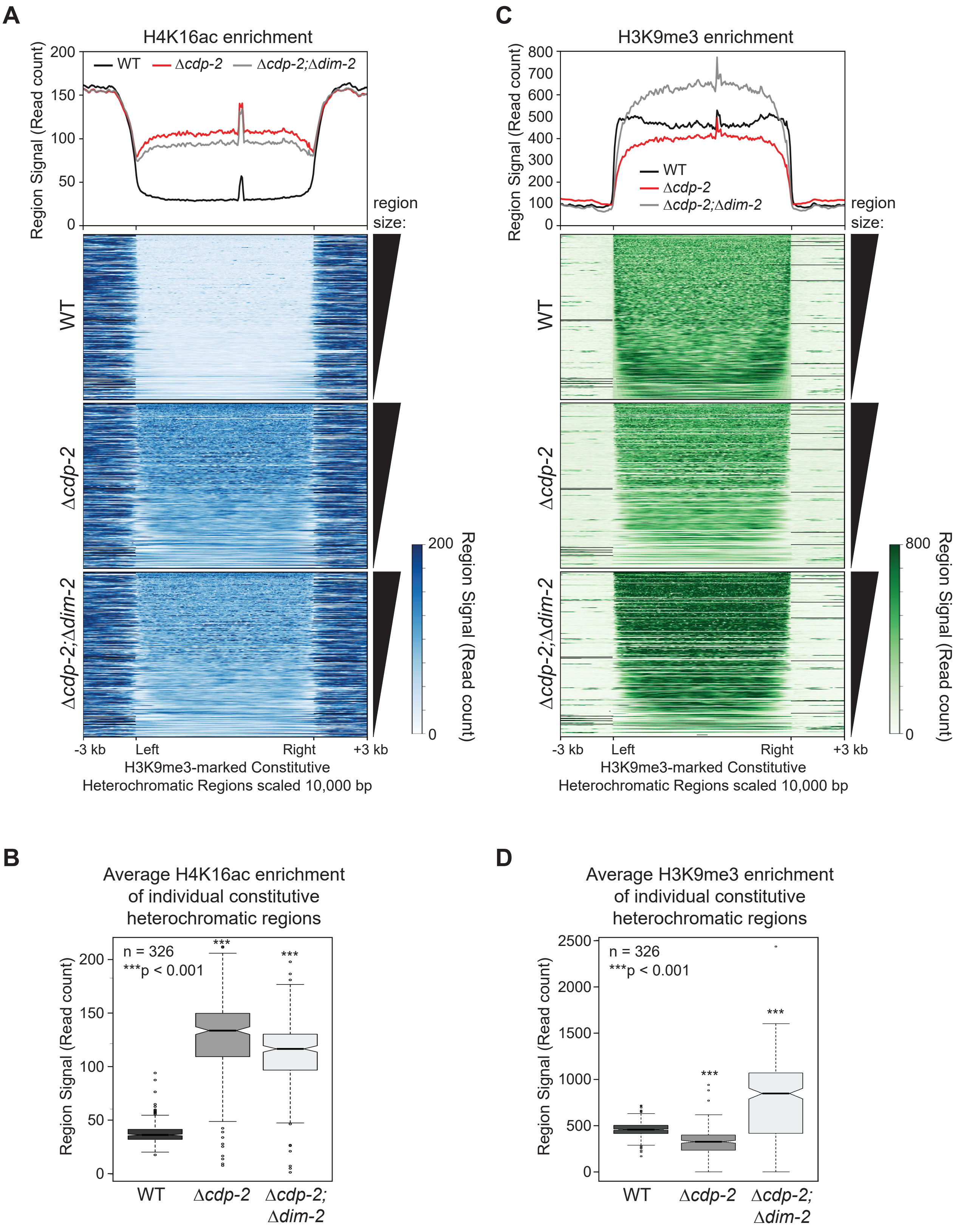


**Figure S22. Loss of CDP-2 and DIM-2 cause gains in H4K16ac and H3K9me3 across constitutive heterochromatic regions in *Neurospora crassa*.** (A, C) Average enrichment profiles (top) and heatmaps (bottom) of the (A) H4K16ac or (C) H3K9me3 enrichment in WT, Δ*cdp-2,* and Δ*cdp-2*;Δ*dim-2* stains over heterochromatic regions (scaled to 10 kb in length). Both heatmaps display regions sorted by size. (B, D) Boxplots of the normalized (B) H4K16ac or (D) H3K9me3 signal (see Materials and Methods) of each heterochromatic region in WT, Δ*cdp-2*, and Δ*cdp-2*;Δ*dim-2* stains.
