## Supplementary material for "Histone deacetylation and cytosine methylation compartmentalize heterochromatic regions in the genome organization of *Neurospora crassa*": Scadden_Supplementary-Fig_S23

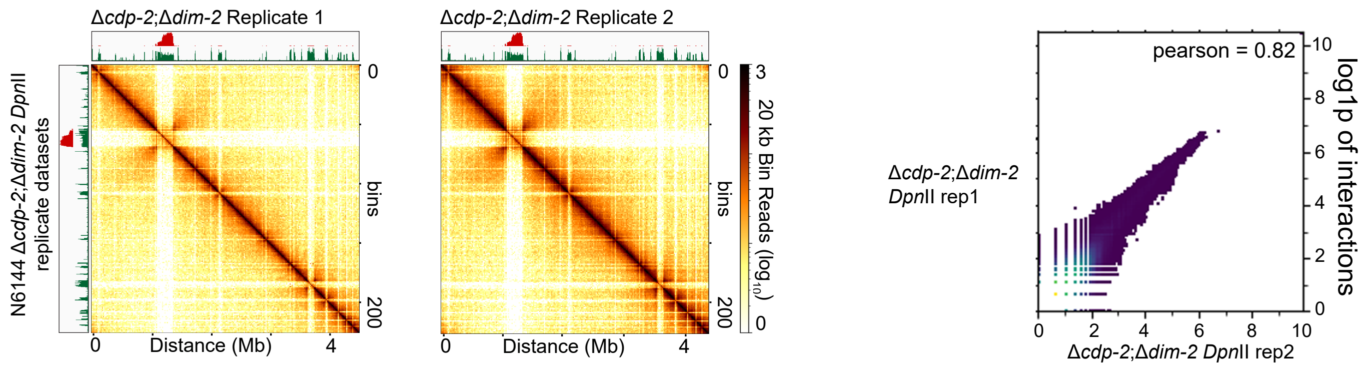


**Figure S23. Replicate analysis of *in situ* Hi-C genome organization datasets, using the *Dpn*II restriction enzyme, of the Δ*cdp-2*;Δ*dim-2* strain of *Neurospora crassa*.** Heatmaps and scatterplots of Δ*cdp-2;*Δ*dim-2* replicates digested with *Dpn*II. In each panel, the left images show the raw count Hi-C heatmaps of genomic interactions across LG II of each *Dpn*II-derived *in situ* Hi-C replicate dataset at 20 kb bin resolution. The right image shows a scatter plot comparing interactions between each Δ*cdp-2;*Δ*dim-2* DpnII replicate matrix at 20kb bin resolution; log1p values of interactions presented. Pearson correlation values of each binary comparison shown at the upper right. Image produced by the hicCorrelate program in hicExplorer.
