## Supplementary material for "Histone deacetylation and cytosine methylation compartmentalize heterochromatic regions in the genome organization of *Neurospora crassa*": Scadden_Supplementary-Fig_S24

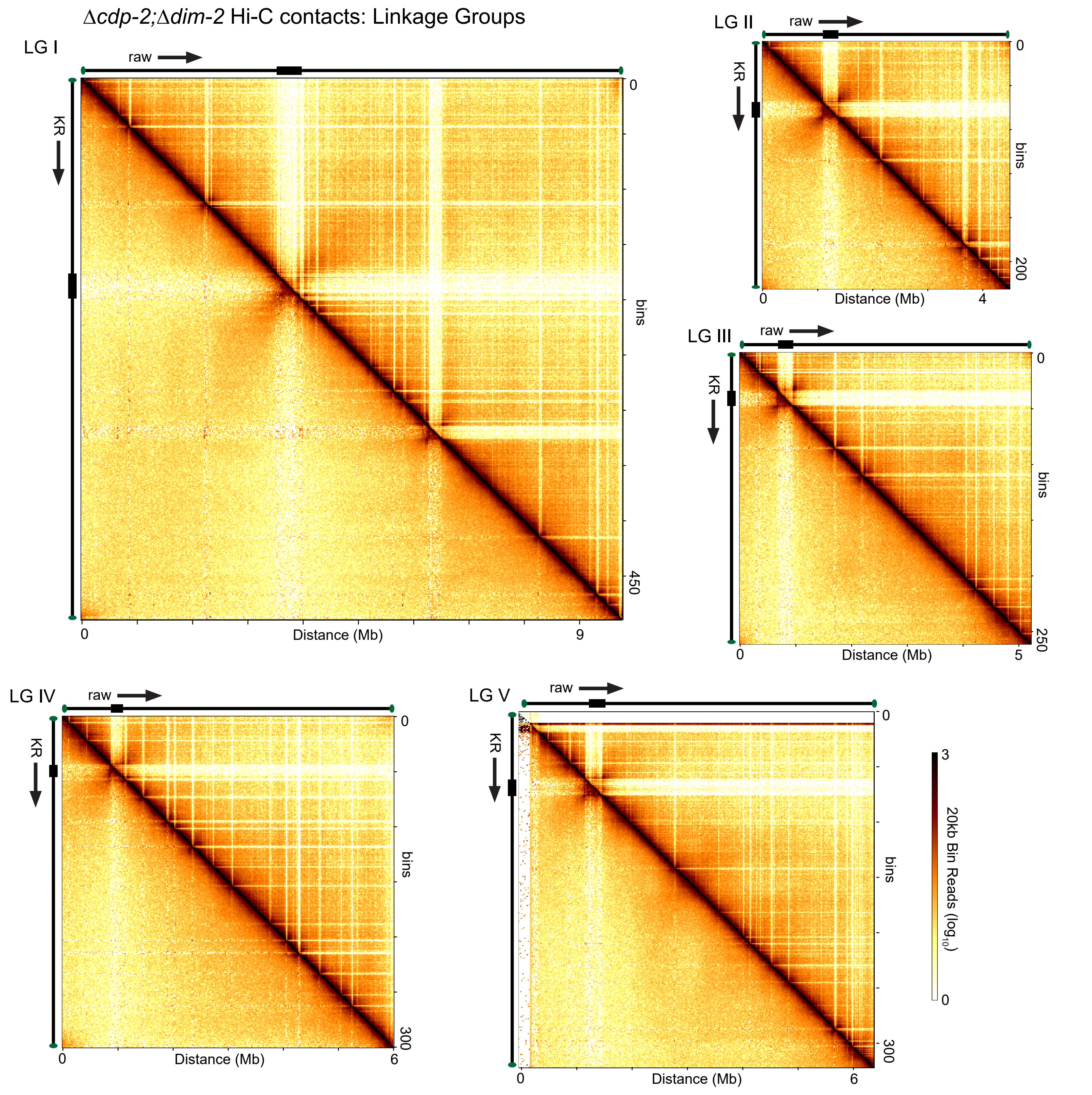


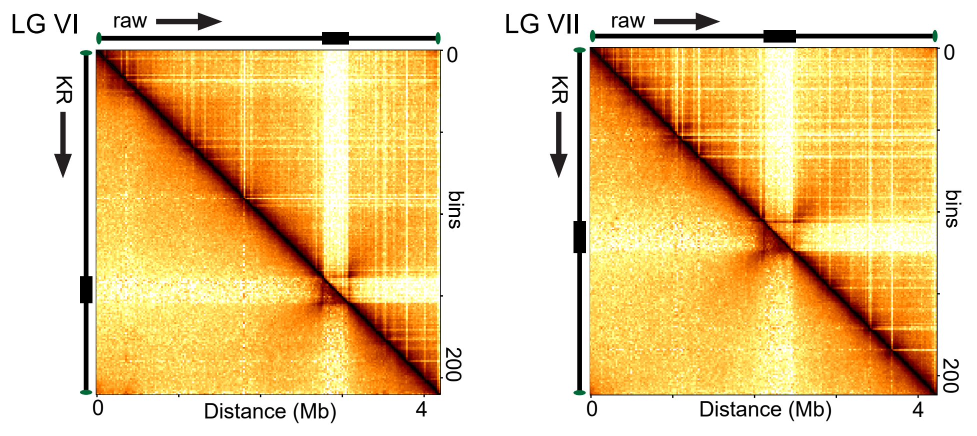


**Figure S24. Chromosome conformation of the seven individual chromosomes of the Δ*cdp-2*;Δ*dim-2* strain using the restriction enzyme *Dpn*II.** Heatmaps displaying *in situ* Hi-C contacts generated with *Dpn*II, either as raw read counts (above diagonal) or KR-corrected counts (below diagonal), for the Δ*cdp-2*;Δ*dim-2* strain across the seven Linkage Groups (LG) at 20 kb bins. Chromosomes are sized according to their length.
