## Supplementary material for "Histone deacetylation and cytosine methylation compartmentalize heterochromatic regions in the genome organization of *Neurospora crassa*": Scadden_Supplementary-Fig_S25

**A**

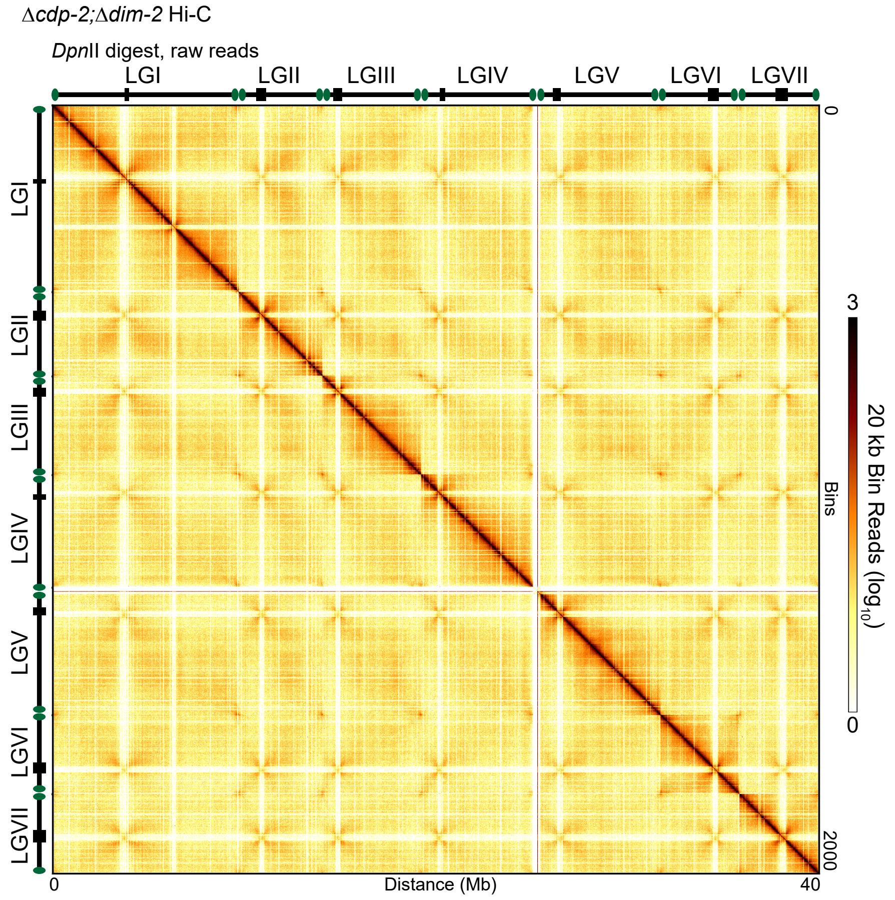


**B**

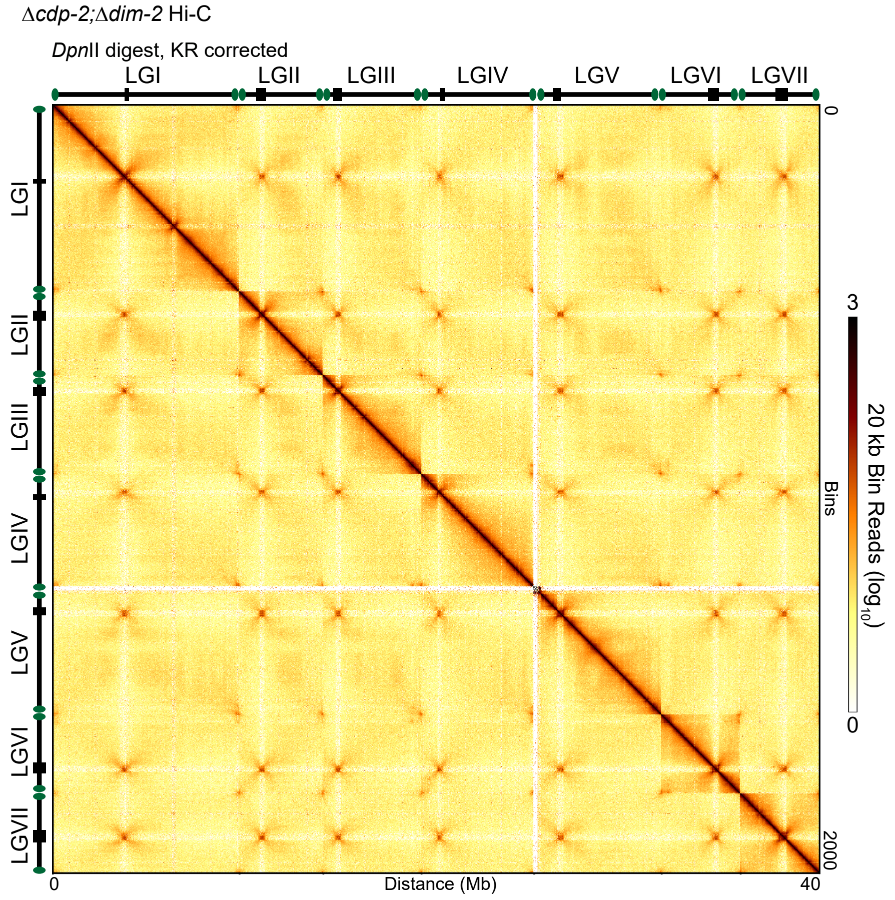


**Figure S25. Chromosome conformation of the whole genome of the Δ*cdp-2*;Δ*dim-2* strain using the restriction enzyme *Dpn*II, either as raw read counts or KR corrected counts.** (A-B) Heatmaps displaying *in situ* Hi-C contacts generated with *Dpn*II, either as (A) raw read counts or (B) KR-corrected counts for the Δ*cdp-2*;Δ*dim-2* strain across the entire *Neurospora crassa* genome at 20 kb bins.
