## Supplementary material for "Histone deacetylation and cytosine methylation compartmentalize heterochromatic regions in the genome organization of *Neurospora crassa*": Scadden_Supplementary-Fig_S28

### Δ*cdp-2* vs. Δ*cdp-2;*Δ*dim-2* - changes in Hi-C contacts: Linkage Groups

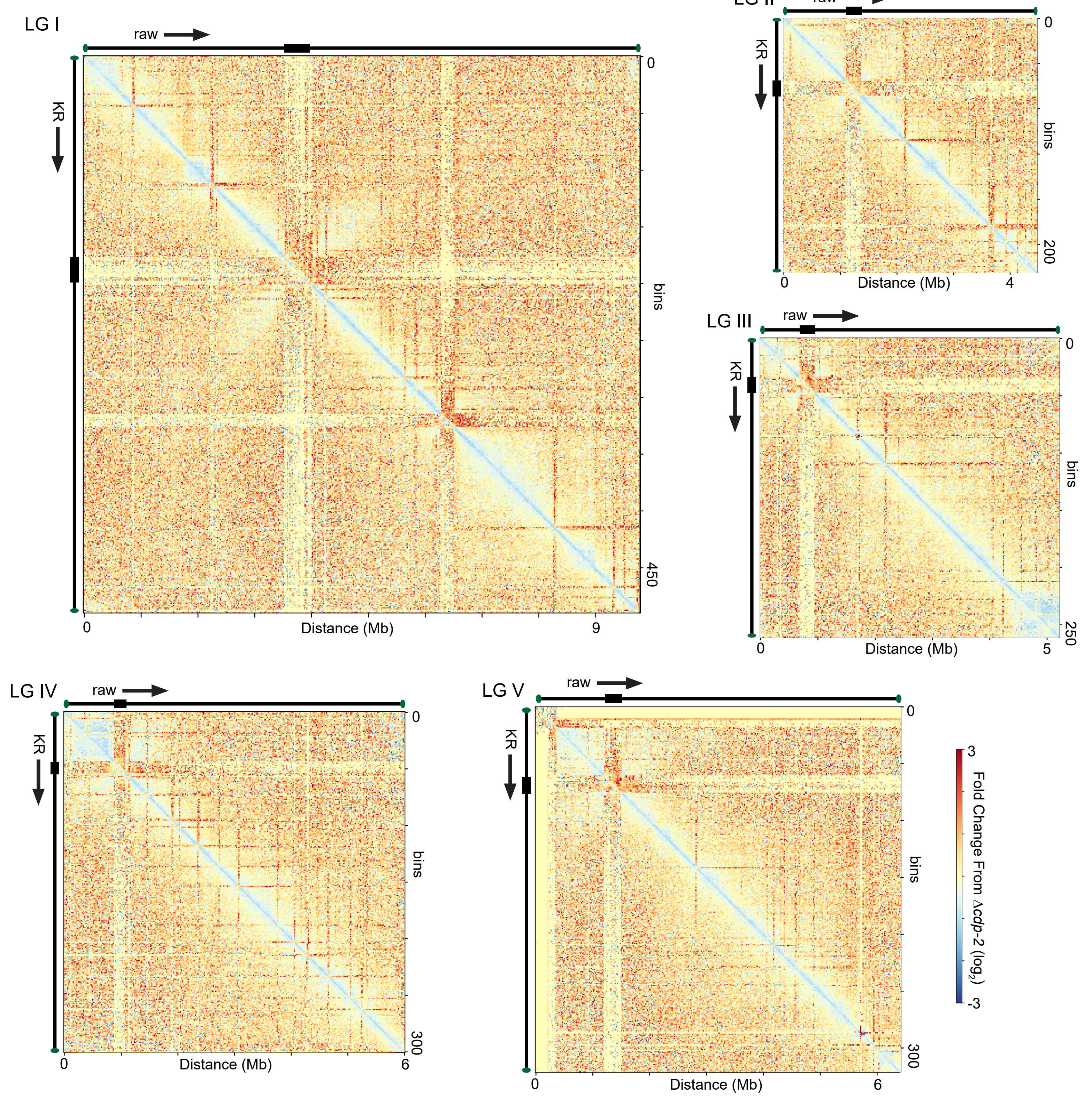


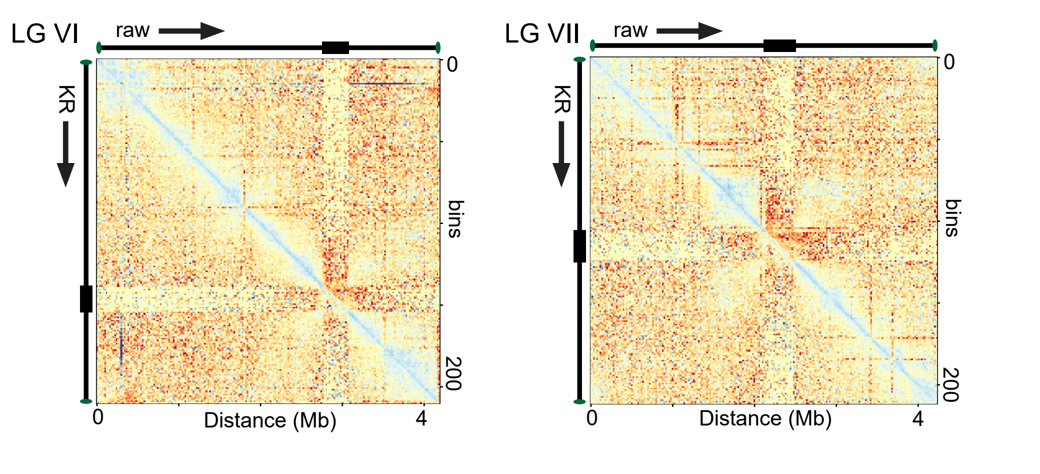


**Figure S28. Changes in chromosome conformation of the seven individual chromosomes between the Δ*cdp-2* and Δ*cdp-2*;Δ*dim-2* strains of Hi-C datasets generated using the restriction enzyme *Dpn*II.** Heatmaps displaying $\log_2$ difference in *Dpn*II generated *in situ* Hi-C contacts, either as raw read count datasets (above diagonal) or the datasets of KR-corrected counts (below diagonal) between the Δ*cdp-2* strain and the Δ*cdp-2*;Δ*dim-2* strain across the seven Linkage Groups (LG) at 20 kb bins. Chromosomes are sized according to their length.
