## Supplementary material for "Histone deacetylation and cytosine methylation compartmentalize heterochromatic regions in the genome organization of *Neurospora crassa*": Scadden_Supplementary-Fig_S29

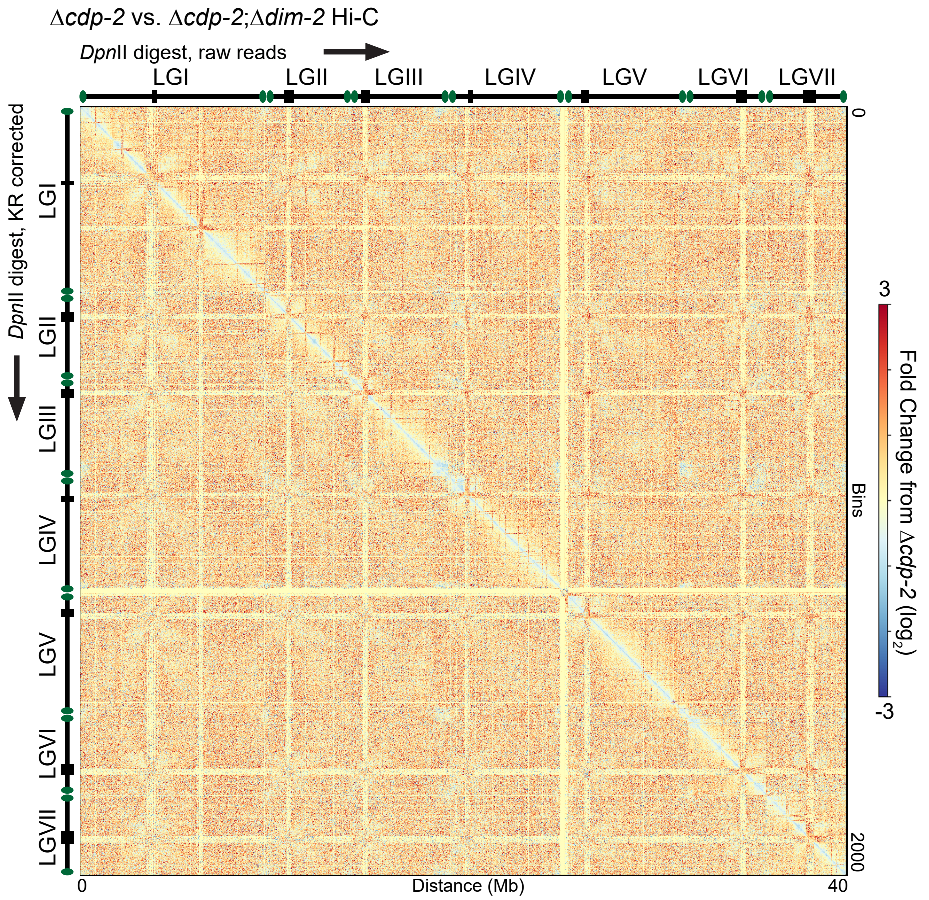


**Figure S29. Changes in chromosome conformation of the entire *Neurospora crassa* genome between the Δ*cdp-2* and Δ*cdp-2*;Δ*dim-2* strains of Hi-C datasets generated using the restriction enzyme *Dpn*II.** Heatmap displaying the changes in *Dpn*II generated *in situ* Hi-C contacts, either as raw read counts (above diagonal) or KR-corrected counts (below diagonal) between the Δ*cdp-2* strain and the Δ*cdp-2*;Δ*dim-2* strain across the entire *Neurospora crassa* genome at 20 kb bins.
