## Supplementary material for "Histone deacetylation and cytosine methylation compartmentalize heterochromatic regions in the genome organization of *Neurospora crassa*": Scadden_Supplementary-File_S1

Scadden et al.

### Materials and Methods

*Strains and growth conditions*

*Neurospora crassa* strains WT N150 or N3752 (both strains are independent propagate strains of 74-OR23-IVA [FGSC #2489]), N3767 (*mat a; his-3 Δcdp-2::hph*), N3613 (*mat A; Δchap::hph*), and N6144 (mat ?; Δ*cdp-2::hph*; Δ*dim-2::hph*) were a gift from Eric U. Selker (University of Oregon). All strains were grown under standard conditions (e.g., 1x Vogels + sucrose media, with necessary supplements) (1).

*Hi-C library construction*

*In situ* Hi-C (2, 3) libraries, which capture ligation products in the nucleus, were constructed as previously described (4), using either the restriction enzyme *Dpn*II (^GATC) to mainly assess euchromatic contacts or the restriction enzyme *Mse*I (T^TAA) to predominantly capture heterochromatic contacts. The entire *in situ* Hi-C protocol adapted for *Neurospora crassa* by isolating spheroplasts is provided in Supplemental File S1. Briefly, Neurospora cultures were grown for four hours at 32°C, crosslinked with formaldehyde and quenched with tris, and treated with a beta-glucanase (Vinotaste) to form spheroplasts. For Hi-C library construction, crosslinked spheroplasts containing 3.5 μg of DNA were disrupted by bead beating (using 150-212 μm Acid Washed Glass Beads [Sigma Aldrich, # G1145-10G]) and the isolated nuclei were made porous by treatment of SDS at 62°C for seven minutes. Nuclear chromatin was digested with the appropriate restriction enzyme (*Dpn*II or *Mse*I; New England Biolabs [NEB]), and overhangs were filled in with Klenow fragment (NEB) using Biotin-14-dATP (Invitrogen, # 19524-016) as well as the standard dTTP, dGTP, and dCTP nucleotides. Blunt-ended fragments were ligated in the nucleus with T4 DNA ligase (NEB), and the resulting DNA loops were purified, Biotin-dATP was removed from unligated ends with T4 DNA polymerase (NEB), and DNA loops were sheared with a Bioruptor Pico (Diagenode). Ligation products were purified with Streptavidin beads (M280 Dynabeads, Invitrogen, # 112.05D), and libraries for Illumina based highthroughput sequencing were constructed with an NEB NEXT Ultra II kit (NEB) per the manufacturer's protocol, except that only eight PCR cycles were used for library amplification to minimize depletion of AT-rich regions in the Neurospora genome (5). Hi-C libraries were sequenced on an Illumina HiSeq 4000 (as 100 nucleotide [nt] paired end reads) or an Illumina NovaSeq 6000 (as 59 nt paired end reads) at the University of Oregon Genomics and Cell Characterization Core Facility.

*Bioinformatic analyses of Hi-C datasets*

All analyses were performed, and all Hi-C images generated, with the HiCExplorer program package (6), as previously reported (4). Previously published *Dpn*II and *Mse*I *in situ* Hi-C data from the WT strain N150 was obtained from the National Center for Biotechnology Information (NCBI) Gene Expression Omnibus (GEO) accession number GSE173593 (4). To normalize WT datasets to compare to mutant strain datasets, the number of total reads were extracted (using the sed command) from the WT R1 and R2 fastq files to provide the number of WT valid reads equal to mutant dataset valid read numbers. Reads were mapped to the previously established nc14 *Neurospora crassa* genome (4) with bowtie2 (7), and used to build the contact matrix with hicBuildMatrix (6); resulting contact matrices were used for all downstream applications, as previously performed (4). Contact quantification was performed by converting hdf5 matrix files to a homer format and counts were extracted into NxN array with the python script dataconvert.py; the python script epigenetic-mark-Quant_v2.py counts intra- and inter-chromosomal bins enriched for specific histone PTMs. Python scripts are available at https://github.com/Klocko-lab/Chip_Quantification).

*Chromatin Immunoprecipitation-sequencing (ChIP-seq) library construction*

ChIP-seq was performed essentially as previously described (8, 9), except that lysing of cells and shearing of chromatin occurred simultaneously with a Bioruptor Pico (Diagenode), using a 15 minute cycle of 30 seconds of sonication and 30 seconds off, in the presence of Halt Protease and Phosphatase Inhibitor Cocktail (Thermo Fisher Scientific, # 78441); protein A/G magnetic agarose beads (Pierce/Thermo Scientific, # PI78609) purified the antibody/chromatin complex; final ChIP-DNA quantification was performed using the Qubit 3.0 HS method; library barcoding was performed using the NEBNext Ultra II barcoding kit for Illumina sequencing (NEB) per the manufacturer's protocols except that eight PCR cycles were used for the final library generation to minimize AT-rich DNA depletion (5). ChIP-seq libraries were sequenced on an Illumina NovaSeq 6000 (Genomics and Cell Characterization Core Facility, University of Oregon). The Klocko Lab ChIP-seq protocol is provided in Supplemental File S2.

*Bioinformatic analyses of ChIP-seq datasets*

Raw ChIP-seq data files, as fastq files, were mapped to version 14 of the *Neurospora crassa* genome (nc14) (4) with bowtie2 (7), and output sam files were converted to sorted bam files using samtools (10), which were used by Deeptools (11) to produce bedgraph and bigwig files, normalized by Reads per Kilobase Per Million Reads (RPKM), for display on the Integrative Genomics Viewer (IGV) (12); IGV images were used for figure creation. Deeptools was also used to create average enrichment signal profiles and heatmaps using bed files for heterochromatic regions (9) or genes. For box plots of the average signal of all features, the average enrichment value per bin for each flanking region was calculated and averaged to obtain a normalization factor to correct the average signal internal to the region; these corrected average signals were plotted in box plots using R and R studio (13, 14). WT H3K9me3 (merged from NCBI GEO accession numbers GSE68897 and GSE98911), WT CenH3 (NCBI GEO accession number GSE71024) and SET-2 H3K27ac (NCBI GEO accession number GSE118495) datasets were previously published (9, 15–17).
