## Supplementary material for "Histone deacetylation and cytosine methylation compartmentalize heterochromatic regions in the genome organization of *Neurospora crassa*": Scadden_Supplementary-File_S2

### Protocol for *in situ* Hi-C.

**Initiated by Jon Galazka, Freitag lab. Modified substantially by A. Klocko, 071019 (updated 040722)**

**Picture of what HiC is:**

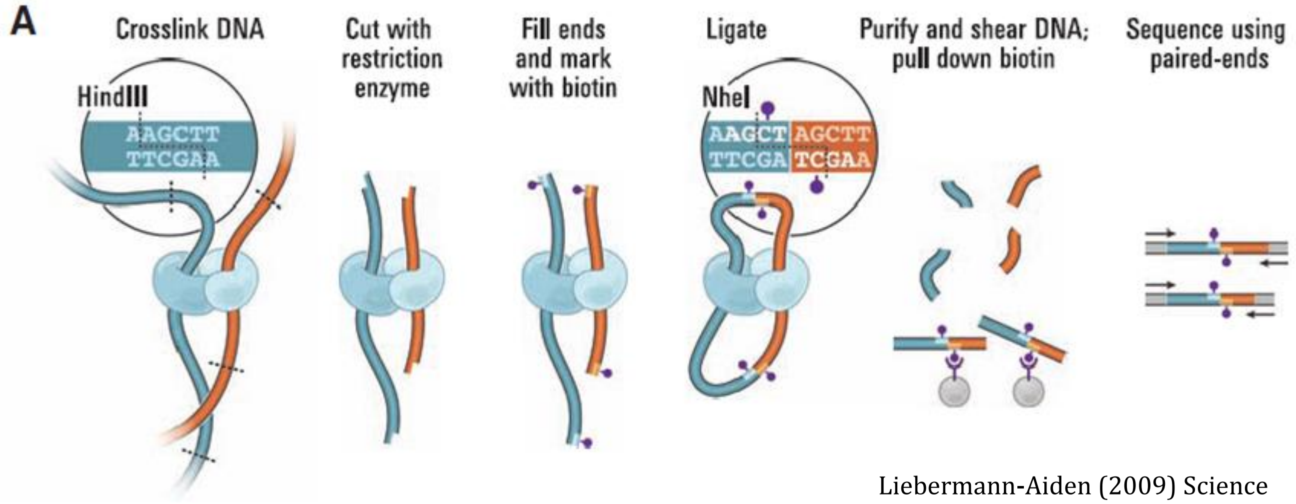


Liebermann-Aiden (2009) Science

**Prepare spheroplasts**

About a week prior, grow six to eight baby slants of the strain(s) on which you wish to perform Hi-C.

*If the strains grow (conidiate) well, this should be enough conidia for two technical replicates (to perform Hi-C in duplicate).*

Day 1:

On the day of culture growth, harvest the conidia from **SIX** (maybe seven) of these strains. To each baby slant:

Add 1mL $dH_2O$, vortex, collect (by decanting or pipetting) into a 15mL conical tube

Add an additional 1mL $dH_2O$ aliquot, vortex, and collect into the same 15mL conical tube

Repeat for the other five to seven slants

Centrifuge 3000 rpm, 1 minute. Discard supernatant (by decanting, ensure there is no loss of material; ok if small amount of liquid remains)

Wash conidia twice with 10mL $dH_2O$

(add 10mL $dH_2O$, resuspend pellet by inverting until pellet breaks up, centrifuge)

Resuspend in 1mL $dH_2O$ and transfer to eppendorf tube.

Count a 1:100 dilution (5µL conidia suspension + 495µL $dH_2O$, vortexed) on hemocytometer (volume of 25 squares = 1 $x10^{-4}$ mL; volume of 5 squares = 0.2 $x10^{-4}$ mL) using the wide-field scope and the 40 x objective by adding 10 µL of 1:100 dilution to the hemocytometer (between the cover slip and the slide).

for math: Count 5 squares, add the conidia numbers up, multiply by the dilution factor (100), and divide this total number of conidia by 0.2 $x10^{-4}$ mL to get the number of conidia per mL in your stock solution. then divide $1.25x10^8$ by the conidia/mL value you just calculated for the volume of your conidia stock to add

Set-up 25 ml culture with 1.25e8 conidia:

Calculate the amount of conidia (of non-diluted sample) to give you $1.25 \times 10^8$ (don't forget the dilution factor!)

Start culture in a 125mL flask with 25mL of solution 1xVogels containing 1.5% sucrose (add amino acids, if needed) and added required conidia suspension.

Incubate at 32°C with shaking 200 RPM for 3-4 hours (four hours preferred)

(after 3 hr, Wt conidia should be 70% germinated, with hyphae extending ~4x the length of the conidia. Mutants can take significantly longer). This growth period yields ~12.5µg of gDNA from fixed germlings.

Cross-link the germlings in each flask by adding 0.7 mL of FRESHLY-OPENED (within last month) formaldehyde (37%) to each flask. This produces a final concentration of 1% for 10 min while shaking at RT, 100 RPM.

Add 3.4 mL of 1.0 M Tris-HCl pH 8.0 (final concentration: 125 mM) for 10 min to quench the reaction (shaking 100 rpm, RT).

Harvest germinated conidia in 1x50mL conical tubes by decanting and centrifuge at 4000 RPM, 2 min. Resuspend in 20 mL Spheroplasting buffer. Add 14.2M βME stock to 30 mM (43.0 µl per tube). Pellet at 4000 RPM, 2 min.

Remove supernatant by pipetting and suspend the pellet in 20 mL Spheroplasting buffer and 1 mM BME (1.4 µl) then pour into 125 mL Erlenmeyer. Add 50 mg VinoTaste (a beta-glucanase, 1 mL from 50 mg/mL stock made in PBS, filtered), incubate at 30 °C for 60 min, shaking at 100 rpm. (store VinoTaste at 4°C for up to one week, make fresh each time!)

Gently harvest by decanting into 50 ml conical tube and pellet at 3500 RPM for 2 min. Wash twice with 20 mL of Spheroplasting buffer. (Note: It is preferred to pipet the supernatant in these steps as decanting may disrupt the pellet)

Wash 3x with 20 mL 1x HindIII/MseI (or 1x DpnII- specific to restriction enzyme you are going to use!) buffer. Pellets should turn pale orange/brown as the cell wall is washed away. Careful, as pellets are slick. Gently resuspend pellet in 3.5mL HindIII/MseI or DpnII buffer, split into 4, 1 mL aliquots (for a total of 4), pellet @ 10k rpm for 1 min, remove majority of supernatant to about 200 µL and centrifuge again before removing remaining liquid.

Snap freeze spheroplasts by storing the tubes at -80 °C.

Continue on with Day 1 protocol here (one last step): Check DNA conc. of spheroplasts

(NOTE: you are sacrificing ONE tube of XL spheroplasts to check the DNA concentration of the culture. Once DNA concentration is found, you discard that sample and get a new tube of spheroplasts for the following steps)

Resuspend 1 pellet in 200 µl TE + 0.5% SDS (10 µL of 10%) + 4 mM EDTA (1.7 µL of 0.5M), and 5 µl proteinaseK (20 mg/ml stock) and resuspend the pellet by pipetting up and down. Put tubes in rack and place in 65°C incubator overnight (decrosslinking the reactions)

Day 2:

Next day, bring volume to 500 µl with TE, add 4 µl RNaseA (10mg/mL stock), vortex, and incubate at 37°C for 30 min.

Extract 1x with 500 µl 25:24:1 phenol:chloroform:IAA to remove cell debris from sample.

- For the extraction: add 500 µl phenol:chloroform:IAA solution, vortex 20 sec, centrifuge 5 min at 13k rpm, remove top layer (aqueous solution) to a new tube

Extract 1x with Chloroform to ensure that no material is visible at the interface.

- Add 500 µl chloroform, vortex 20 sec, centrifuge 5 min at 13k rpm, remove top layer (aqueous solution) to a new tube

Precipitate with 1/10 vol NaAcetate (3M, pH6.0: 50 µL), 1 vol Isopropanol (100%: 500 µL)

Add solutions, invert or vortex tubes, incubate at -80°C for 5-10 min, spin 13.0k rpm for 5 min, discard supernatant.

Wash pellet 1x with 70% ethanol (remove supernatant, flow 1mL of 70% ethanol over sample, quickly remove solution again – no need to resuspend pellet). Dry in speedvac 10min @ RT, and resuspend in 50 µl TE.

Check concentration by Qubit HS – use 1 µL of the sample. Should be ~2.5 to 5 µg in each tube of crosslinked spheroplasts.

*Optional: electrophorese 15 µl out on gel + 3 µL of dye (0.8% TBE or TAE should be fine)*

**Digest with DpnII or MseI** (or HindIII, if your spheroplasts have been concentrated in that buffer)

Note: now that you have the DNA concentration of one pellet of spheroplasts, you throw out what you just worked on, and use NEW spheroplast tubes (stored at -80°C).

Calculate the amount of spheroplasts needed for 3.5 µg gDNA (If you resuspend a spheroplast pellet in one tube in 1x DpnII buffer up to 100 µL, and it's gDNA concentration was 75.87ng/µL in 50 µL TE [above], you'd need 94.9 µL of spheroplasts).

*Optional: if you want to use a different enzyme relative to what buffer you used to wash your conidia, do a buffer exchange:*

- *Resuspend your conidia pellet in 1mL of a 1x concentration of your new buffer.*
- *Centrifuge 30 seconds, 3000 rpm (you should have a nice pellet here). Remove supernatant*
- *Repeat for a total of three times*

Suspend spheroplasts containing 3.5 µg gDNA in 270 µl 1x DpnII (or 1x HindIII/MseI) digestion buffer. Keep on ice.

Add 50 µL of hydrated Glass Beads (Sigma-Aldrich # G1145, Acid washed, 150-212 µm, resuspended in $dH_2O$) to the 270 µL chromatin.
Vortex 5 minutes: 30 sec on, 30 sec on ice.
Let beads settle and move 270 µL of supernatant (as well as the broken-apart debris) to a new ***screw top 2 mL*** tube.

Add 30 µl of 6.25% SDS (to 0.625%), incubate at 62°C (+/- 3°C) for 7 min. Put immediately on ice.

(NOTE: this step gets rid of transient XLs, inactivates nucleases, and makes the nuclear membrane porous)

Place on ice and add 33 µl 10% Triton X-100 (to 1%) and mix by pipetting. (NOTE: Triton x100 binds the SDS, inactivates it)

Add 8.3 µl of 10x DpnII or 10x HindIII/MseI buffer (to account for volume of SDS, Triton and Enzyme).

Add 200 units DpnII (or MseI), which is 20 µL of 10 U/µL, ~~(or HindIII: 10 µl of 20 U/µl, NEB)~~. Parafilm tube to seal solution in.

Digest for at least 16 hr with tumbling (or on a nutator) at 37°C. Put water bath into cold room and set to 16°C.

Day 3:
**Perform HiC with biotinylated nucleotides**

Centrifuge 3000rpm, 10 minutes.
Remove supernatant and gently resuspend pellet in 83 µL 1x DpnII or HindIII/MseI buffer.

To the resuspended pellet sample, add:
- 2.1 µl of dCTP/dGTP/dTTP mixture aliquot (each nucleotide at 4mM concentration)*; discard aliquot after use*
- 7.0 µl of 0.4 mM Biotin-14-dATP (~1/18 of 125 µl tube from Invitrogen, cat# 19524-016))
- Add 1.5 µl NEB 10x DpnII buffer (or Buffer 2.1 for HindIII/MseI) *(buffer choice dependent on enzyme used!)*
- Add 1.4 µl water
- Add 5.0 µl of Klenow (large fragment) at 5 U/µl (25 U)
  - Mix by pipetting with a P200; 100 µl final volume

*NOTE for AK: use Biotin-dATP for either enzyme: MseI (5' T^TAA) ~~HindIII, (5' A^AGCTT)~~, or DpnII (5' ^GATC) should be compatible with Biotin-dATP. No real reduced ligation efficiency observed with MseI. We will try using 50% less Biotin-dATP too (7.0uL).* For biotin HiC library, add each dNTP to 30 µM: (NOTE: NEB website says reactions need to have 30µM of each dNTP for blunting- PERHAPS USE LESS VOLUME FOR THIS REACTION, SO WE CAN ADD LESS BIOTIN dATP! This blunting reaction fills in restriction enzyme overhangs with NTPs, including biotin-dATP. Math below, adding 0.7µL to a 100 µL reaction gives ~28 µM – not exact, but it works.)

Incubate at 37°C with tumbling for 60 min. Place on ice immediately.
Add 110 µL 1x DpnII or 1x MseI/HindIII buffer.

Prepare 420 µL (total volume) ligation reactions with T4 DNA ligase (NEB).
*This ligation is at a smaller volume for an in situ Hi-C experiment (Rao et al., 2014, Belaghzal et al., 2017); we removed the previous SDS digestion per advice of Belaghzal et al.!*
**NOTE: add ingredients in THIS order (from top to bottom).**

To the **210 µl of chromatin** (~3.5 µg, conc. of ~8.4 µg/ml), carried over in the same tube from the previous step, add:
- 117.6 µl water
- 42 µl 10x T4 DNA Ligation Buffer *(use the 10x T4 DNA Ligase Buffer we get from NEB that comes with the ligase enzyme)*
- 42 µl 10% Triton X-100 (1% final)
- 4.2 µl 10 mg/ml BSA (100 µg/mL final)
- 4.2 µl 400 U/µL NEB T4 DNA ligase (1675 NEB units, 25 Invitrogen units)
  - Total volume is 420 µL; MIX by gently pipetting. Ligate at 16°C for **4 hours**.

Add 12.5 µl of proteinaseK (20mg/mL stock), mix, and decrosslink overnight at 65°C.
Turn down water bath in cold room to 12°C

Day 4:
**Purify 3C library**

Add another 12.5 µl of proteinaseK and decrosslink for another 2 hours at 65°C. Cool to RT. Final volume here is 445µL
In downtime, label (with your strain #s) two sets of 1.7mL Eppendorf tubes and two sets of 2.0mL microcentrifuge tubes for following steps

Add 450µL of saturated phenol:chloroform:IAA (pH 8.0) and vortex for 20 sec. Spin for 5 min at 13000 rpm.

Transfer supernatant to new eppie tube and extract with 450 µl phenol:chloroform:IAA (pH 8.0), vortexing 20 sec.

Spin for another 5 min at 13000rpm.

Transfer supernatant to a 2.0mL microcentrifuge tube and add TE to bring the final volume to ~500 µL

Add 50 µL (0.1x volume) of 3M Na Acetate pH 6.0 AND 2 µl glycogen (20mg/mL) and mix well by vortexing.

Add 1250 µl (2.5x voume) of 100% ethanol (or 1320 µl of 95% EtOH) and mix by inversion.

Incubate at -80°C for 30 minutes. Spin for 15 min at 13,000 rpm (16k x g).

Discard supernatant by pipetting and wash once with 1 mL of 70% ethanol. Centrifuge 13,000 rpm for 2 min.

Discard supernatant and speedvac pellets for 10 min at RT. Pellets will look "glassy".

Resuspend the pellet in 50 μl of TE by pipetting.

Add 2 μl RNaseA (10mg/mL stock) and incubate at 37°C for 30 min. Add 450 μL TE (500 μL total)

Extraction 1x with 500 μL Phenol:chloroform:IAA and 1x with 500 μL chloroform:
[For each extraction: Add organic solvent, vortex for 20 seconds, centrifuge for 5 min (13k rpm), remove aqueous (top) layer to new tube.]

Transfer supernatant (~500 μL) to new 2.0mL microfuge tube, add 1/10 vol. 3 M NaAcetate pH 6.0 (50 μL) and vortex to mix.

Add 2.5 volume (1250 μL) of 100% ethanol (or 1320 μl of 95% EtOH) and invert tubes.

Incubate at -80°C for 20 min.

Spin at 13,000 rpm (16k x g) for 10 min in microfuge.

Discard supernatant and wash once with 1 mL of 70% ethanol.

Centrifuge 13,000 rpm for 3 min.

Discard supernatant and speedvac pellets for 10 min at RT. Pellets will look "glassy" but be smaller than 1st precipitation.
CAREFUL: pellets may be so dry that they fly away from the edges of the tubes. Close tubes immediately after speedvac finishes (while tubes are still in rotor) so pellets aren't lost.

Resuspend in 25 μl of TE/10 by pipetting; solution should look "gooey" and should "stick" to the walls of the tubes a bit.
Store frozen at -20°C.

Day 5:
**Shear Hi-C library**

Remove Biotin-dATP from unligated ends:
To the 25 μl DNA (from the previous step), add:
17.0 μl water
5 μl NEB buffer 2.1
1.0 μl dATP/dGTP mixture aliquot (each nucleotide at 10mM concentration)*; discard aliquot after use*
1.7 μl (5 U) of T4 DNA polymerase
final volume of 50 μl.
Incubate at 12 C for 2 hours.

Quench by adding 1 μl of 0.5 M EDTA. Add an additional 450 μL TE/10.

Extraction 1x with 500 μL Phenol:chloroform:IAA and 1x with 500 μL chloroform:
[For each extraction: Add organic solvent, vortex for 20 seconds, centrifuge for 5 min (13k rpm), remove aqueous (top) layer to new tube.]

Transfer supernatant (~500 μL) to new 2.0mL microfuge tube, add 50 μL NaAcetate (3M, pH 6.0) and vortex to mix.
Add 2.5 volume (1250 μL) of 100% ethanol (or 1320 μl of 95% EtOH) and invert tubes.
Incubate at -80°C for 20 min. Centrifuge 14000rpm 10min (*Turn on Bioruptor here).*
Wash 1x with 1 mL of 70% EtOH [add 70% EtOH, quick spin and remove supernatant]
Dry 10 min speedvac (close tube lids immediate after the spin is done to prevent pellet loss, as pellets will be loose).
Resuspend in 250 μl of TE/10 (vortex), and move to a bioruptor tube (clear polystyrene tube).

Break apart DNA ligation loops with the Bioruptor in CENT278.
*NOTE: this step is to break apart the DNA ligation "loops" to make DNA libraries capable of being prepped for HT-seq*
Add six tubes to the Bioruptor (with blanks, if needed) and run the protocol called "15min-30sec-ON":
The protocol is 15-minute total: 30 seconds on, 30 seconds off

Bind to Streptavidin beads (Invitrogen, Dynabeads M280, Cat# 112.05D) – Wash beads:
Pipette (25 μl x number of samples) of bead slurry into a new Eppendorf tube:
Bind to magnetic rack and remove supernatant
Add 1mL of 2xBW buffer (10 mM Tris-HCl pH 8.0, 1 mM EDTA, 2 M NaCl) and resuspend beads
Bind to magnetic rack, remove supernatant
After three total washes, resuspend beads in 50 μl x the number of samples with 2xBW buffer.
Add 50 μL of washed beads and 200 μl additional 2xBW buffer to sheared DNA (as long as ratio of washed beads: HiC library is 1:1, it can be ANY volume).

Bind for 30 min at RT, mixing by pipetting every 10 min.

Wash beads (on magnetic rack) TWICE with 1000 µL 1xBW buffer, ONCE with 200 µL 1xBW buffer, TWICE with 200 µL TE/10.
For washes: place tubes in rack, let beads be drawn to magnet side, aspirate away supernatant, remove tube from rack, and add buffer – pipetting up and down to resuspend beads before adding the tube back to the rack.

Resuspend in 25 µl TE/10; move to a new 1.7mL eppie tube. Store at -20°C

Day 6:

**Prepare Illumina library (Klocko lab protocol)** NOTE: this protocol below uses the NEBNext Ultra II kit from NEB.

END PREP, in a 0.2mL PCR tube (in this order)

- 30 µL $dH_2O$
- 7 µL NEBNext Ultra II End Repair reaction buffer (green top)
- 3 µL NEBNext End Repair Enzyme Mix (green top)
- 20 µL HiC library on streptavidin beads (using most of the reaction to have more final DNA product after 8 cycles)
  - Mix the reaction well by pipetting when you add the beads

Use program CHIP20 on the PCR machine, with heated lid (99°C):

CHIP20:

- 20°C for 30 minutes
- 65°C for 30 minutes
- 4°C HOLD

Ligate hairpin adapter

To the 60 µL end prep reaction (previous step), directly add these reagents:

- 1 µL NEBNext Ligation Enhancer (red top)
- 2.5 µL NEBNext Adaptor for Illumina (red top)
- 30 µL NEBNext Ultra II Ligation Master Mix (red top)
  - Mix reaction well by pipetting

Use the program CHIP20-LIG

CHIP20-LIG: 20°C, with NO heated lid

Ligate Adapters in the PCR machine for **15 minutes**!

Add 3 µL of USER Enzyme (red top) to the ligation mixture. Mix well by pipetting.

Use program CHIP37 - incubate at 37°C for 15 minutes (with heated lid at 99°C)

Wash beads 5x with 200 µL 1x BW buffer (move to a new 0.2mL PCR tube after the second wash),
wash 1x more with 200 µL TE/10
Resuspend beads in 15 µL TE/10

Enrich DNA fragments

In the same 0.2mL PCR tube (with the 15 µL HiC DNA/streptavidin bead mixture – use all the reaction here), add:

- 5 µL Universal PCR primer
- 5 µL Index Primer (USE A DIFFERENT INDEX PRIMER PER REACTION; THIS ADDS THE UNIQUE BARCODE)
- 25 µL NEBNext Ultra II Q5 Master Mix

Mix reaction, place into PCR machine, using CHIPAMP protocol: RECORD ADAPTERS USED FOR EACH SAMPLE HERE!

- 1 cycle 98°C 30 seconds
- 8 cycles of:
  - 98°C, 10 sec
  - 65°C, 75 sec

**DO EIGHT PCR CYCLES TO MINIMIZE AT-RICH REGION DEPLETION!!!!!**

- 1 cycle 65°C, 5 minutes
- hold at 4°C

Place on magnetic rack. Remove the supernatant (which is 50 µL) to a NEW 0.2mL PCR tube

Ampure XP (Agencourt) purify the PCR reaction

- Mix 50 µL Ampure XP beads (warmed to RT) and the 50 µL PCR reaction
  - (no need for extra PEG8000/NaCl, as we want to get rid of fragments below 200bp, which are primer dimers)
- Bind DNA to beads for 15 minutes at RT

Put on magnetic rack for 5 min (Ampure beads are magnetic and will go to the magnet side of the tube)
Remove and discard supernatant (DNA is on Ampure beads)
wash 2x with 200 µL FRESH 80% EtOH (just flow EtOH over the beads – do NOT resuspend beads)
remove last of EtOH with P20 pipette
Dry 5 min at RT
Take off magnetic rack
Add 25 µL TE/10 to elute DNA, resuspend beads by pipetting (may need to use some force)
Keep tube at RT for 2 minutes on a NON-magnetic rack
Place tubes back on magnetic rack. wait 1 minute for beads to be drawn to one side
Save supernatant (25 µL) with HiC library to NEW 1.7mL eppie tubes. Store at -20$^{o}$C.

****Normally we'd run a gel to check the libraries, but with 8 PCR cycles we actually get less material back out. So by running a gel, we'd lose critical sample we may want to sequence.**
Typical concentrations of libraries after 8 PCR cycles are around 0.1 to 1.0 ng/µL. As long as we get about this much DNA back out, we can assume we have an appropriately barcoded library for sequencing

Check concentration by Qubit HS (use new standards and 2 µL per sample for Qubiting).
Pool for seq

For HT-seq at the UO core, sample should be a minimum of 2nM in 20 µL, **although its best to submit around 5nM to 10nM in 50 µL**. Use TE/10 for final volume. Calculate volumes to pool in Excel sheet.

At the UO core, we will purchase 100nt PE reads, which provide the maximum amount of data (and are the cheapest). We can do 75nt or 150nt PE reads, but they are either more expensive and the data is not needed (150nt) or rarer to be sequenced (75nt – not many people request these, so we'd wait longer). We HAVE to do PE reads – absolutely required!!!

PAIRED ENDS ARE CRITICAL… each ligation product will have two sides of an intra/inter-chromosomal interaction, where each side of the ligation product (flanking a DpnII/MseI site) must be mapped to the Neurospora genome

When loading the lane into iLab, have Brett generate a PO number, input the sample names and barcodes into the form, and select the i7 side as the location of the barcode!! Andy will do any lane ordering.

For data analysis, use the hicExplorer program (separate protocol) on the Bioinformatics computers (Thing 1, Thing 2, or Horton).

NEBNext Barcodes

| Kit #7335L | Kit #7500L |
|---|---|
| Index 1: ATCACG | Index 13: AGTCAA |
| Index 2: CGATGT | Index 14: AGTTCC |
| Index 3: TTAGGC | Index 15: ATGTCA |
| Index 4: TGACCA | Index 16: CCGTCC |
| Index 5: ACAGTG | Index 18: GTCCGC |
| Index 6: GCCAAT | Index 19: GTGAAA |
| Index 7: CAGATC | Index 20: GTGGCC |
| Index 8: ACTTGA | Index 21: GTTTCG |
| Index 9: GATCAG | Index 22: CGTACG |
| Index 10: TAGCTT | Index 23: GAGTGG |
| Index 11: GGCTAC | Index 25: ACTGAT |
| Index 12: CTTGTA | Index 27: ATTCCT |

**Buffers and Enzymes**

10x HindIII/MseI Buffer (need ~15mL of 10x per culture)
*(this buffer is equivalent to NEB Buffer 2.1)*
*MseI has 100% activity in NEB Buffer 2.1*
500 mM NaCl
100 mM Tris-HCl pH 7.9 @ 25
100 mM $MgCl_2$
Dilute to 1x with water and add BSA to 100 µg/mL.

10X DpnII Buffer (need ~15mL of 10x per culture)
*(this buffer is equivalent to NEB Buffer 3.1)*
1000 mM NaCl
500 mM Tris HCl pH 7.9 @ 25°C
100 mM $MgCl_2$
Dilute to 1x with water
add BSA to 100 µg/mL

For 50mL of 10x DpnII
10mL of 5M NaCl
25mL of 1M Tris HCl
5mL of 1M $MgCl_2$
10mL $dH_2O$
filter sterilize

For 10mL of **1x** DpnII or HindIII/MseI Buffer
1mL of 10X Buffer
10 µL of BSA stock (100mg/mL)
9mL $dH_2O$
filter sterilize

Spheroplasting buffer - 500 ml
91 g Sorbitol (1 M)
43.3 ml $K_2HPO_4$ (Dibasic, 1M stock solution)
6.7 ml $KH_2PO_4$ (Monobasic, 1M stock solution)
$dH_2O$ to 500mL
pH to 7.5 with KOH or Phosphoric acid
Autoclave

TE/10
10mM Tris-HCl, pH 8.0
0.1mM EDTA
Filter sterilize or autoclave
(Make fresh every time!!)

For 50mL of TE/10
0.5mL Tris HCl, pH 8.0
10 µL EDTA
49.5 mL $dH_2O$
Filter sterilize

1xBW buffer
5mM Tris-HCl pH8.0
0.5mM EDTA
1M NaCl
0.05% Tween-20
Filter sterilize

For 50mL of 1x BW Buffer
0.25 mL of 1M Tris pH 8.0
50 µL of 0.5M EDTA
10 mL of 5M NaCl
25 µL 100% Tween-20
Filter sterilize

2xBW buffer
10mM Tris-HCl pH 8.0
1mM EDTA
2M NaCl
Filter Sterilize

Vino Taste - 10 mL
0.500 g of Vino Taste powder
1mL 10X PBS
Bring up to 10 mL with $dH_2O$ *(make sure to dissolve the Vinotaste solid completely before adding rest of water!)*
Filter sterilize

1x Vogels, 1.5% sucrose (500mL)
10mL 50X Vogels
7.5g sucrose
490mL $dH_2O$
autoclave
