## Supplementary material for "Histone deacetylation and cytosine methylation compartmentalize heterochromatic regions in the genome organization of *Neurospora crassa*": Scadden_Supplementary-File_S3

**Klocko Lab ChIP protocol for Neurospora**

(modified from Selker lab ChIP protocol: Tamaru, H. et al., 2003; Nat Genet. _Updated 061721 by AK_)

Day 1:

1. Inoculate a 4mL liquid shaking overnight culture (typically 32ºC in Vogel's) from conidia (~$10^6$)
    a. A small sample of the strain collected from a baby slant with a pre-moistened stick will typically provide enough conidia for an overnight culture.
    b. Growth conditions can be varied; be sure to add nutrients that are required for the strain.
    c. Grow overnight for 16 to 18 hours at 32$^{o}$C

Day 2:

1. Collect mycelia by filtration and wash with 1x PBS
    a. (dilute 10x PBS 1:10 with distilled $H_2O$)
2. Add mycelia to 10mL of PBS in a 125mL flask
3. Add 250 µL of 37% formaldehyde straight out of bottle (final concentration at 1%) and shake gently (100rpm) for 15 minutes at room temp.
4. Quench the formaldehyde with 0.5mL 2.5M glycine
5. Collect mycelia by filtration through a Buchner funnel (with a vacuum)
6. Wash the crosslinked mycelia pellet with PBS on a Buchner funnel (with a vacuum)
7. Transfer the mycelial pellet to a 1.5mL Bioruptor (clear, polystyrene) tube containing 0.25mL ChIP lysis buffer
    a. or, if frozen Ogataea pellets, start here: resuspend them in 0.25mL ChIP Lysis buffer
8. Add in Halt protease inhibitor cocktail (1:100 dilution, so 2.5 µL for a typical 250 µL ChIP)
9. Lyse mycelia/chromatin in the Bioruptor (which uses sonication):
    a. 15-minute protocol (called "15min-30sec-ON" protocol) - 30 seconds on, 30 seconds off
10. Centrifuge samples for 5 minutes at 5500rpm (~3000 x g)
11. Transfer 250 µL (all) of the supernatant to clean 2.0 mL Screwtop tube
12. In a separate 2 mL screw-top tube, save 20 µL at -80 ºC for "input" sample
13. *Optional: preclear the lysate with protein A/G beads to remove any proteins that non-specifically "stick" to the beads. With histone mark ChIPs, AK has found this isn't required.*
    a. *Add 20 µL of protein A/G beads to 250 µL lysate and incubate at 4 ºC for 1-3 hours on a nutator*
    b. *microcentrifuge at 5000 rpm for 1 minute; transfer the pre-cleared supernatant into a new tube*
14. Add the desired antibody
    a. Use 2 µL antibody for each 250 µL of cell lysate (the supernatant from step 10 above)
    b. Can also use 300 µL lysate and 3 µL Antibody; note: for anti-FLAG, use 10 µL of anti-flag beads
15. Incubate overnight at 4 ºC on a rotator

Day 3:

1. Remove (20 µL X number of reactions, plus an extra 10 µL) from the protein A/G magnetic bead stock solution (stored at 4$^{o}$C) and move into a 1.7mL Eppendorf tube
2. Wash protein A/G magnetic beads 3x with ChIP Lysis buffer using the magnetic rack
    a. Remove from rack, add in 1mL ChIP Lysis buffer to resuspend beads, add tube back to rack and wait ~1 min until the beads move to the magnet side. Remove the 1mL supernatant. Repeat.
    b. Add (20 µL X number of reactions, plus an extra 10 µL) of ChIP lysis buffer to resuspend beads. This is the "stock solution" of beads you will use to add to the lysate + Ab samples
3. Add 20 µL of equilibrated protein A/G agarose beads to the cell lysate + antibody samples from the

previous night; parafilm tube tops.

4. Incubate for 2 hours at 4 ºC on the nutator to allow antibody binding to the protein A/G beads
5. Quickly centrifuge the tubes down to get all liquid/beads out of the tube caps.
6. Use a magnetic rack to collect the beads and discard supernatant
    a. Allow beads to move to magnet, and carefully pipette up and discard the supernatant
7. Add 1 ml of cold ChIP lysis buffer (without protease inhibitors) and incubate for 10 minutes at 4ºC on a rotator/nutator
8. Use a magnetic rack to collect the beads on the tube side, remove supernatant by going down the non-magnet side and discard
9. Repeat steps 6-8 (once more with ChIP lysis buffer)
10. Wash a third time (as in steps 6-8) with ChiP lysis buffer + 0.5M NaCl
11. Wash a fourth time (as in steps 6-8) with LiCl wash buffer
12. Wash a fifth time (as in steps 6-8) with TE
13. Add 62.5 µL of TES and incubate at 65 ºC for 10 minute to elute from beads.
    a. Mix tubes several times during incubation
    b. This step disrupts the Ab quaternary structure and release the crosslinked protein/DNA
14. Use a magnetic rack to collect the save supernatant in 2mL screw-top tubes
15. Repeat steps 13-14 with another 62.5 µL of TES to collect a second sample of the ChIP DNA into the same 2 mL screw top tube, for a total volume of 125 µL.
16. *Optional: if input samples are needed, get the saved 20 µL input sample from the -80$^o$C (from "Day 2, Step 12") and add 105 µL of TES into that 2 mL screw top tube with the input sample.*
17. De-crosslink by incubating all input or immunoprecipitation (IP) samples (125 µL total volume) for 6-16 hours in a 65 ºC incubator.

Day 4:

1. Add 125 µL of water and 2.5 µL 10mg/mL of RNAse A
2. Incubate for 2 hours at 50 ºC
3. Add 6.25 µL 20mg/mL Proteinase K and incubate for 2 hours at 50 ºC
4. Add 250 µL phenol/chloroform/IAA and extract aqueous layer
    a. (vortex for 30 seconds, microcentrifuge at 7.5k rpm for 5 min)
5. Add 250 µL chloroform and extract aqueous layer
    a. (vortex for 30 seconds, microcentrifuge at 7.5k rpm for 5 min)
6. Add 2 µL glycogen, 25 µL 3M Na-Acetate pH 6.0, 865 µL 100% EtOH (or 910.5 µL of 95% EtOH)
7. Quickly vortex the tubes, and then quickly spin them down in the centrifuge
8. precipitate overnight at -80 ºC.

Day 5:

1. Next morning, spin 10 min at 13k rpm to pellet the ChIP DNA; discard the supernatant
2. Slowly add 500 µL 70% EtOH, re-centrifuge the tube for 4 min, 13k rpm
3. Remove supernatant and discard (get the last 20 µL out with a P20 pipette)
4. dry down in speedvac (10 min at room temperature)
5. Resuspend the library pellet in 30 µL TE
6. Qubit HS the samples you obtained, to be sure there is some DNA present
    a. Anything above 0.1 ng/µL should be OK for sequencing
7. Can store samples at -20oC if you want to barcode the library later

Barcode the library:

**<u>NOTE: use this barcoding protocol below, using the NEBNext Ultra II kit from NEB.</u>**

<u>END PREP, in a 0.2mL PCR tube</u>

12.5µL ChIP DNA from step 4 above

(we are going to assume we have less than the 1µg total DNA maximum - crosscheck with Qubit HS data above)

37.5µL $dH_2O$

7µL NEBNext Ultra II End Repair reaction buffer (green top)

3µL NEBNext End Repair Enzyme Mix (green top)

Mix reaction well by pipetting

Use program CHIP20 on the PCR machine, with heated lid (99$^o$C)

CHIP20:

20$^o$C for 30 minutes

65$^o$C for 30 minutes

4$^o$C HOLD

<u>Ligate hairpin adapter</u>

To the 60µL end prep reaction (previous step), directly add these reagents:

30µL NEBNext Ultra II Ligation Master Mix (red top)

1µL NEBNext Ligation Enhancer (red top)

2.5µL NEBNext Adaptor for Illumina (red top)

Mix reaction well by pipetting

Use the program CHIP20-LIG

CHIP20-LIG: 20$^o$C hold, with NO heated lid

Ligate Adapters in the PCR machine for **<u>15 minutes</u>**!

Add 3µL of USER Enzyme (red top) to the ligation mixture, MIX WELL

Use program CHIP37

Incubate at 37$^o$C for 15 minutes (with heated lid at 99$^o$C)

(can store samples here at -20$^o$C, per the NEBNext protocol)

Ampure XP (Agencourt) purify the PCR reaction

Mix 93.5µL Ampure XP beads (warmed to RT) to the 93.5µL PCR reaction

(no need for extra PEG8000/NaCl; we want to remove fragments below 200bp, which are unligated adapters)

Bind DNA to beads for 15 minutes at RT

Put on magnetic rack for 5 min

(Ampure beads are magnetic and will go to the magnet side of the tube)

Remove and discard supernatant (DNA is on Ampure beads)

wash 2x with 200µL FRESH 80% EtOH

remove last of EtOH with P20 pipette

Dry 5 min at RT; take tube off magnetic rack

Add 15µL TE/10 to elute DNA, resuspend beads by pipetting (may need to use some force)

Keep tube at RT for 2 minutes on a NON-magnetic rack

Place tubes back on magnetic rack. wait 1 minute for beads to be drawn to one side

Save supernatant (15µL) with ChIP library to NEW 0.2mL PCR tubes (one tube per sample) for *immediate* use in the next PCR step.

Enrich DNA fragments by PCR

In the same 0.2mL PCR tube

15µL ChIP DNA with ligated adapter

(using all the reaction from above. Can just use half, if you want: 7.5µL library and 7.5µL $dH_2O$)

5µL Index Primer

(USE A DIFFERENT INDEX PRIMER PER REACTION; THIS ADDS THE UNIQUE BARCODE)

5µL Universal PCR primer

25µL NEBNext Ultra II Q5 Master Mix

Mix reaction, place into PCR machine, using CHIPAMP protocol:

RECORD ADAPTERS USED FOR EACH SAMPLE ON THIS PROTOCOL!

1 cycle 98°C 30 seconds

8 cycles of:

98°C, 10 sec

65°C, 75 sec **DO EIGHT PCR CYCLES TO MINIMIZE AT-RICH REGION DEPLETION!!!**

1 cycle 65°C, 5 minutes

hold at 4°C

Ampure XP (Agencourt) purify the PCR reaction

Mix 50µL Ampure XP beads (warmed to RT) and the 50µL PCR reaction

(no need for extra PEG8000/NaCl, as we want to get rid of fragments below 200bp, which are primer dimers)

Bind DNA to beads for 15 minutes at RT

Put on magnetic rack for 5 min

(Ampure beads are magnetic and will go to the magnet side of the tube)

Remove and discard supernatant (DNA is on Ampure beads)

wash 2x with 200µL FRESH 80% EtOH

remove last of EtOH with P20 pipette

Dry 5 min at RT

Take off magnetic rack

Add 25µL TE/10 to elute DNA, resuspend beads by pipetting (may need to use some force)

Keep tube at RT for 2 minutes on a NON-magnetic rack

Place tubes back on magnetic rack. wait 1 minute for beads to be drawn to one side

Save supernatant (25µL) with HiC library to NEW 1.7mL eppie tubes.

Store these barcoded libraries at -20 °C

Check and record concentration by Qubit HS.

Later (next day or following week):

Pool for seq

- For HT-seq at the UO core, sample should be 10nM in 50µL+. Use TE/10 for final volume.

- Single reads are fine: we just want to identify the genomic loci with these histone marks / purified proteins. Use the MiSeq at the UO core.

- Fragment analyzer data from the UO core should show that we have ~330 to 630bp band fragments (library will be 200-500 bp, and adapters add on ~130 bp)

NEBNext Barcodes

| Kit #7335L | Kit #7500L |
|---|---|
| Index 1: ATCACG | Index 13: AGTCAA |
| Index 2: CGATGT | Index 14: AGTTCC |
| Index 3: TTAGGC | Index 15: ATGTCA |
| Index 4: TGACCA | Index 16: CCGTCC |
| Index 5: ACAGTG | Index 18: GTCCGC |
| Index 6: GCCAAT | Index 19: GTGAAA |
| Index 7: CAGATC | Index 20: GTGGCC |
| Index 8: ACTTGA | Index 21: GTTTCG |
| Index 9: GATCAG | Index 22: CGTACG |
| Index 10: TAGCTT | Index 23: GAGTGG |
| Index 11: GGCTAC | Index 25: ACTGAT |
| Index 12: CTTGTA | Index 27: ATTCCT |

*Note: Sigma claims that deoxycholate (DOC) disrupts the interaction between the M2 FLAG antibody and the epitope; however, we have performed successful ChIP of FLAG tagged proteins using these buffers. One may wish to remove DOC for ChIP with FLAG tagged proteins.*
*AK has never used Doxycholate for his histone PTM ChIPs, and has removed that ingredient from the protocol above*

Recipes

ChIP Lysis Buffer (NO proteinase inhibitors, 200mL)

| | |
|---|---|
| 164mL | Water |
| 10mL | 1M HEPES, pH 7.5 stock |
| 5.6mL | 5M NaCl stock |
| 400 µL | 0.5M EDTA stock |
| 20mL | 10% Triton X-100 stock |
| *autoclave to sterilize* | |

ChIP Lysis Buffer + 0.5M NaCl (No protease inhibitors, 200mL)

| | |
|---|---|
| 149.6mL | Water |
| 10mL | 1M HEPES, pH 7.5 stock |
| 20mL | 5M NaCl stock |
| 400 µL | 0.5M EDTA stock |
| 20mL | 10% Triton X-100 stock |
| *autoclave to sterilize* | |

LiCl wash buffer (No protease inhibitors, 200mL)

| | |
|---|---|
| 177.6mL | Water |
| 2mL | 1M Tris-HCL pH 8.0 stock |
| 10mL | 5M LiCl stock |
| 10mL | 10% IGEPAL CA-630 (NP40) stock |
| 400 µL | 0.5M EDTA stock |
| *autoclave to sterilize* | |

TE (e.g., ChIP library wash buffer) (50mL)

| | |
|---|---|
| 49.4 mL | Water |
| 0.5 mL | 1M Tris-HCl, pH 8.0 stock |
| 0.1 mL | 0.5M EDTA stock |
| *Filter sterilize.* | |

TES (e.g., ChIP elution buffer) (50mL)

| | |
|---|---|
| 41.5mL | Water |
| 2.5mL | Tris-HCL pH 8 |
| 1.0mL | 0.5M EDTA |
| 5mL | 10% SDS |
| *Filter sterilize.* | |

TE/10 (e.g., ChIP library wash and elution buffer) (10mL)

| | |
|---|---|
| 9.9 mL | Water |
| 0.1 mL | 1M Tris-HCl, pH 8.0 stock |
| 2 µL | 0.5M EDTA stock |
| *Filter sterilize. MAKE FRESH EVERY TIME* | |
