## Supplementary material for "Histone deacetylation and cytosine methylation compartmentalize heterochromatic regions in the genome organization of *Neurospora crassa*": Scadden_Supplementary-Table_S1

**Supplemental Table S1.** Total and valid read numbers for individual replicates and merged data files for all Hi-C datasets presented in this manuscript.

| Strain Number | Relevant Genotype | Enzyme Used | Replicate | Sequenced Reads | Pairs Mapped | Hi-C Contacts | % Valid Reads |
|---|---|---|---|---|---|---|---|
| N3767 | Δ*cdp-2* | DpnII | 1 | 52052815 | 39844797 | 14931477 | 28.6852 |
| | | DpnII | 2 | 19762488 | 11102603 | 4835585 | 24.4685 |
| | | DpnII | 6 | 67416294 | 61941330 | 10841025 | 16.0807 |
| | | DpnII | Sample 1 | 23627005 | 20019345 | 4946733 | 20.9368 |
| | | | merged | 162858602 | 132908075 | 35449017 | 21.7667 |
| | | MseI | 1 | 36845367 | 29148159 | 9184936 | 24.9283 |
| | | MseI | 2 | 30992812 | 26100502 | 476356 | 15.3144 |
| | | | merged | 67838179 | 55248661 | 13897271 | 20.4859 |
| N3613 | Δ*chap* | DpnII | 1 | 17965310 | 14830891 | 4228813 | 23.5388 |
| | | DpnII | 2 | 18518179 | 15611982 | 4516955 | 24.3920 |
| | | | merged | 36483489 | 30442873 | 8730054 | 23.9288 |
| | | MseI | 1 | 17331784 | 15149664 | 2574138 | 14.8521 |
| | | MseI | 2 | 14508998 | 12724548 | 2195477 | 15.1318 |
| | | | merged | 31840782 | 27874212 | 4763900 | 14.9616 |
| N6144 | Δ*cdp-2*;Δ*dim-2* | DpnII | 1 | 30810072 | 26116220 | 10359637 | 33.6242 |
| | | DpnII | 2 | 30517329 | 25492915 | 12778286 | 41.8722 |
| | | | merged | 61327401 | 51609135 | 23100733 | 37.6679 |
